# Downstream metabolites of (+)-*cis*-12-oxo-phytodienoic acid function as noncanonical bioactive jasmonates in *Arabidopsis thaliana*

**DOI:** 10.1101/2024.05.01.592109

**Authors:** Rina Saito, Yuho Nishizato, Tsumugi Kitajima, Misuzu Nakayama, Yousuke Takaoka, Nobuki Kato, Minoru Ueda

**Affiliations:** Department of Molecular and Chemical Life Sciences, Graduate School of Life Sciences, Tohoku University, Sendai 980-8578, Japan; Department of Chemistry, Graduate School of Science, Tohoku University, Sendai 980-8578, Japan

## Abstract

(+)-*cis*-12-oxo-phytodienoic acid (*cis*-OPDA) is a biosynthetic precursor of the plant hormone (+)-7-*iso*-jasmonoyl-L-isoleucine (JA-Ile). It functions as an endogenous chemical signal independent of the JA-Ile receptor COI1-JAZ in *Arabidopsis thaliana*. The bioactive form of *cis*-OPDA that induces COI1-JAZ-independent gene expression remains unknown.

In this study, we hypothesized that the genuine bioactive forms of *cis*-OPDA are the downstream metabolites, which upregulate the expression of the OPDA marker genes such as *ZAT10*/*ERF5* in a JA-Ile-independent manner. These downstream metabolites function independently of the JA-Ile-COI1-JAZ-MYCs canonical jasmonate signaling module, and its electrophilic nature is essential for its bioactivity.

## Introduction

Jasmonic acid and related fatty acid-derived oxylipins are collectively referred to as jasmonates. Jasmonate, (+)-7-*iso*-jasmonoyl-L-isoleucine (JA-Ile; Fig. 1), is a lipid-derived plant hormone that regulates growth, reproduction, and defense responses against pathogens and chewing insects ^1-3^. In plant cells, JA-Ile is synthesized from α-linolenic acid via the biosynthetic precursor (+)-*cis*-12-oxo-phytodienoic acid (*cis*-OPDA) (Fig. 1A), which is reduced to OPC-8 by OPDA reductase 3 (OPR3), followed by oxidation to naturally occurring (+)-7-*iso*-jasmonic acid (JA, a *cis*-form) through three rounds of β-oxidation and conjugation with L-isoleucine (Ile) by GH3 enzyme jasmonic acid-resistant 1 (JAR1) or *At*GH3.10 to produce JA-Ile ^4-8^. Recently, an OPR3-independent route for JA-Ile biosynthesis from *cis*-OPDA was reported (Fig. 1B) ^9^. In this bypass route, three rounds of β-oxidation occurred from *cis*-OPDA to dinor-*cis*-OPDA (dn-*cis*-OPDA), tetranor-*cis*-OPDA (tn-*cis*-OPDA), and (+)-7-*iso*-4,5-didehydro JA (4,5-ddh-JA), which is then reduced to JA by OPR2 (Fig. 1B). An increase in JA-Ile levels in plant cells leads to the triggering of protein-protein interactions between the F-box protein CORONATINE INSENSITIVE 1 (COI1) and jasmonate-ZIM domain (JAZ) repressors, causing proteasomal degradation of JAZ to derepress transcription factors, such as MYC2, and activating gene expression ^10-13^.

**Fig. 1.**
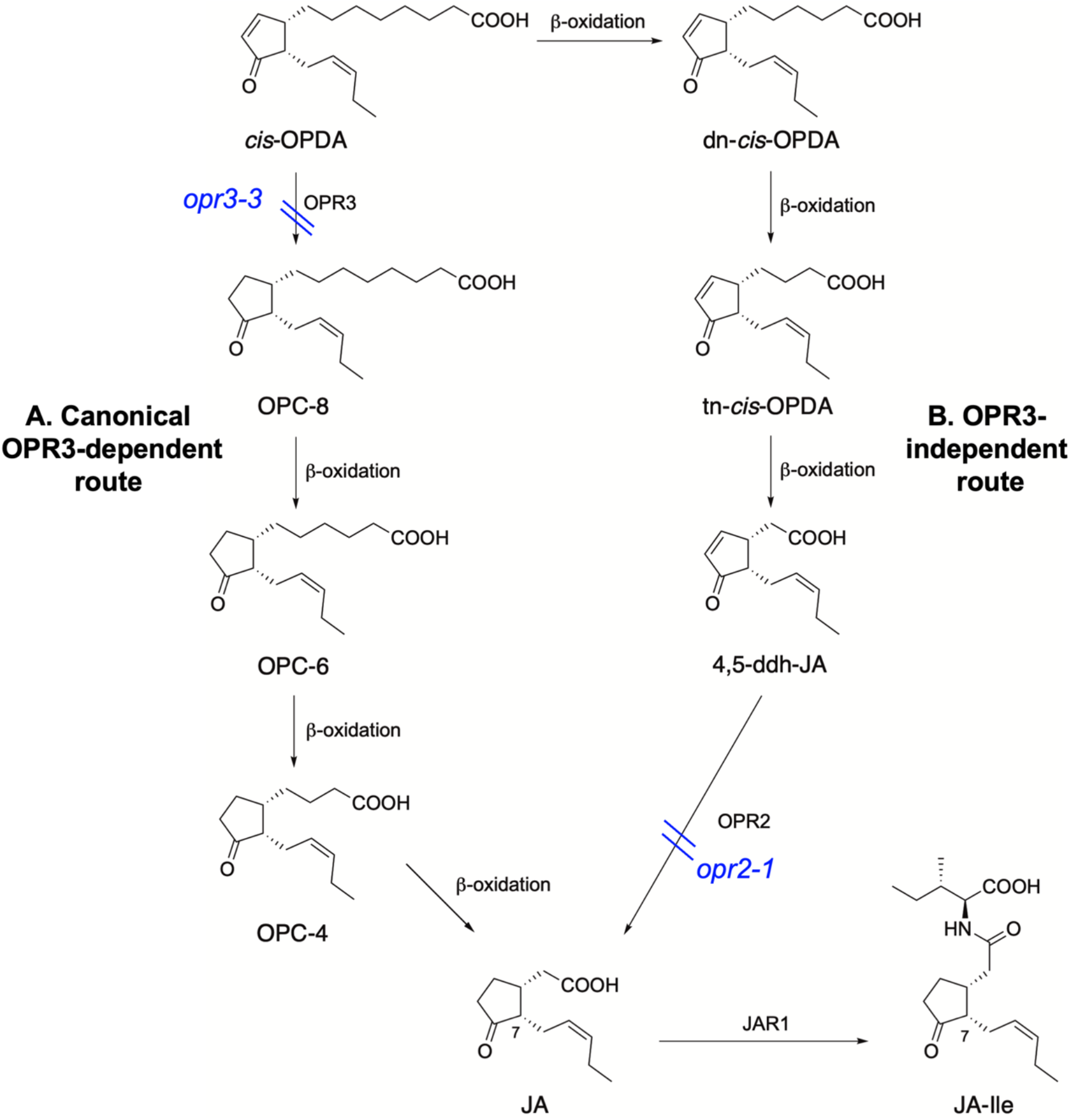
Biosynthetic pathway of JA-Ile *in planta*. **(A)** Canonical OPR3-dependent synthesis of JA-Ile. In this route, *cis*-OPDA is reduced to OPC-8 by OPR3, followed by oxidization to JA through three β-oxidation and conjugation with Ile by JAR1. **(B)** OPR3-independent synthesis of JA-Ile. Through three β-oxidation, *cis*-OPDA is oxidized to 4,5-ddh-JA, which is then reduced to JA by OPR2.

Several reports have demonstrated JA-Ile-independent biological activities of the biosynthetic intermediate *cis*-OPDA in *Arabidopsis thaliana*, tomato, and maize ^14-23^. Taki *et al*. demonstrated *cis-*OPDA-specific expression of genes, including genes encoding a transcription factor involved in salt-tolerance zinc finger (*ZAT10*), ethylene-responsive transcription factor (*ERF5)*, dehydration-responsive element binding protein (*DREB2A*), cold-responsive zinc finger (*ZAT12*), and iron deficiency-induced transcription factors (*FIT1*) by comparing the gene expression mediated by *cis*-OPDA and JA ^16^. *ZAT10* encodes a transcription factor (TF) involved in salt tolerance ^16^, and *ERF5* encodes an ethylene-responsive TF involved in wounding and cold stress responses ^24^. *DREB2A* encodes a TF involved in drought and salt stress ^25,26^. *ZAT12* encodes a TF in cold stress and salt tolerance ^27,28^. *FIT1* encodes a TF in iron acquisition ^29^. Biochemical, yeast two-hybrid (Y2H), and pull-down assays demonstrated that *cis*-OPDA is not recognized by *Arabidopsis* COI1-JAZ1/3/9 co-receptor pairs ^12,30^, implying that *cis*-OPDA functions via a COI1-independent mode of action (MOA). However, in previous studies on *cis*-OPDA in *A. thaliana*, their biological activities have been distinguished from those of JA-Ile using the *Arabidopsis* mutants *opr3-1* and *jar1-1*, in which the biosynthetic genes of JA-Ile (*OPR3* or *JAR1*) are impaired ^31^. This could be because *cis*-OPDA was not converted into JA-Ile in *opr3-1* and *jar1-1*. Nevertheless, recent studies have indicated that *cis*-OPDA can be converted into JA-Ile in *opr3-1* or *jar1-1* through OPR3-independent (Fig. 1B) ^9,32^ or JAR1-independent ^33^ bypass routes owing to the leaky nature of these mutant lines. As a result, the participation of *cis*-OPDA in JA-Ile-independent signaling pathways in *A. thaliana* remains debatable ^34,35^. To date, the COI1-independent MOA of *cis*-OPDA has been reliably reported only in the thermotolerance of *Marchantia polymorpha*, in which the electrophilic α,β-unsaturated cyclopentenone moiety of *cis*-OPDA plays an essential role ^36^.

In this paper, we investigated the function of *cis*-OPDA in *A. thaliana* and the non-leaky mutant lines *coi1-1* ^*37*^ and *opr2-1 opr3-3* ^9^ (Fig. 1) with impaired JA signaling and biosynthesis to identify the genuine bioactive form of *cis*-OPDA in *A. thaliana*.

## Results

### Analyses of *cis*-OPDA-induced gene expression in *A. thaliana* and the mutant lines on JA signaling

*cis*-OPDA (Fig. 1) was chemically synthesized using previously reported methods ^38,39^. The subsequent experiments were performed using synthetic *cis*-OPDA. Gene expression analysis was performed based on the information from a previously reported comprehensive DNA microarray analysis ^16^: the impact of (-)-JA (Fig. S1) or *cis*-OPDA was respectively evaluated on the expression of three JA marker genes, *OPR3, JAZ8*, and *MYC2* (Fig. 2A-C), and five *cis*-OPDA-specific marker genes, *ZAT10, ERF5, DREB2A, ZAT12*, and *FIT1* (Fig. 2D-H) ^16,31^.

**Fig. 2.**
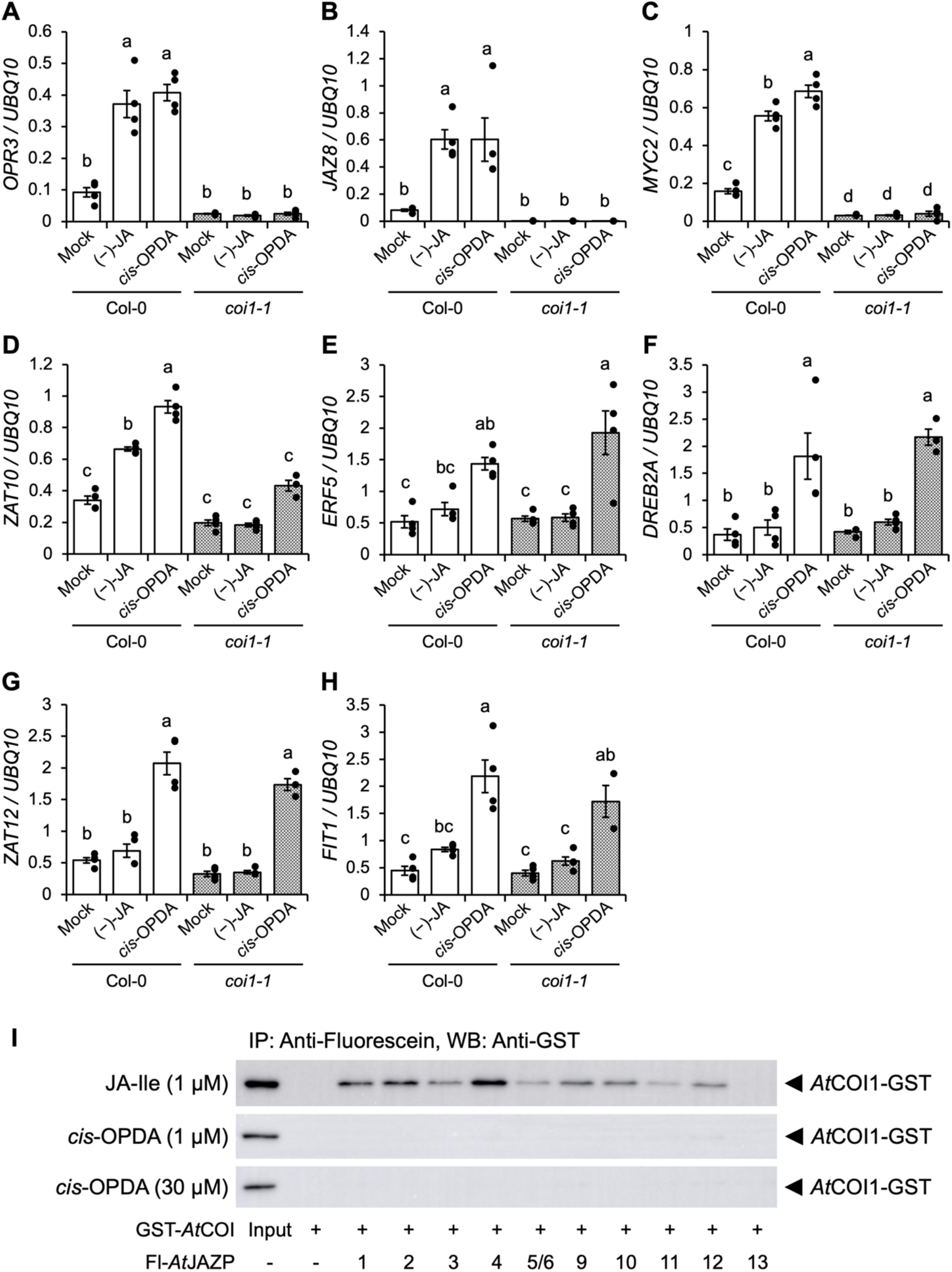
Validation of COI1-independence of *cis*-OPDA. (**A** to **H**) Gene expression analysis by RT-qPCR in 10-day-old WT (Col-0, white bar) and *coi1-1* mutant (gray bar) treated with 30 μM (-)-JA or *cis*-OPDA for 30 min and with no treatments (mock). The data are presented as mean ± SD (n = 3–4). Samples were normalized to the *UBQ10* level. Significant differences were evaluated by the ANOVA/Tukey Kramer test (*p* < 0.05). The experiments were repeated three times with similar results. (**I**) Evaluation of the affinity between *cis*-OPDA and COI1-JAZs. Pull-down assay of GST-*At*COI1 with Fl-*At*JAZPs in the presence of JA-Ile (1 μM) or *cis*-OPDA (1/30 μM). Fl-*At*JAZ13 was used as a negative control because JAZ13 has no canonical JAZ degron sequence, which is necessary for JA-Ile perception. GST-*At*COI1 bound to Fl-*At*JAZPs was pulled down with anti-fluorescein antibody and Protein G magnetic beads and analyzed by immunoblotting (anti-GST-HRP conjugate for detection of GST-*At*COI1). Arrowheads indicate GST-*At*COI1 (95 kDa). Experiments were repeated three times with similar results (shown in **Fig. S2 and S3**).

All compounds induced the expression of JA marker genes, *OPR3, JAZ8*, and *MYC2* in Col-0, and their expression was impaired in *coi1-1* (Fig. 2A-C). *cis*-OPDA induced the expression of *OPR3, JAZ8*, and *MYC2* in a COI1-dependent manner (Fig. 2A-C). In contrast, complex effects were observed regarding *ZAT10* expression, which was induced by all compounds in Col-0 (Fig. 2D). In the *coi1-1* mutant, *ZAT10* expression was not induced by (-)- JA, whereas moderate *ZAT10* expression was observed for *cis*-OPDA treatment (Fig. 2D). These findings imply that the expression of *ZAT10* by (-)-JA is COI1-dependent, whereas that of *cis*-OPDA depends on two distinct pathways, COI1-dependent and COI1-independent. Additionally, the expression of *ERF5, DREB2A, ZAT12*, and *FIT1* by *cis*-OPDA was not or slightly affected in *coi1-1*, confirming COI1-independence (Fig. 2E-H). In all experiments, *cis*-OPDA affected the expression of JA and OPDA marker genes. In addition, *cis*-OPDA showed no affinity for the functional COI1-JAZ1-6/9-12 co-receptor pairs in the pull-down assay (Fig. 2I and S2 and S3), confirming that *cis*-OPDA is not ligands for functional COI1-JAZ co-receptors.

We also examined the effect of *cis*-OPDA on the expression of JA and OPDA marker genes in the *myc2myc3myc4* triple mutant, in which the master TFs of JA signaling, MYC2/MYC3/MYC4, were impaired ^40^ (Fig. S4). The expression of the JA marker genes *OPR3* and *JAZ8* was suppressed in *myc2myc3myc4* (Figs. S4A and B). Different effects were observed for the five OPDA marker genes, *ZAT10, ERF5, DREB2A, ZAT12*, and *FIT1*; expressions of *ZAT10* and *DREB2A* were moderately suppressed, and that of *ERF5, ZAT12*, and *FIT1* were enhanced or not affected (Fig. S4C-G), respectively. The current results demonstrated that *cis*-OPDA mediated the expression of all OPDA marker genes independently of the canonical COI1-JAZ-MYC signaling pathway.

### *cis*-OPDA induces the expression of OPDA-marker genes independent of the conversion into JA-Ile

Given that *cis*-OPDA is a major biosynthetic precursor of JA-Ile, we examined whether *cis*-OPDA-induced gene expression depends on the *in planta* conversion into JA-Ile. This was investigated using the synthesized *cis*-OPDA-*d*_*5*_ (Fig. 3A and Scheme S1). The conversion of *cis*-OPDA into JA-Ile was examined using *Arabidopsis opr2-1opr3-3* double mutant with impaired conversion of *cis*-OPDA into JA-Ile ^9^. UPLC-MS/MS analysis demonstrated that *cis*-OPDA-*d*_*5*_ was converted into JA-*d*_*5*_-Ile within 30 min in Col-0, whereas this conversion was suppressed in *opr2-1opr3-3* (Fig. 3A). For the JA marker gene, *cis*-OPDA-induced expression of *JAZ8* in Col-0 was significantly suppressed in *opr2-1opr3-3* (Fig. 3B). In contrast, (-)-JA-mediated *JAZ8* expression in Col-0 cells was not affected by *opr2-1opr3-3* (Fig. 3B). Additionally, *cis*-OPDA-induced expression of *ZAT10* in Col-0 was not suppressed in *opr2-1opr3-3* (Fig. 3C). The expression of *ERF5, DREB2A, ZAT12*, and *FIT1* was not affected or suppressed in *opr2-1opr3-3* (Fig. 3D-3G). The undetectable level of JA-Ile in *opr2-1opr3-3* in Fig. 3A ^9^ implies that *cis*-OPDA induces the expression of OPDA maker genes independently of their conversion to JA-Ile.

**Fig. 3.**
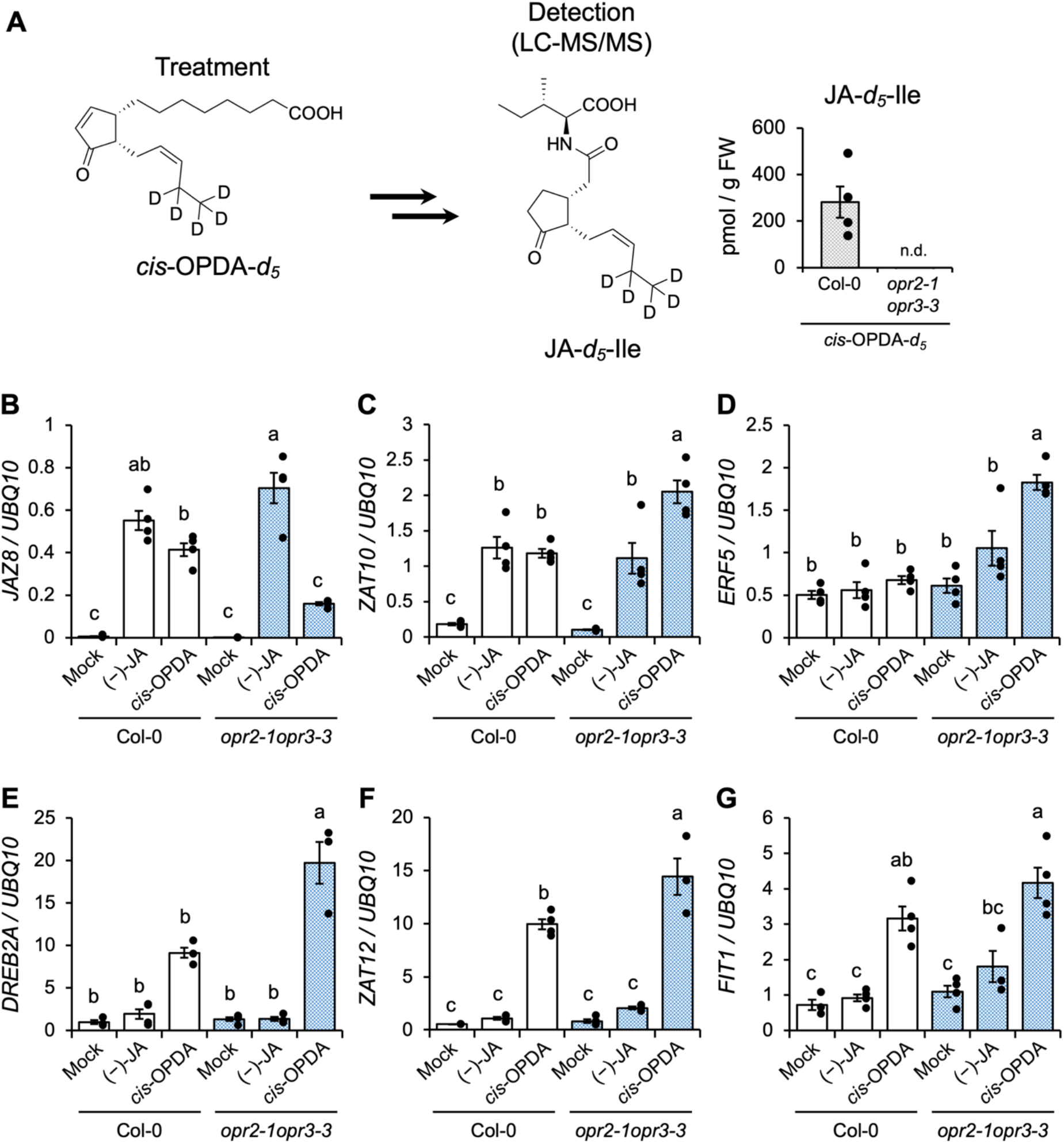
Gene expression analyses induced by *cis*-OPDA in *opr2-1opr3-3*. (**A**) Schematic diagram of the experimental design assessing the conversion of *cis*-OPDA to JA-Ile in Col-0 or *opr2-1opr3-3* by using deuterium-labeled precursors (*cis*-OPDA-*d*_*5*_); accumulation (pmol/fresh weight (g)) of JA-*d*_*5*_-Ile in 10-day-old WT (Col-0) and *opr2-1opr3-3* mutant treated with 30 μM *cis*-OPDA-*d*_*5*_ for 30 min. The data are presented as mean ± SD (n = 4). The experiments were repeated three times with similar results. (**B** to **G**) Gene expression analysis by RT-qPCR in 10-day-old WT (Col-0, white bar) and *opr2-1opr3-3* mutants (blue bar) treated with 30 μM (-)-JA or *cis*-OPDA for 30 min and with no treatments (mock). The data are presented as mean ± SD (n = 3–4). Samples were normalized to the *UBQ10* level. Significant differences were evaluated by the ANOVA/Tukey Kramer test (*p* < 0.05). The experiments were repeated three times with similar results. **Fig3D should be reexamined!**

### The *Arabidopsis cis*-OPDA transporter mutant and metabolism showed that *cis*-OPDA is not a genuine bioactive form

*cis*-OPDA is converted to JA in the peroxisomes, and the peroxisomal ATP-binding cassette (ABC) transporter COMATOSE (CTS) is involved in the import of *cis*-OPDA into the peroxisomes in *A. thaliana* (Fig. 4A, left) ^35^. Then, we examined whether *cis*-OPDA itself is a genuine bioactive form by using the *Arabidopsis cis*-OPDA transporter mutant. In the *cts1* mutant line, in which CTS is impaired, wound-induced accumulation of JA-Ile was significantly suppressed but not completely abolished ^41-44^. In the *cts1* mutant line, the level of *cis*-OPDA was significantly increased (Fig. 4A, right), however, to our surprise, *cis*-OPDA-induced expressions of *ZAT10, ERF5, DREB2A, ZAT12*, and *FIT1* were suppressed in *cts1* (Fig. 4B). Peroxisomal conversion of *cis-*OPDA into JA-Ile occurs through the canonical OPR3-dependent or OPR3-independent route via several downstream metabolites in *A. thaliana* (Fig. 1 and 3A). This finding indicated that the downstream metabolites, not *cis*-OPDA itself, are genuine bioactive forms.

**Fig. 4.**
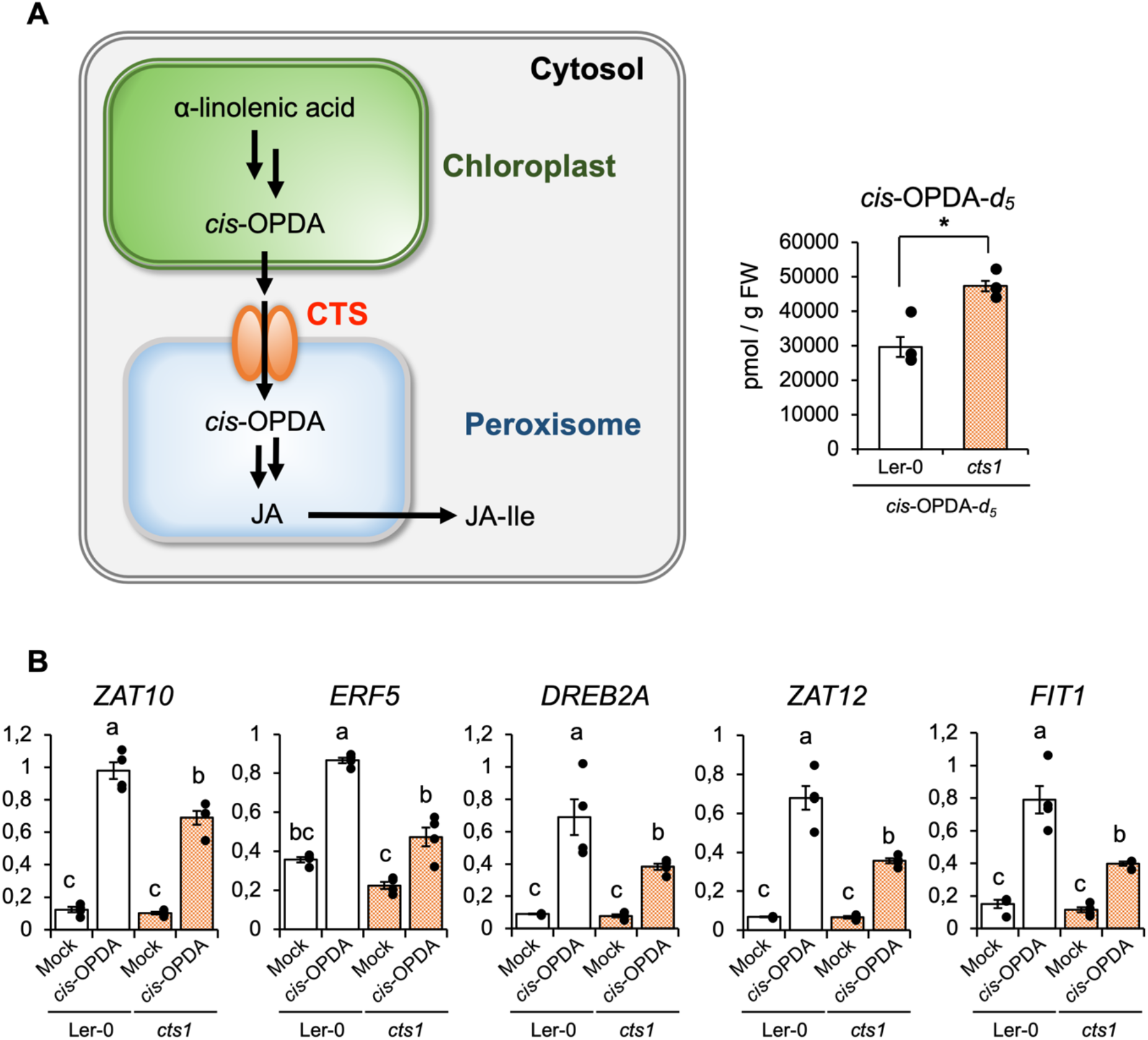
Downstream metabolites of *cis*-OPDA are the genuine bioactive form. (**A**) Left: Conversion of *cis*-OPDA into JA-Ile. The CTS transporter is localized on the peroxisomal membrane and involved in the import of *cis*-OPDA into the peroxisome. *cis*-OPDA is oxidized to dn-*cis*-OPDA, tn-*cis*-OPDA, and 4,5-ddh-JA in the peroxisome and finally converted to JA-Ile. Right: Accumulation (pmol/fresh weight (g)) of deuterium-labeled *cis*-OPDA-*d*_*5*_. 10-day-old *Arabidopsis* WT (Ler-0, white bar) and *cts1* mutant (orange bar) were treated with *cis*-OPDA-*d*_*5*_ (30 μM). The data are presented as mean ± SD (n = 4). Significant differences were evaluated by the Student’s t-test (\**p* < 0.05). (**B**) Gene expression analysis by RT-qPCR in 10-day-old WT (Ler-0, white bar) and *cts1* mutant (orange bar) with no treatments (mock) and 30 μM *cis*-OPDA treatment for 30 min. The data are presented as mean ± SD (n = 3–4). Samples were normalized to the *UBQ10* level. Significant differences were evaluated by the ANOVA/Tukey Kramer test (*p* < 0.05). The experiments were repeated three times with similar results.

### The downstream metabolites of *cis*-OPDA, dn-*cis*-OPDA, tn-*cis*-OPDA, and (+)-7-*iso*-4,5-didehydrojasmonic acid, are bioactive forms of *cis*-OPDA

*cis*-OPDA significantly upregulated the expressions of *ERF5, DREB2A, ZAT12*, and *FIT1* in *opr2-1opr3-3* compared to Col-0 (Fig. 3D-G). This finding implies that metabolites located downstream of *cis*-OPDA and upstream of JA in the OPR3-independent bypass route (Fig. 1B) are potential candidates for the bioactive form of *cis*-OPDA. As a result, we performed UPLC-MS/MS analyses of the downstream metabolites of *cis*-OPDA in the canonical OPR3-dependent (OPC-4, Fig. 1A) and OPR3-independent routes (dn-*cis*-OPDA, tn-*cis*-OPDA, and 4,5-ddh-JA; Fig. 1B) using Col-0 and *opr2-1opr3-3* (Fig. 5). For UPLC-MS/MS analysis, Col-0 and *opr2-1opr3-3* were treated with *cis*-OPDA-*d*_*5*_. Compared to Col-0, *cis*-OPDA-*d*_*5*_-treatment enhanced the accumulation of 4,5-ddh-JA-*d*_*5*_ and tn-*cis*-OPDA*-d*_*5*_ in the *opr2-1opr3-3* mutant but did not affect the level of dn-*cis*-OPDA*-d*_*5*_ (Fig. 5). Among these metabolites, the accumulation of tn-*cis*-OPDA*-d*_*5*_ (ca.4200 pmol/g FW in *opr2-1opr3-3* and ca. 2400 pmol/g FW in Col-0) and 4,5-ddh-JA*-d*_*5*_ (ca.3700 pmol/g FW in *opr2-1opr3-3* and ca. 1400 pmol/g FW in Col-0) was significantly higher than dn-*cis*-OPDA*-d*_*5*_ (ca.400 pmol/g FW). Notably, the accumulation of tn-*ci*s-OPDA*-d*_*5*_ and 4,5-ddh-JA*-d*_*5*_ was enhanced in *opr2-1opr3-3*. In contrast, the accumulation of OPC-4*-d*_*5*_, a downstream metabolite of the OPR3-dependent route, was below the detection limit in *opr2-1opr3-3* (Fig. 5).

**Fig. 5.**
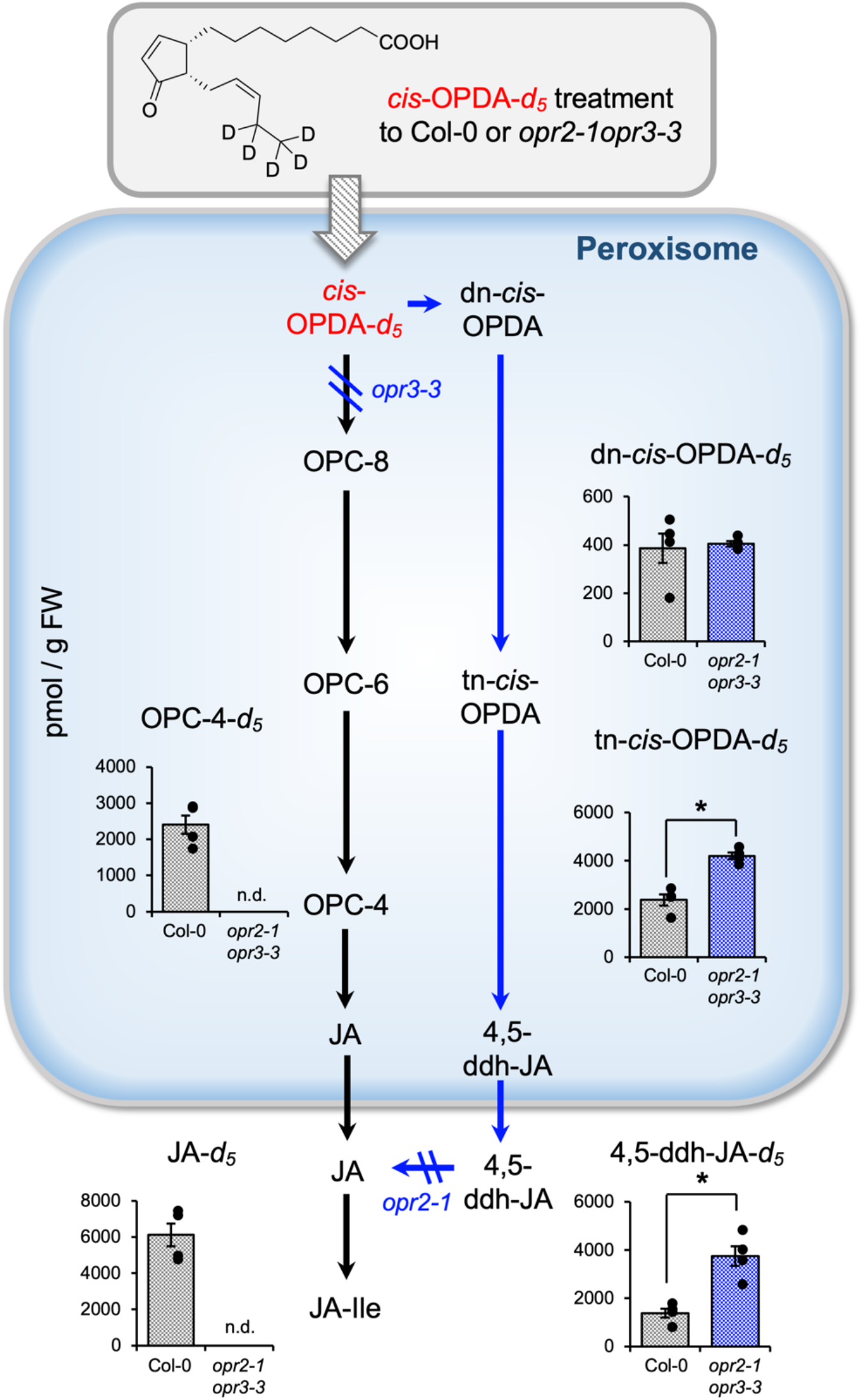
Conversion of *cis*-OPDA-*d*_*5*_ into jasmonate derivatives *in planta*. 10-day-old WT (Col-0) and *opr2-1opr3-3* were treated with *cis*-OPDA-*d*_*5*_ (30 μM) for 30 min. The data are presented as mean ± SD (n = 4). Significant differences were evaluated by Student’s t-test (\**p* < 0.05). The experiments were repeated three times with similar results.

### The downstream metabolites of *cis*-OPDA were similarly effective as *cis*-OPDA on the expression of OPDA-marker genes

We examined the effects of dn-*cis*-OPDA, tn-*cis*-OPDA, and 4,5-ddh-MeJA, the methyl ester of 4,5-ddh-JA (Fig. 6A and Scheme S2 and S3), on the expression of OPDA marker genes in *A. thaliana*. Tn*-cis*-OPDA and 4,5-ddh-MeJA were equally effective in upregulating expression of *ZAT10, ERF5, DREB2A, ZAT12*, and *FIT1* in the *opr2-1opr3-3* mutant (Fig. 6B, C, and S5A). Tn*-cis*-OPDA and 4,5-ddh-MeJA upregulated the expression of OPDA marker genes in a concentration-dependent manner (Fig. 6D and E).

**Fig. 6.**
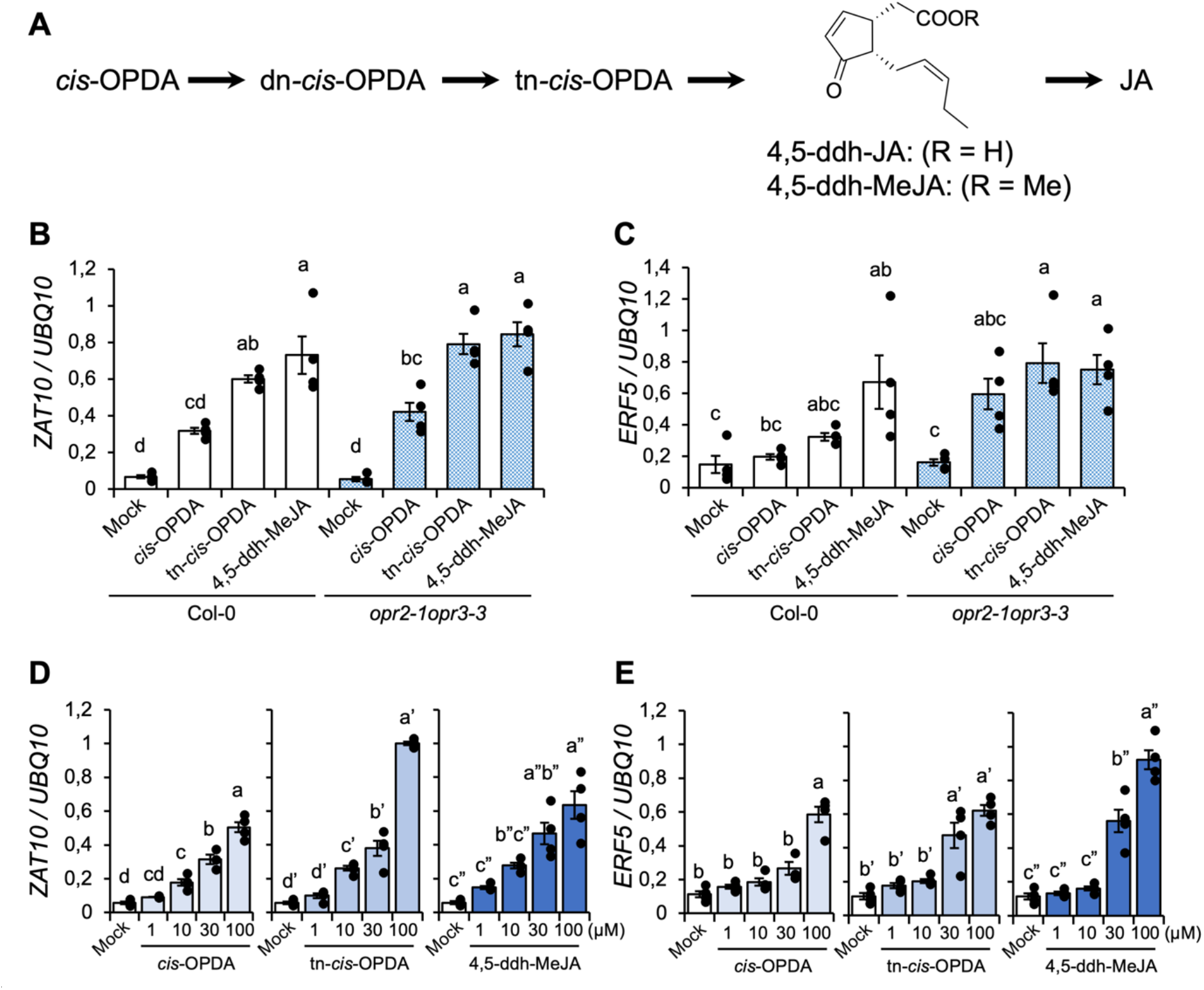
Gene expression mediated by *cis*-OPDA, tn-*cis*-OPDA and 4,5-ddh-MeJA in the mutant lines. (**A**) OPR3-independent bypassing route and chemical structure of 4,5-ddh-JA and 4,5-ddh-MeJA. (**B** and **C**) Gene expression analysis by RT-qPCR in 10-day-old WT (Col-0, white bar) and *opr2-1opr3-3* mutants (blue bar) treated with 30 μM *cis*-OPDA, tn-*cis*-OPDA, and 4,5-ddh-MeJA for 30 min. Mock indicates findings from WT and *opr2-1opr3-3* mutant without any treatments. (**D** and **E**) Gene expression analysis by RT-qPCR in 10-day-old *opr2-1opr3-3* mutants treated with 1, 10, 30, or 100 μM *cis*-OPDA (left), tn-*cis*-OPDA (middle), and 4,5-ddh-MeJA (right) for 30 min. Mock indicates findings from WT and *opr2-1opr3-3* mutants without any treatments. The data are presented as mean ± SD (n = 3–4). Samples were normalized to the *UBQ10* level. Significant differences were evaluated by ANOVA/Tukey Kramer test (*p* < 0.05). The experiments were repeated three times with similar results.

### The downstream metabolites of *cis*-OPDA caused gene expression through their electrophilic property

Monte *et al*. reported that *cis*-OPDA and dn-*cis*-OPDA upregulate the expression of *HSP* genes in *A. thaliana* and *M. polymorpha* in a COI1-independent manner ^36^. In addition, the upregulated gene expression depends on the electrophilic properties of *cis*-OPDA and dn-*cis*-OPDA because dn-*iso*-OPDA, which is an isomer of dn-*cis*-OPDA and less reactive as an electrophile, is significantly less effective than dn-*cis*-OPDA. Therefore, we compared the effects of tn-*cis*-OPDA and tn-*iso*-OPDA (Fig. 7A), 4,5-ddh-MeJA and 3,7-ddh-MeJA (Fig. 7A), corresponding to an *iso*-isomer of 4,5-ddh-MeJA, on the expression of *ZAT10, ERF5*, and *HSP* genes. The effect on the expression of *DREB2A, ZAT12*, and *FIT1* genes was also investigated (Fig. S6). As shown in Fig. 7 and S6, tn-*cis*-OPDA and 4,5-ddh-MeJA upregulated the expression of *ZAT10* (Fig. 7B), *ERF5* (Fig. 7C), *DREB2A* (Fig. S5A), *ZAT12* (Fig. S5B), and *FIT1* (Fig. S5C) whereas tn-*iso*-OPDA and 3,7-ddh-MeJA did not. Similarly, *cis*-OPDA, tn-*cis*-OPDA, and 4,5-ddh-MeJA upregulated the expression of *HSP*, whereas tn-*iso*-OPDA and 3,7-ddh-MeJA did not (Fig. 7D-F). This suggests that the electrophilic nature of tn-*cis*-OPDA and 4,5-ddh-JA might be responsible for upregulating *ZAT10, ERF5, DREB2A, ZAT12, FIT1*, and *HSP*s.

**Fig. 7.**
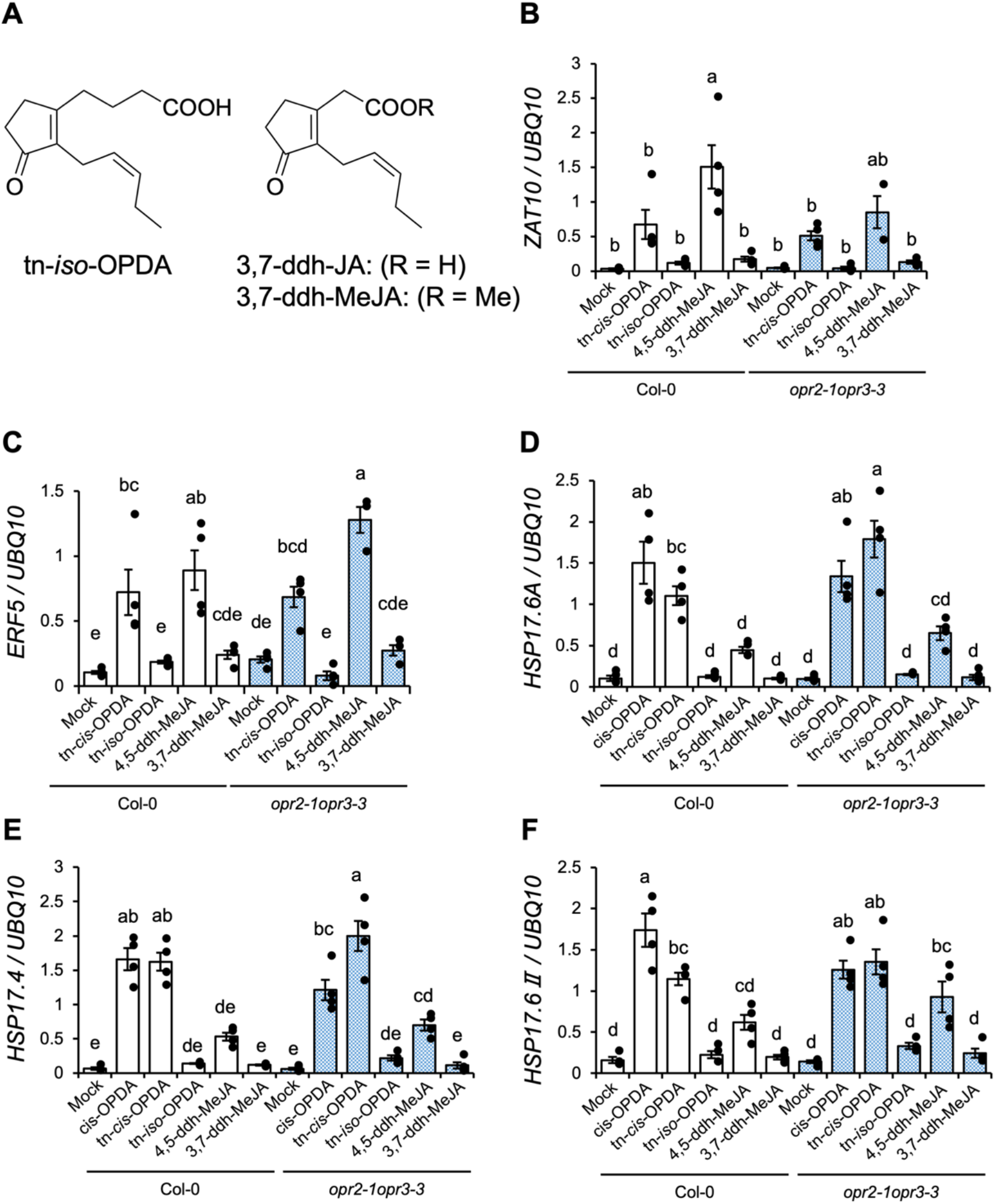
*cis*-OPDA, tn-*cis*-OPDA, and 4,5-ddh-MeJA mediated gene expression through their electrophilic properties. (**A**) Chemical structures of tn-*iso*-OPDA, 3,7-ddh-JA, and 3,7-ddh-MeJA. (**B** to **F**) Gene expression analysis by RT-qPCR in 10-day-old WT (Col-0, white bar) and *opr2-1opr3-3* mutant (blue bar) or without any treatments (mock) or treated with 30 μM tn-*cis*-OPDA, tn-*iso*-OPDA, 4,5-ddh-MeJA, and 3,7-ddh-MeJA for 30 min (**B, C**) or 30 μM *cis*-OPDA, tn-*cis*-OPDA, tn-*iso*-OPDA, 4,5-ddh-MeJA, and 3,7-ddh-MeJA, for 180 min (**D, E, F**). The data are presented as mean ± SD (n = 3–4). Samples were normalized to the *UBQ10* level. Significant differences were evaluated by the ANOVA/Tukey Kramer test (*p* < 0.05). The experiments were repeated three times with similar results.

## Discussion

Previous studies reported that *cis*-OPDA is a bioactive jasmonate in *A. thaliana* ^14-21^. Here, we reexamined its bioactivities by combining bioassays using the complete loss-of-function *Arabidopsis* mutant lines, *coi1-1*, and *opr2-1opr3-3*, and *in vitro* biochemical assays to assess the affinity between *cis*-OPDA and COI1-JAZs, and concluded that *cis*-OPDA is not the genuine bioactive form responsible for the expression of OPDA-marker genes. Our findings revealed that the downstream metabolites of *cis*-OPDA in the OPR3-independent biosynthetic route of JA-Ile are responsible for *cis*-OPDA-induced expression of *ZAT10* and *ERF5* (Fig. 8).

**Fig. 8.**
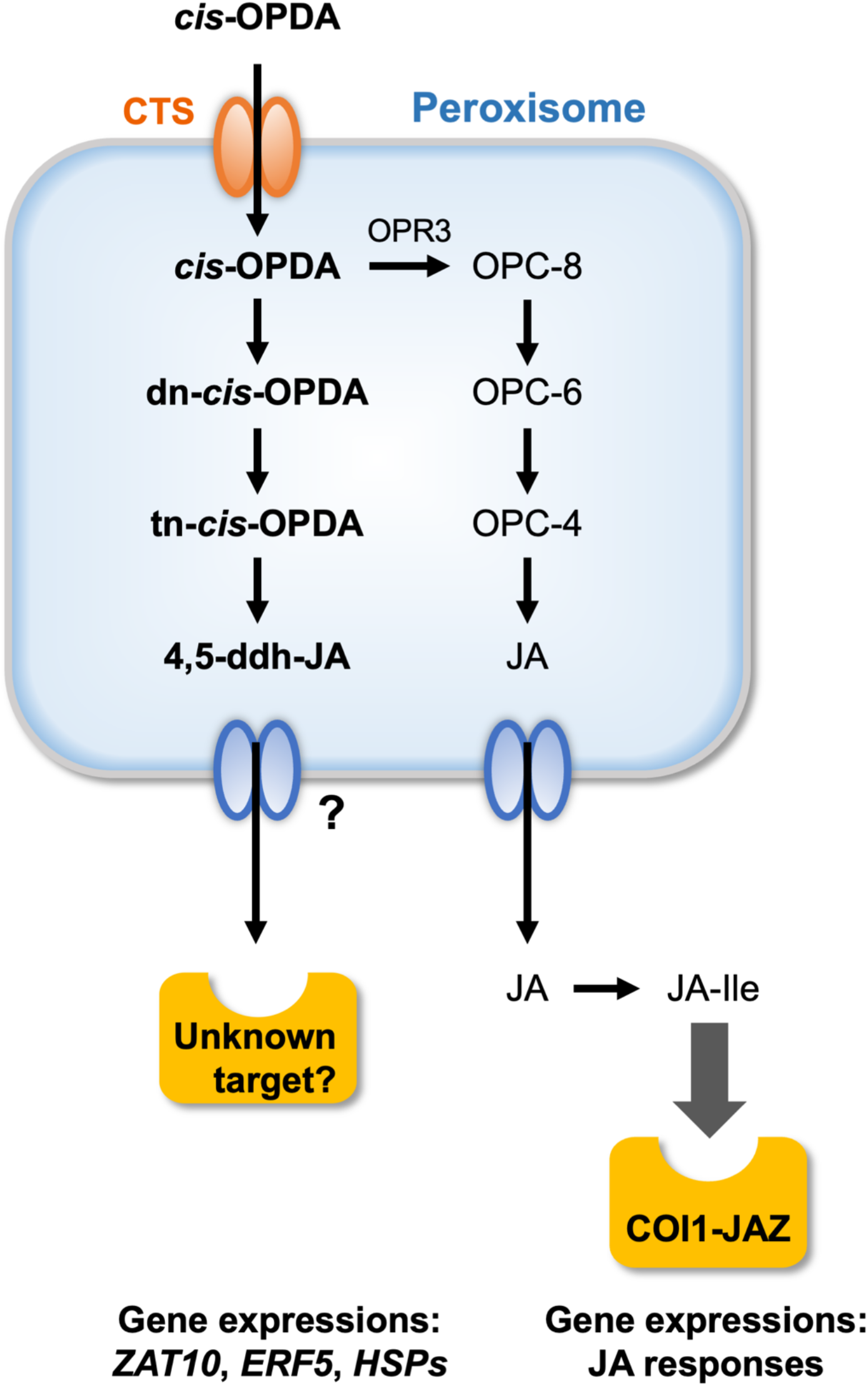
A model demonstrating that downstream metabolites of *cis*-OPDA function as bioactive jasmonates independent of canonical JA signaling. According to the hypothesis, downstream metabolites of *cis*-OPDA are responsible for the expression of OPDA marker genes such as *ZAT10*/*ERF5* through the unknown target.

The finding that the expression of OPDA marker genes was suppressed in the *cts1* mutant lines ^43^, in which *cis*-OPDA transport to peroxisomes was inhibited, strongly supports our conclusion (Fig. 4). This finding demonstrated that the downstream metabolites of *cis*-OPDA, and not *cis*-OPDA itself, were genuine bioactive forms. Next, we focused on the fact that *cis*-OPDA caused higher expression of *ERF5, DREB2A, ZAT12*, and *FIT1* in *opr2-1opr3-3* mutant compared to Col-0 (Fig. 3D-G). This result indicates that common metabolites of *cis*-OPDA are responsible for the upregulation of these genes in *opr2-1opr3-3*. Chini *et al*. reported that the OPR3-independent JA biosynthetic pathway operates weakly in Col-0 and is predominant in *opr2-1opr3-3*/*opr3-3* mutants ^9^. In *opr2-1opr3-3*, the downstream metabolites of the OPR3-independent route, tn-*cis*-OPDA and 4,5-ddh-JA, accumulated more than in Col-0, whereas little accumulation was observed for the downstream metabolites of *cis*-OPDA in the canonical OPR3-dependent route (Fig. 5). The accumulation of metabolites in *opr3-3* was previously reported using deuterium-labeled α-linolenic acid ^9^; however, we used deuterium-labeled *cis*-OPDA-*d*_*5*_ to exclude the effect of metabolites from the branched metabolic routes of α-linolenic acid, such as hydroperoxide lyase/isomerase, epoxyalcohol synthase, divinyl ether synthase, and 9-LOX routes ^4,45^. As a result, downstream metabolites of *cis*-OPDA in the OPR3-independent route are expected to be bioactive forms of *cis*-OPDA. Unfortunately, we could not identify one of them or all of them function as endogenous chemical signals. Identification of the genuine endogenous signal must await the target identification and subsequent evaluation of ligand-receptor binding affinity. Considering that UPLC-MS/MS analysis revealed that the accumulation of tn-*cis*-OPDA and 4,5-ddh-JA was increased in *opr2-1opr3-3* compared to Col-0, while the accumulation of dn-*cis*-OPDA was unchanged (Fig. 5), suggesting that tn-*cis*-OPDA and 4,5-ddh-JA are plausible endogenous signaling molecule (Fig. 8). However, our current data cannot exclude the possible function of dn-*cis*-OPDA as an endogenous chemical signal. Dn-*cis*-OPDA is converted into tn-*cis*-OPDA and then 4,5-ddh-JA through β-oxidation in the peroxisome and this conversion cannot be impaired because of the lack of a mutant line (Fig. 1). But, 4,5-ddh-JA is the most downstream metabolite produced in peroxisome, and previous report have shown that 4,5-ddh-JA is reduced to JA by cytosol-localized OPR2 (Figure 8).^9^ Thus, 4,5-ddh-JA is a promising candidate as an endogenous signaling molecule.

Our findings also indicated that the electrophilic reactivity of 4,5-ddh-JA and tn-*cis*-OPDA may be responsible for the expression of OPDA marker genes *ZAT10* and *ERF5*. Monte *et al*. demonstrated that *cis*-OPDA and dn-*cis*-OPDA upregulate *Mp*COI1-independent expression of *HSP* genes in *M. polymorpha* through electrophilic reactivities ^36^. They concluded that electrophilic reactivity is essential for *HSP* expression because it is not induced by dn-*iso*-OPDA, an isomer of dn-*cis*-OPDA with significantly lower electrophilic reactivity, due to the tetra-substituted α,β-unsaturated ketone moiety ^36,46^. Similarly, tn-*iso*-OPDA and 3,7-ddh-JA, *iso*-isomers of tn-*cis*-OPDA and 4,5-ddh-JA with little electrophilic reactivity, did not induce the expression of OPDA maker genes (Fig. 7 and S6).

Downstream metabolites of *cis*-OPDA function as chemical signals that cause the upregulation of OPDA marker genes independently of COI1 and MYCs (Fig. 2 and S4). COI1-independent gene upregulation may be attributed to non-COI1-targets of *cis*-OPDA, such as cyclophilin 20-3 and glutathione S-transferase 19, which are proposed putative targets of *cis*-OPDA (Fig. 8) ^18,47^. Further studies identifying the non-COI1-target of *cis-*OPDA will enable detailed genetic studies of the unknown MOA of *cis-*OPDA.

In this study, we examined the effects of *cis*-OPDA on *coi1-1* and *opr2-1opr3-3* mutants in which JA signaling and biosynthesis were impaired. In addition, we demonstrated that *cis*-OPDA had no affinity for any of the functional COI1-JAZ co-receptor pairs. Combining the results of gene expression and UPLC-MS/MS analyses, downstream metabolites of *cis*-OPDA are responsible for the expression of OPDA marker genes through a non-COI1 target.

## Materials and Methods

### Plant Materials

*A. thaliana* ecotype Col-0 and Ler-0 seeds were surface-sterilized in 5% sodium hypochlorite with 0.3% Tween 20 and vernalized for 2–3 d at 4 °C. All seedlings were grown (118 μmol m^-2^ s^-1^) under a 16 h light/8 h dark cycle at 22 °C in a CL-301 growth chamber (Tomy Seiko Co., Ltd., Japan). Seedlings were grown in 1/2 Murashige and Skoog (MS) liquid medium. The mutant lines used in this study were *coi1-1* ^*37*^, *opr2-1opr3-3* ^*9*^, *myc2myc3myc4* ^*40*^, and *cts1* ^*41*^. The seeds of *coi1-1* and *opr2-1opr3-3* mutants were kindly gifted from Professor Roberto Solano and Dr. Andrea Chini (CNB-CSIC, Spain). The *cts1* mutant was purchased from the Nottingham *Arabidopsis* Stock Centre (NASC). To select homozygous *coi1-1*, heterozygous *coi1-1* seeds were geminated on a 1/2 MS plate containing 10 μM JA, and well-grown 4 d seedlings (homozygous *coi1-1*) were transferred to a 1/2 MS liquid medium. After 6 d of culture, the seedling was treated with each compound ((-)-JA, *cis*-OPDA, *cis*-OPDA-*d*_*5*_, dn-*cis*-OPDA, tn-*cis*-OPDA, 4,5-ddh-MeJA, tn-*iso*-OPDA, 3,7-ddh-MeJA). For *cts1*, surface-sterilized seeds were pipetted onto 1/2 MS plate medium, and seed coats were disrupted using sterile tweezers to stimulate germination ^42^.

### Chemical synthesis

#### Synthesis of (+)-*cis*-OPDA-*d*_*5*_

(+)-12-(*R*)-Hydroxy-phytodienoic alcohol-*d*_*5*_ was synthesized as described in the Supplementary Text. To a solution of the diol (+)-12-(*R*)-hydroxyphytodienoic alcohol-*d*_*5*_ (32.7 mg, 0.11 mmol) in acetone (9.8 mL), Jones reagent (4.0 M solution) at −20 °C was added until the orange color of the reagent persisted (8 drops). After 10 min of stirring at −20 °C, *i*-PrOH was added to quench the remaining reagent. AcOEt/*n*-hexane (1:1) and H_2_O were then added, and the water layer was extracted with AcOEt. The combined organic layers were washed with saturated aqueous NaCl, dried over Na_2_SO_4,_ and concentrated under reduced pressures. The residue was purified by medium-pressure chromatography (Isolera, eluent: 0.1:88:12 AcOH/*n*-hexane/EtOAc to 0.1:99.9 AcOH/EtOAc) to obtain *cis*-OPDA-*d*_*5*_ (14.4 mg, 42%) as a colorless oil. [α]d^23^ +127.2 (*c* 0.34, CHCl_3_). ^1^H NMR (400 MHz, CDCl_3_) δ_H_: 7.74 (dd, *J* = 6.0, 2.8 Hz, 1H), 6.19 (dd, *J* = 6.0, 2.0 Hz, 1H), 5.32–5.45 (m, 2H), 2.93–3.02 (m, 1H), 2.40–2.56 (m, 2H), 2.36 (t, *J* =7.8 Hz, 2H), 2.08–2.19 (m, 1H), 1.09–1.82 (m, 12H); ^13^C NMR (100 MHz, CDCl_3_) δ_C_: 211.1, 179.7, 167.3, 132.9, 132.5, 127.0, 49.8, 44.3, 34.0, 30.7, 29.6, 29.1, 28.9, 27.6, 24.6, 23.8, 19.3–20.3 (m), 12.4–13.3 (m); IR (neat) cm^−1^: 2929, 2222, 1707, 1213; HRMS (ESI, negative) *m/z* [M-H]^-^ calcd for C_18_H_22_D_5_O_3_: 296.2279, found: 296.2268.

### Synthesis of tn-*cis*-OPDA

(+)-8-(*R*)-hydroxyphytodienoyl alcohol was synthesized as described in the Supplementary Text. To a solution of (+)-8-(*R*)-hydroxyphytodienoyl alcohol (18.0 mg, 81.1 μmol) in acetone (7 mL), Jones reagent (4.0 M solution) was added at −20 °C dropwise until the color of the reagent persisted (7 drops). *i*-PrOH and AcOEt/*n*-hexane (1:1) were then added to quench the remaining reagents. The mixture was extracted using EtOAc. The organic layer was washed with saturated aqueous NaCl, dried over Na_2_SO_4_, and then filtered. The reaction mixture was purified using medium-pressure chromatography (Isolera, eluent: 88:12:0.1 n-hexane/EtOAc/AcOH to 100:0.1 EtOAc/AcOH) to obtain tn-*cis*-OPDA as a colorless oil (18.5 mg, 96%). [α]d^23^ +207.2 (*c* 0.14, CHCl_3_); ^1^H NMR (400 MHz, CDCl_3_) δ_H_: 7.75 (dd, *J* = 5.8, 2.8 Hz, 1H), 6.21 (dd, *J* = 5.8, 1.8 Hz, 1H), 5.52-5.27 (m, 2H), 3.09–2.92 (m, 1H), 2.65–2.30 (m, 4H), 2.27–1.93 (m, 3H), 1.89–1.59 (m, 3H), 1.34–1.11 (m, 1H), 0.97 (t, *J* = 7.6 Hz, 3H); ^13^C NMR (100 MHz, CDCl_3_) δ_C_: 210.56, 178.52, 166.28, 133.21, 132.91, 126.71, 49.67, 44.04, 33.96, 30.19, 23.79, 22.76, 20.80, 14.01; IR (neat) cm^−1^: 3050, 1732, 1702, 1109; HRMS (ESI negative) *m/z* [M-H]^-^ calcd for C_14_H_19_O_3_: 235.1340, found: 235.1328.

### Synthesis of 4,5-ddh-MeJA

(+)-Methyl 4,5-ddh-6-*epi*-cucurbate was synthesized as described in the Supplementary Text. To a solution of (+)-methyl 4,5-ddh-6-*epi*-cucurbate (9.8 mg, 43.7 μmol) in acetone (4.5 mL), Jones reagent (4.0 M solution) was added at −20 °C until the orange color of the reagent persisted (16 drops). After 10 min of stirring at −20 °C, *i*-PrOH was added to quench the remaining reagent. Subsequently, *n*-hexane and H_2_O were added, and the water layer was extracted with *n*-hexane. The combined organic layers were washed with saturated aqueous NaCl, dried over Na_2_SO_4,_ and concentrated under reduced pressures. The residue was purified by medium-pressure chromatography (Isolera, eluent: 98:2 *n*-hexane/EtOAc to 80:20 *n*-hexane/EtOAc) to obtain 4,5-ddh-*cis*-MeJA (5.3 mg, 55%) as a colorless oil. Diastereomeric purity of 4,5-ddh-*cis*-MeJA was > 99% by ^1^H NMR spectroscopy (δ_H_ =7.71 (dd, *J* = 5.7, 2.7 Hz, 1H) for 4,5-ddh-*cis*-MeJA; 7.63 (dd, *J* = 5.7, 2.4 Hz, 1 H) for the *trans* isomer). [α]d^24^ +127.4 (*c* 0.26, CHCl_3_). ^1^H NMR (400 MHz, CDCl_3_) δ_H_: 7.71 (dd, *J* = 5.7, 2.7 Hz, 1H), 6.22 (dd, *J* = 5.7, 2.0 Hz, 1H), 5.45 (dtt, *J* = 10.9, 7.4, 1.8 Hz, 1H), 5.32 (dddt, *J* = 10.9, 6.8, 5.0, 1.4 Hz, 1H), 3.72 (s, 3H), 3.55-3.45 (m, 1H), 2.76 (dd, *J* = 16.4, 5.8 Hz, 1H), 2.57 (dt, *J* = 6.8, 5.0 Hz, 1H), 2.53 (brdt, *J* = 15.7, 5.0 Hz, 1H), 2.21 (dd, *J* = 16.4, 10.1 Hz, 1H), 2.13 (brdt, *J* = 15.7, 6.8 Hz, 1H), 2.05 (quintet, *J* = 7.4 Hz, 2H), 0.97 (t, *J* = 7.4 Hz, 3H); ^13^C NMR (100 MHz, CDCl_3_) δ_C_: 210.03, 172.34, 165.87, 133.66, 133.19, 126.17, 51.87, 48.32, 40.41, 34.44, 24.39, 20.79, 13.85; IR (film) cm^−1^:2961, 1733, 1717, 1198; HRMS (ESI, positive) *m/z* [M+Na]^+^ calcd for C_13_H_18_NaO_3_: 245.1154, found: 245.1148.

### Gene expression analyses

Surface sterilized *A. thaliana* seeds were germinated (118 μmol m^-2^ s^-1^) under a 16 h light/8 h dark cycle at 22 °C in a CL-301 growth chamber (TOMY SEIKO Co., Ltd., Japan) after vernalization in the dark at 4 °C for 2 d. 10-day-old seedlings (5–6 seedlings/sample) grown in 1/2 MS liquid medium were treated with each compound ((-)-JA, *cis*-OPDA, dn-*cis*-OPDA, tn-*cis*-OPDA, 4,5-ddh-MeJA, tn-*iso*-OPDA, 3,7-ddh-MeJA). Based on the results of previously reported comprehensive DNA microarray analyses ^16^, we performed gene expression analyses by adding *cis*-OPDA to *Arabidopsis* WT (Col-0) and the *coi1-1* mutant. *Arabidopsis* WT and *coi1-1* plants were treated with (-)-JA (a mixture of *cis* and trans = 5/95; fig. S1) or *cis*-OPDA, to examine their effects on the expression of three JA marker genes, *OPR3, JAZ8*, and *MYC2*, as well as *cis*-OPDA-specific marker genes, *ZAT10, ERF5, DREB2A, ZAT12*, and *FIT1* ^16,31^. Each sample was frozen after treatment for 30 min, and total RNA was extracted using ISOGEN (NIPPON GENE, Japan). First-strand cDNA was obtained using ReverTra Ace^®^ reverse transcriptase (TOYOBO, Japan) with oligo-dT primers. A StepOnePlus Real-Time PCR System (Life Technologies, USA) was used for quantitative PCR (qPCR). PCR conditions were followed: an initial hold at 95 °C for 30 s, followed by a two-step PCR program of 95 °C for 5 s and 60 °C for 30 s for 40 cycles. Primer sequences used in this study are listed in Supplementary Table S1. *Polyubiquitin 10* was used as the reference gene.

### UPLC-MS/MS analysis of *cis*-OPDA metabolites

Surface sterilized *A. thaliana* seeds were germinated (118 μmol m^-2^ s^-1^) under a 16 h light/8 h dark cycle at 22 °C in a CL-301 growth chamber (TOMY SEIKO Co., Ltd., Japan) after vernalization in the dark at 4 °C for 2 d. 10-day-old seedlings (5 seedlings/sample) grown on 1/2 MS liquid medium were treated with *cis*-OPDA-*d*_*5*_ (30 μM). The frozen plant materials were homogenized and extracted using 600 μL of ice-cold (pre-cooled at −20 °C) 25% MeOH/H_2_O. The samples were sonicated for 10 min and extracted for 30 min at 4 °C using a rotator. After centrifugation at 15,000 g for 15 min, supernatants were collected. The pooled supernatants were purified by one-step reversed-phase polymer-based solid-phase extraction using Oasis^®^ HLB cartridges (Waters, USA). Before sample loading, the SPE sorbent was conditioned with 1 mL of 100% MeOH and equilibrated with 1 mL of 0.1% HCOOH/H_2_O (v/v). After the samples had been loaded onto the Oasis^®^ HLB column, interfering compounds were removed by washing with 1 mL of 25% MeOH/H_2_O, following which the pre-concentrated analytes on the sorbent were eluted with 2 mL of 50% MeOH/H_2_O and 2 mL 100% MeOH. The eluent solution was evaporated to dryness at 30 °C under reduced pressure through freeze-drying. The samples were then resuspended in 100 μL of 100% MeOH.

The samples were examined by UPLC-MS/MS using a TripleTOF 5600 system (AB SCIEX, USA) operating in the negative mode. Liquid chromatography separation was performed with an Eclipse Plus C18 RRHD 1.8 μm (Φ1.05 × 50 mm, Waters, USA) at 35 °C and a flow rate of 0.3 mL/min. The elution was performed using a gradient of water (solvent A) and MeOH (solvent B), both containing 0.1% formic acid (v/v). The proportion of solvent B in the eluent was increased linearly from 10% to 60% for 2 min and from 60% to 90% for 8 min of the elution phase, followed by maintaining a flow of 90% B for 1 min. The column was re-equilibrated with 10% solvent B for 3 min.

### Pull-down assay

For the pull-down experiments using fluorescein-tagged JAZ peptides (Fl-*At*JAZPs), purified GST-*At*COI1 (5 nM), Fl-*At*JAZP (10 nM), and each compound (JA-Ile or *cis*-OPDA) in 350 μL of incubation buffer (50 mM Tris-HCl buffer, pH 7.8, 100 mM NaCl, 10% glycerol, 0.1% Tween20, 100 nM inositol-1,2,4,5,6-pentakisphosphate (IP5)) were combined with anti-fluorescein antibody (0.2 μL, GeneTex, GTX26644, USA), and incubated for 10–15 h at 4 °C with rotation. After incubation, the samples were combined with SureBeads™ Protein G (10 μL in 50% incubation buffer slurry; Bio-Rad, Hercules, CA, USA). After 3 h of incubation at 4 °C with rotation, the samples were washed three times with 350 μL of wash buffer (phosphate-buffered saline containing 0.1% Tween 20). The washed beads were resuspended in 35 μL of SDS-PAGE loading buffer containing dithiothreitol (DTT, 100 mM). After heating for 10 min at 60 °C, the samples were subjected to SDS-PAGE and analyzed by western blotting. The bound GST-COI1 protein was detected using an anti-GST HRP conjugate (RPN1236, GE Healthcare, USA, 5,000-fold dilution in blocking buffer (Nakalai Tesque, Inc., Japan)).

### Statistical analyses

Samples were analyzed in triplicate, and the data are presented as the mean ± standard deviation (SD). An analysis of variance (ANOVA) was used for data analysis. Different letters within a column indicate statistically significant differences by the Tukey-Kramer multiple range test (*p* < 0.05, CoStat version 6.400). A Student’s t-test was performed to examine differences between the two groups.

## Acknowledgments

Seeds of the *opr2-1opr3-3* double mutant and *coi1-1* mutant were provided by Professor Roberto Solano and Dr. Andrea Chini (CNB-CSIC, Spain).

## Author contributions

Conceptualization, M.U.; methodology, M.U., R.S., N.K., and Y.T.; validation, M.U., Y.T., R.S., and N.K..; formal analysis, R.S., Y.N., N.K., M.N., T.K., H.Y.; investigation, R.S., Y.N., N.K., M.N., T.K., H.Y.; resources, R.S., Y.N., N.K., M.N., T.K., H.Y.; data curation, M.U., Y.T., R.S., and N. K.; writing—original draft preparation, M.U.; writing—review and editing, M.U., Y.T., N.K., R.S.; visualization, Y.T., R.S., and M.N.; supervision, M.U.; project administration, M.U.; funding acquisition, M.U. All authors have read and agreed to the published version of the manuscript.

## Data availability statement

All data needed to evaluate the conclusions in the paper are present in the paper or the Supplementary Materials.

