## Supplementary material for "Downstream metabolites of (+)-*cis*-12-oxo-phytodienoic acid function as noncanonical bioactive jasmonates in *Arabidopsis thaliana*": 02May2024_supplementary_ddhJA.pdf

Supplementary Materials for

### Downstream metabolites of (+)-*cis*-12-oxo-phytodienoic acid function as bioactive jasmonates in *Arabidopsis thaliana* independent of canonical jasmonate signaling.

Rina Saito, Yuho Nishizato, Tsumugi Kitajima, Misuzu Nakayama, Yousuke Takaoka, Nobuki Kato, Minoru Ueda*

**This PDF file includes:**

Figs. S1 to S6
Tables S1
Supplementary Text
References

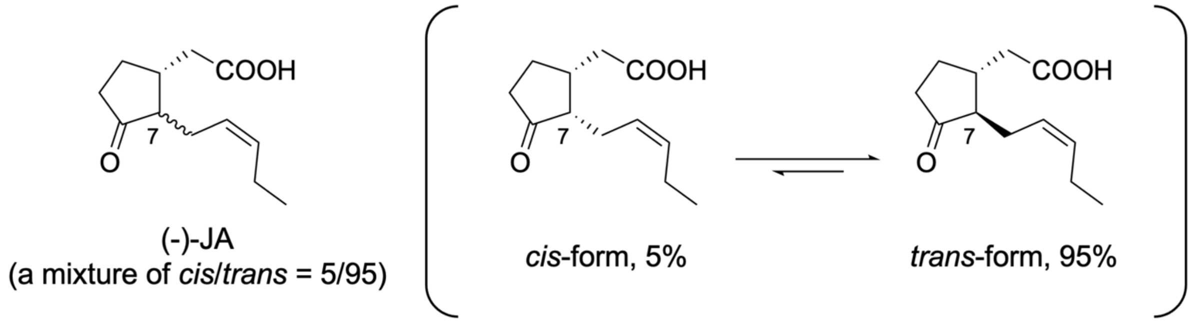


**Fig. S1. Chemical structure of (-)-JA used in this study.** A *cis*-form and a *trans*-form are in equilibrium in chemically synthesized (-)-JA.

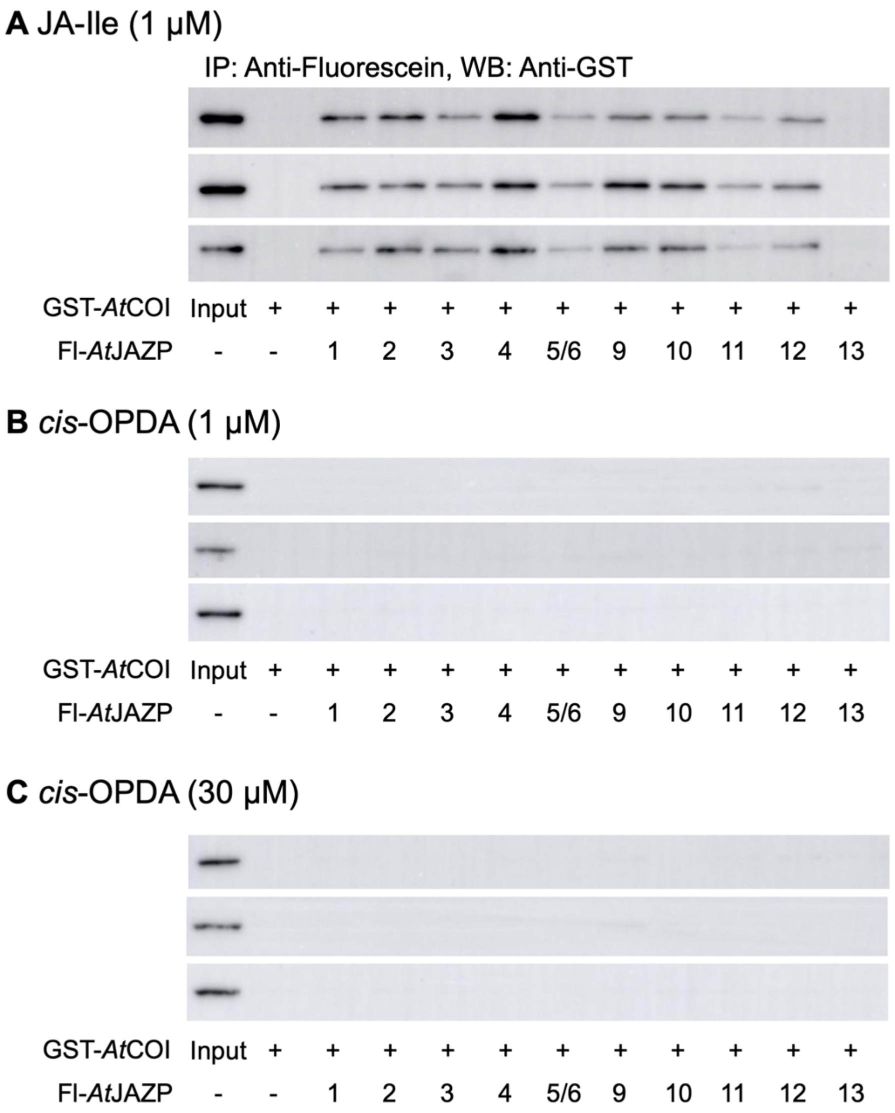


**Fig. S2. Results of three independent replications are shown in Figure 2I.** Pulldown assay of GST-*At*COI1 with Fl-*At*JAZPs in the presence of JA-Ile (1 μM, **A**), *cis*-OPDA (1 μM, **B**), or *cis*-OPDA (30 μM, **C**).

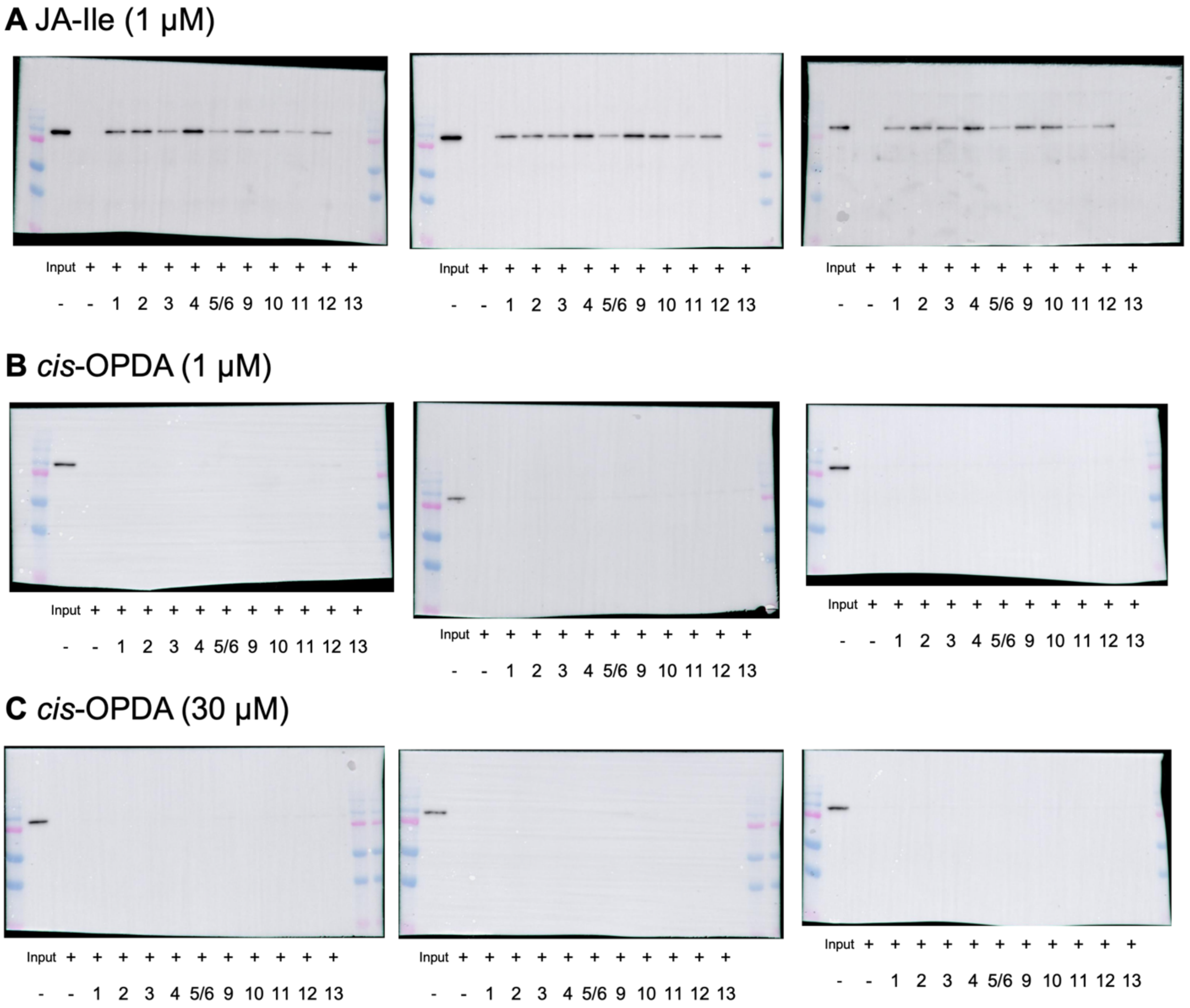


**Fig. S3. Uncropped images of Figure S2.** Pulldown assay of GST-*At*COI1 with Fl-*At*JAZPs in the presence of JA-Ile (1 µM, **A**), *cis*-OPDA (1 µM, **B**), or *cis*-OPDA (30 µM, **C**).

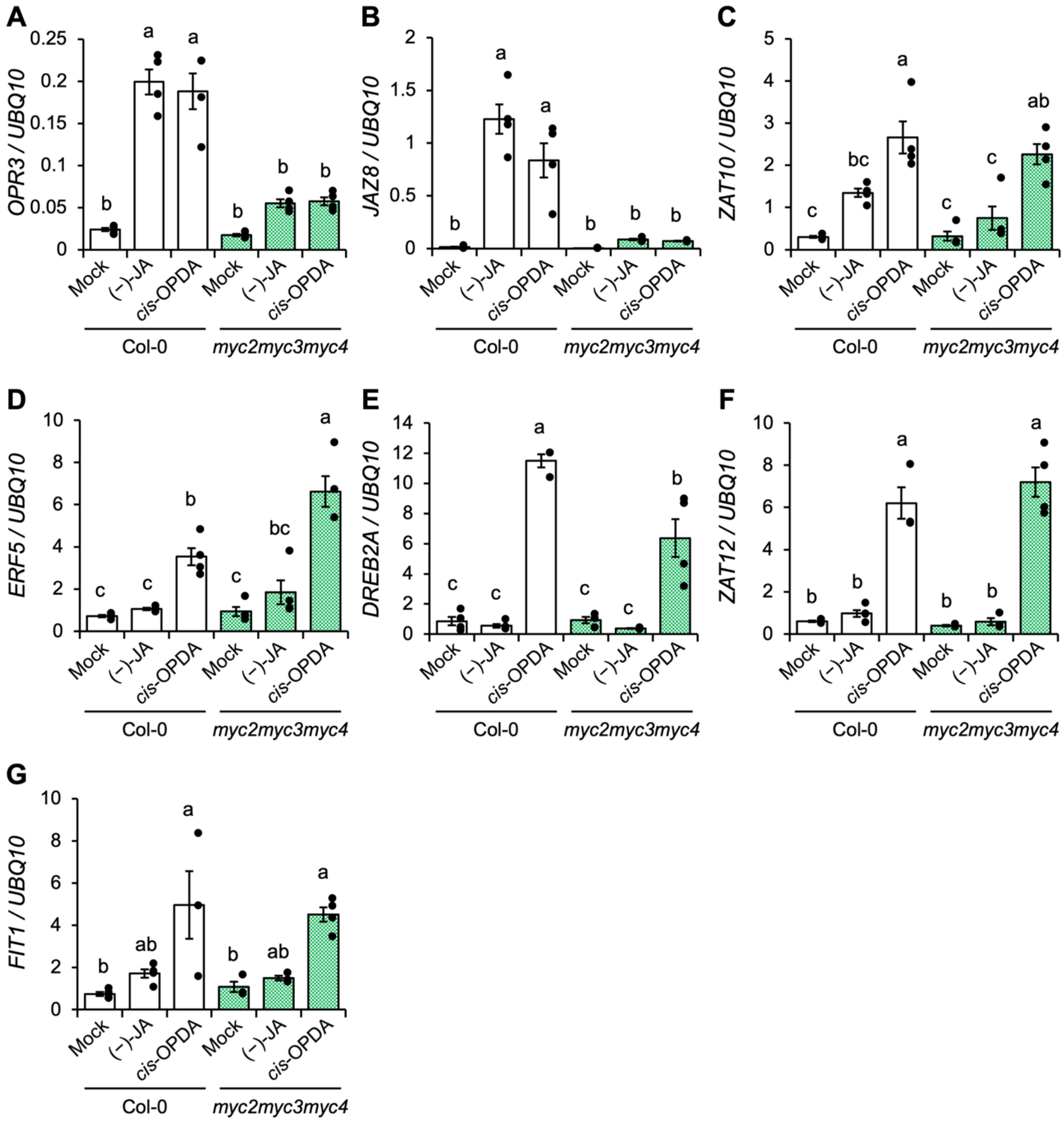


**Fig. S4. Gene expression analysis induced by *cis*-OPDA in WT and *myc2myc3myc4* mutant.** Gene expression analysis by RT-qPCR in 10-day-old WT (Col-0, white bar) and *myc2myc3myc4* mutant (green bar) with or without compounds ((-)-JA, *cis*-OPDA, 30 µM) treatment for 30 min. The results are mean with s.d. (n = 3–4). Samples were normalized to the *UBQ10* level. Significant differences were evaluated by ANOVA/Tukey Kramer test ($p < 0.05$). Experiments were repeated three times with similar results.

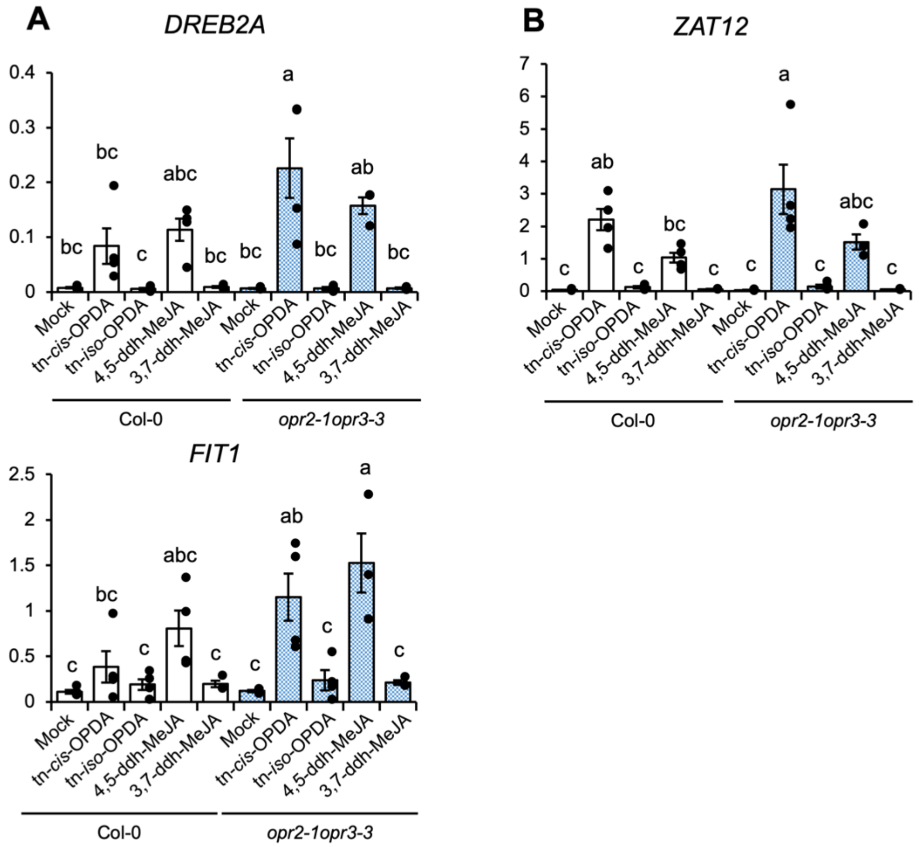


**Fig. S5. tn-*cis*-OPDA and 4,5-ddh-MeJA mediated gene expression of *DREB2A*, *ZAT12*, and *FIT1* through their electrophilic properties.** (**A-C**) Gene expression analysis by RT-qPCR in 10-day-old WT (Col-0, white bar) and *opr2-1opr3-3* mutant (blue bar) or without any treatments (mock) or treated with 30 μM tn-*cis*-OPDA, tn-*iso*-OPDA, 4,5-ddh-MeJA, and 3,7-ddh-MeJA for 30 min. The data are presented as mean ± SD (n = 3–4). Samples were normalized to the *UBQ10* level. Significant differences were evaluated by the ANOVA/Tukey Kramer test ($p < 0.05$). The experiments were repeated three times with similar results.

**Table S1. Gene sequences of all primers for RT-qPCR experiments used in this study.**

| Primer for RT-qPCR | Sequence |
|---|---|
| *UBQ10*_Fw | GGCCTTGTATAATCCCTGATGAATAAG |
| *UBQ10*_Rv | AAAGAGATAACAGGAACGGAAACATAGT |
| *OPR3*_Fw | CGGTTCAAGATTGATGGAGA |
| *OPR3*_Rv | CGATTATCAAACTCAGAGGC |
| *JAZ8*_Fw | CGGGTCGGATCCTCCAAAC |
| *JAZ8*_Rv | CGTCGTGAATGGTACGGTGAAG |
| *MYC2*_Fw | GTGCGGGATTAGCTGGTAAA |
| *MYC2*_Rv | ATGCATCCCAAACACTCCTC |
| *ZAT10*_Fw | CTCGGTTTGACTTTCCGGTCA |
| *ZAT10*_Rv | CAGTCAACAAATTCCTACACAACTCTC |
| *ERF5*_Fw | CCGCTTCTGTCGCCGTTATC |
| *ERF5*_Rv | CGTCCACGTCAGCATACACATC |
| *DREB2A*_Fw | GTTGCCAACGGTTCATACAG |
| *DREB2A*_Rv | CGTCGAAGAATCCATTACCATC |
| *ZAT12*_Fw | CACGGTGACTACGTTGAAGAAATC |
| *ZAT12*_Rv | CTCCAACTTGAGATTCAAATTGTC |
| *FIT1*_Fw | CTCCTTCTCCGGACACATACC |
| *FIT1*_Rv | CCTTGATTTAAAAGTGATCCAGTG |
| *HSP17.6A*_Fw | TCCTCCTGAGCCAAAGAAACC |
| *HSP17.6A*_Rv | CAACGAACACCAAGAGGTAG |
| *HSP17.4*_Fw | GTATGGAGAATGGGGTGTTGTCG |
| *HSP17.4*_Rv | GCTTTCCAACTTCAGAGTTCCTC |
| *HSP17.6II*_Fw | CTTCCTCCTCCGGAACCAAAG |
| *HSP17.6II*_Rv | CCATATCCCTCACGCATTCC |

#### Supplementary Text

##### General procedures of chemical syntheses

All chemical reagents and solvents were obtained from commercial suppliers (Kanto Chemical Co. Ltd., Wako Pure Chemical Industries Co. Ltd., Nacalai Tesque Co. Ltd., Tokyo Chemical Industry Co. Ltd., Sigma-Aldrich Co. LLC.) and used without further purification. All anhydrous solvents were either dried by standard techniques and freshly distilled before use or purchased in anhydrous form and used as supplied. Reversed-phase high–performance liquid chromatography (HPLC) was carried out on a PU–4180 plus pump equipped with UV–4075 and MD–4010 detectors (JASCO, Tokyo, Japan). $^{1}H$ and $^{13}C$ NMR spectra were recorded on a JNM–ECS–400 spectrometer (JEOL, Tokyo, Japan) in deuterated chloroform using TMS as an internal standard. Fourier transform infrared (FT/IR) spectra were recorded on an FT/IR–4100 (JASCO, Tokyo, Japan). High–resolution (HR) electrospray ionization (ESI)–mass spectrometry (MS) analyses were conducted using a microTOF II (Bruker Daltonics Inc., MA, USA). Optical rotations were measured using a JASCO P–2200 polarimeter (JASCO, Tokyo, Japan). Flash chromatography was performed on an Isolera system (Biotage Ltd., North Carolina, USA). TLC analyses were performed on Silica gel F254 (0.25 mm or 0.5 mm, MERCK, Germany) or RP–18F254S (0.25 mm, MERCK). All reactions were carried out under air unless stated otherwise.

~~**Synthesis of *cis*-OPDA-d$_5$:**~~

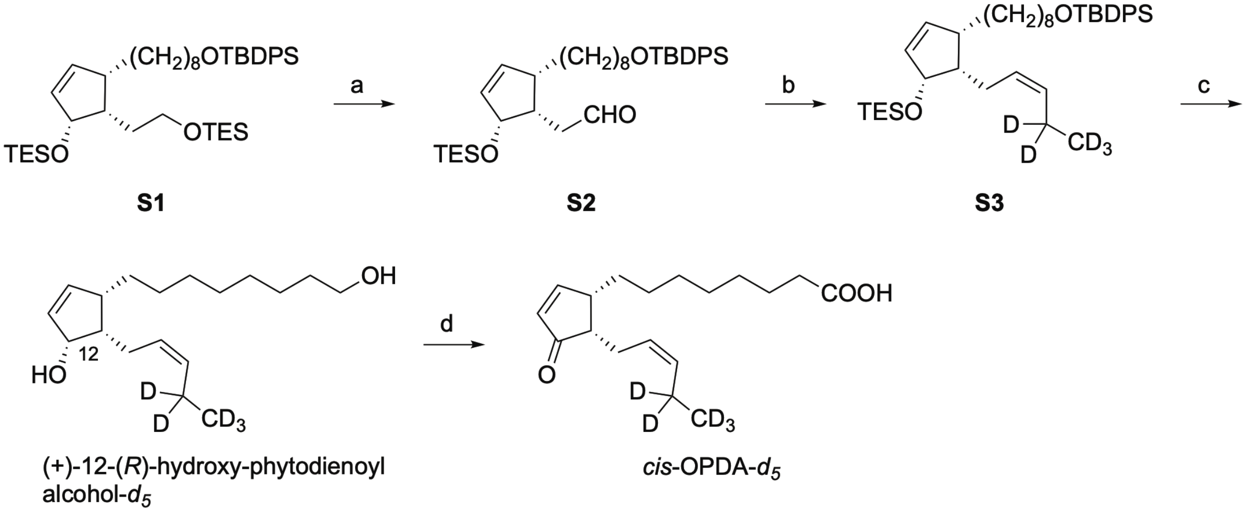


**Scheme S1. Synthesis of *cis*-OPDA-*d*$_5$.** *Reagents and conditions*: (a) $(COCl)_2$, DMSO, $CH_2Cl_2$; $NEt_3$; (b) $[Ph_3PCH_2CD_2CD_3]^+Br^-$, NaHMDS, THF, 57% (2 steps); (c) TBAF, THF, reflux, 94%; (d) Jones reagent, acetone, -20 °C, 42%.

**Synthesis of diene S3**. To a solution of DMSO (147 µL, 2.07 mmol) in $CH_2Cl_2$ (1.5 mL) was added oxalyl chloride (83.3 µL, 971 µmol) at -78 °C under an argon atmosphere. After the reaction mixture was stirred at -78 °C for 10 min, a solution of **S4** (134 mg, 185 µmol) in $CH_2Cl_2$ (1.7 mL) was slowly added. After stirring the reaction mixture at -65 °C for 1 h, $Et_3N$ (287 µL, 2.06 mmol) was slowly added. After the reaction mixture was stirred at -65 °C for 3 h, the mixture was gradually warmed to room temperature with stirring. The reaction mixture was quenched with saturated aqueous $NH_4Cl$. The mixture was extracted with *n*-hexane. The organic layer was washed with saturated aqueous NaCl, dried over $Na_2SO_4$, and filtered. The reaction mixture was concentrated under reduced pressure to afford the crude **S5** (123 mg) as a pale yellow oil. The crude product was used for the following reaction without further purification.

To an ice-cold suspension of $[Ph_3PCH_2CD_2CD_3]^+Br^-$ (277 mg, 708 µmol) in THF (2.6 mL) was added NaHMDS (343 µL, 1.9 M in THF, 652 µmol). The resulting orange-red mixture was stirred at room temperature for 40 min and cooled to -78 °C. To this solution was added a solution of the above aldehyde in THF (2.5 mL) dropwise. The resulting solution was stirred at -78 °C for 2 h and added DMF (401 µL), then at room temperature for 2 h, quenched with saturated $NH_4Cl$, extracted with *n*-hexane. The combined organic layers were dried over $Na_2SO_4$ and concentrated under reduced pressure. The residue was purified by medium-pressure chromatography (Isolera, eluent: *n*-hexane/EtOAc = 99:1 to *n*-hexane/EtOAc = 90:10) to give **S1**-*d*$_5$ (66.7 mg, 57% in 2 steps) as a colorless oil. $[\alpha]_D^{26}$ +0.37 (*c* 0.99, $CHCl_3$). $^1$H NMR (400 MHz, $CDCl_3$) $\delta_H$: 7.69-7.64 (m, 4H), 7.42-7.28 (m, 6H), 6.12 (dd, *J* = 5.7, 2.6 Hz, 1H), 5.84 (ddd, *J* = 5.7, 2.4, 1.2 Hz, 1H), 5.44 (dt, *J* = 10.8, 6.8 Hz, 1H), 5.35 (d, *J* = 10.8 Hz, 1H), 4.50 (dd, *J* = 6.8, 2.4 Hz, 1H), 3.64 (t, *J* = 6.6 Hz, 2H), 2.39 (brs, 1H), 2.20 (t, *J* = 6.8 Hz, 2H), 1.97 (dt, *J* = 13.6, 6.8 Hz 1H), 1.59-1.50 (m, 8H), 1.29-1.18 (m, 6H), 1.04 (s, 9H), 0.95 (t, *J* = 7.0 Hz, 9H), 0.57 (q, *J* = 7.0 Hz, 6H); $^{13}$C NMR (100 MHz, $CDCl_3$) $\delta_C$: 140.24, 135.6, 134.2, 132.7, 131.6, 129.5, 128.8, 127.6, 76.3, 64.0, 47.3, 46.0, 32.6, 32.4, 30.0, 29.7, 29.4, 28.0, 26.9, 25.8, 23.2, 22.6, 7.0, 5.2 (deuterated carbons were not detected); IR (neat) $cm^{-1}$: 2930, 1462, 1110, 739; HRMS (ESI, positive) *m/z* $[M+Na]^+$ Calcd. for $C_{40}H_{59}D_5NaO_2Si_2$: 660.4651, Found: 660.4641.

**Synthesis of (+)-12-(*R*)-hydroxy-phytodienoyl alcohol-*d5*.** To a solution of the above olefin **S1-** *d5* (66.7 mg, 105 μmol) in THF (15 mL) was added TBAF (1.8 mL, 1.0 M in THF, 1.8 mmol). The solution was heated under reflux for 2 h. After being cooled to room temperature, the solvent was removed under reduced pressure. The residue was purified by medium-pressure chromatography (Isolera, eluent: *n*-hexane/EtOAc = 90:10 to *n*-hexane/EtOAc = 20:10) to give (+)-12-(*R*)-hydroxy-phytodienoyl alcohol-*d5* (28.1 mg, 94%) as a colorless oil. $[\alpha]_D^{23}$ +35.5 (*c* 1.11, $CHCl_3$). $^1H$ NMR (400 MHz, $CDCl_3$) $\delta_H$: 6.23 (dd, $J$ = 5.8, 2.8 Hz, 1H), 5.96 (ddd, $J$ = 5.8, 2.2, 1.4 Hz, 1H), 5.46 (dt, $J$ = 11.0, 5.5 Hz, 1H), 5.42 (d, $J$ = 11.0 Hz, 1H), 4.51 (brs, 1H), 3.64 (t, $J$ = 6.6 Hz, 2H), 2.43-2.51 (m, 1H), 2.36-2.29 (m, 1H), 2.22-2.16 (m, 1H), 2.12-2.05 (m, 1H), 1.46-1.04 (m, 14H); $^{13}C$ NMR (100 MHz, $CDCl_3$) $\delta_C$: 141.82, 132.43, 132.07, 128.06, 76.66, 63.07, 46.22, 46.10, 33.64, 32.82, 29.92, 29.62, 29.45, 28.15, 25.78, 23.15, 19.92 (quintet, $J_{C\text{-}D}$ = 18.8 Hz), 13.20 (septet, $J_{C\text{-}D}$ = 19.3 Hz); IR (neat) $cm^{-1}$: 3350, 2855, 2927, 1057; HRMS (ESI, positive) *m/z* $[M+Na]^+$ Calcd. for $C_{18}H_{27}D_5NaO_2$: 308.2614, Found: 308.2604.

***Synthesis of tn-cis-OPDA*:**

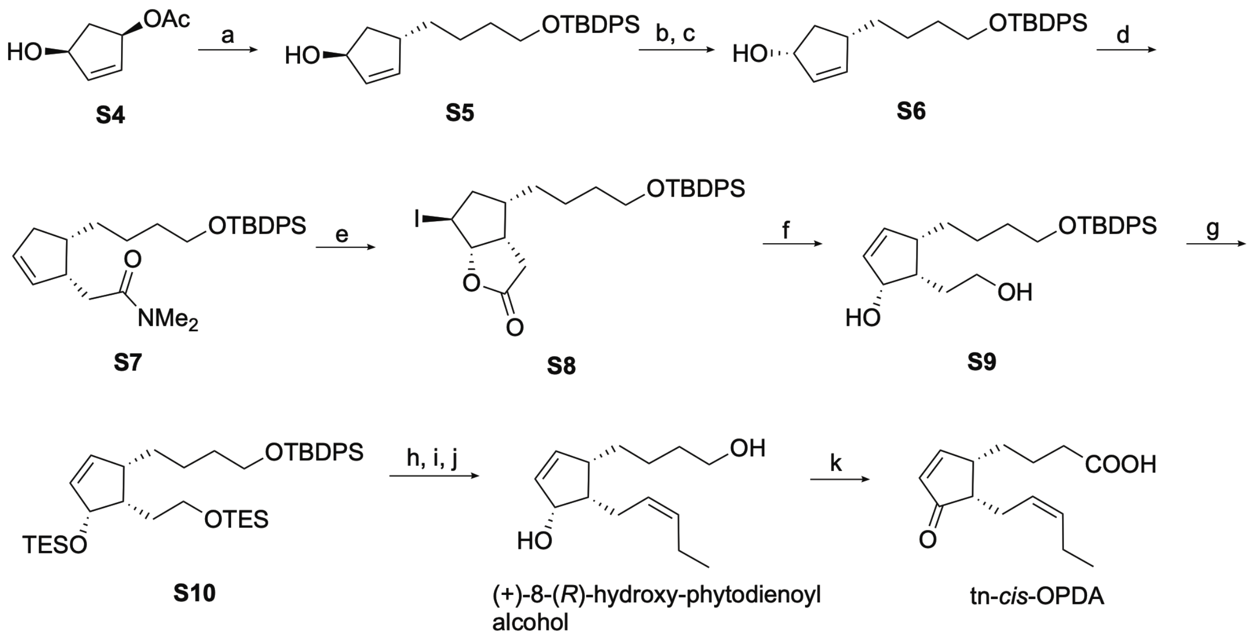


**Scheme S2. Synthesis of tn-*cis*-OPDA.** *Reagents and conditions*: (a) TBDPSO$(CH_2)_4$MgCl, CuCN, THF, -18 °C, 83%; (b) $Ph_3P$, AcOH, DIAD, toluene, -78 °C; (c) LiOH, $H_2O$, MeOH, THF, 44% (2 steps); (d) MeC$(OMe)_2$$NMe_2$, xylene, reflux, 67%; (e) $I_2$, buffer (pH 6.0), THF, 78%; (f) DBU, THF, reflux; $LiAlH_4$, -30 °C, 55%; (g) TESCl, imidazole, DMF, 82%; (h) $(COCl)_2$, DMSO, $CH_2Cl_2$, -78 °C; $NEt_3$; (i) $Ph_3P^+CH_2CH_2CH_3Br^-$, NaHMDS, DMF, THF; (j) TBAF, THF, reflux, 66% (3 steps); (k) Jones reagent, acetone, -20 °C, 96%.

**Synthesis of cyclopentenol S5.** To CuCN (222 mg, 2.47 mmol) was added TBDPSO$(CH_2)_4$MgCl (1.22 M in THF, 14 mL, 37.7 mmol) (*3*) slowly at -30 °C under an argon atmosphere. After 10 min stirring at -30 °C, acetate **S4** (1.74 g, 12.2 mmol) in THF (26 mL) was added dropwise. The mixture was warmed to -18 °C over 90 min and quenched by saturated aqueous $NH_4Cl$. The water layer was extracted with EtOAc. The combined organic layers were washed with saturated aqueous NaCl, dried over $Na_2SO_4$, and filtered. The residue was purified by medium-pressure chromatography (Isolela, eluent: *n*-hexane/EtOAc = 94:6 to *n*-hexane/EtOAc = 50:50) to give **S10** (4.01 g, 83%) as a yellow oil. $[\alpha]_D^{27}$ -62.6 (*c* 1.03, $CHCl_3$); $^1$H NMR (400 MHz, $CDCl_3$) $\delta_H$: 7.72-7.60 (m, 4H), 7.47-7.32 (m, 6H), 5.94 (ddd, *J* = 5.7, 2.1, 0.5 Hz, 1H), 5.82 (dt, *J* = 5.7, 2.1 Hz, 1H), 4.90-4.79 (m, 1H), 3.65 (t, *J* = 6.4 Hz, 2H), 2.90-2.78 (m, 1H), 1.90 (ddd, *J* = 14.1, 7.6, 2.8 Hz, 1H), 1.75 (ddd, *J* = 14.1, 7.1, 5.3 Hz, 1H), 1.61-1.51 (m, 2H), 1.43-1.18 (m, 4H), 1.05 (s, 9H); $^{13}$C NMR (100 MHz, $CDCl_3$) $\delta_C$: 140.38, 135.73, 134.23, 132.57, 129.66, 127.73, 77.34, 63.90, 44.17, 40.74, 35.70, 32.76, 27.00, 24.26, 19.36; IR (neat) cm$^{-1}$: 3330, 3070, 1110, 823; HRMS (ESI, positive) *m/z* $[M+Na]^+$ Calcd. for $C_{25}H_{34}NaO_2Si$: 417.2220, Found: 417.2210.

**Synthesis of cyclopentenol S6.** To a solution of alcohol **S5** (4.01 g, 10.2 mmol), $Ph_3P$ (4.50 g, 17.2 mmol), and AcOH (1 mL, 17.5 mmol) in toluene (88 mL) was added DIAD (6 mL, 28.0 mmol) at -78 °C under an argon atmosphere. The reaction was carried out at the same temperature for 3 h and quenched with saturated aqueous $NaHCO_3$. After being vigorously stirred at room temperature, the water layer was extracted with *n*-hexane. The combined organic layers were washed with saturated aqueous NaCl, dried over $Na_2SO_4$, and filtered. The residue was roughly purified by medium-pressure chromatography (Isolela, eluent: 98:2 *n*-hexane/EtOAc to 80:20 *n*-hexane/EtOAc) to give the mixture as a colorless oil (2.45 g). To a solution of the mixture (2.45 g) in MeOH (24 mL) and THF (55 mL) was added 1M LiOH in $H_2O$ (22.5 mL, 22.5 mmol). After stirring at room temperature for 2.5 h, the organic solvent was removed under reduced pressure. The resulting mixture was extracted with $Et_2O$. The combined organic layers were washed with saturated aqueous NaCl, dried over $Na_2SO_4$, and filtered. The residue was purified by medium-pressure chromatography (Isolela, eluent: *n*-hexane/EtOAc = 93:7 to *n*-hexane/EtOAc = 40:60) to give **S6** (1.77 g, 44% in 2 steps) as a yellow oil. $[\alpha]_D^{26}$ -10.5 (*c* 0.85, $CHCl_3$); $^1$H NMR (400 MHz, $CDCl_3$) $\delta_H$: 7.72-7.60 (m, 4H), 7.47-7.32 (m, 6H), 5.87 (dt, *J* = 5.6, 1.6 Hz, 1H), 5.77 (dt, *J* = 5.6, 2.0 Hz, 1H), 4.85-4.75 (m, 1H), 3.66 (t, *J* = 6.4 Hz, 2H), 2.58-2.42 (m, 2H), 1.61-1.51 (m, 2H), 1.50-1.15 (m, 5H), 1.05 (s, 9H); $^{13}$C NMR (100 MHz, $CDCl_3$) $\delta_C$: 138.82, 135.73, 134.24, 133.09, 129.66, 127.74, 77.36, 63.94, 44.56, 40.62, 36.71, 32.75, 27.01, 24.20, 19.36; IR (neat) cm$^{-1}$: 3336, 3049, 1110, 823; HRMS (ESI, positive) *m/z* $[M+Na]^+$ Calcd. for $C_{25}H_{34}NaO_2Si$: 417.2220, Found: 417.2214.

**Synthesis of *N*, *N*-dimethyl acetamide S7.** A solution of alcohol **S6** (1.77g, 4.49 mmol) and $MeC(OMe)_2NMe_2$ (90% purity, 3.8 mL, 23.4 mmol) in xylene (55 mL) was stirred at reflux temperature for 4 h under an argon atmosphere, added $MeC(OMe)_2NMe_2$ (90% purity, 4 mL, 24.6 mmol) again, and stirred at reflux temperature for an additional 6 h. The solvent was removed under reduced pressure. The residue was purified by medium-pressure chromatography (Isolela, eluent: *n*-hexane/EtOAc = 95:5 to *n*-hexane/EtOAc = 60:40) to give **S7** (1.40 g, 67%) as a brown oil. $[\alpha]_D^{26}$ -56.6 (*c* 0.87, $CHCl_3$); $^1$H NMR (400 MHz, $CDCl_3$) $\delta_H$: 7.72-7.60 (m, 4H), 7.47-7.32 (m, 6H), 5.85-5.78 (m, 1H), 5.76-5.70 (m, 1H), 3.66 (t, *J* = 6.4 Hz, 2H), 3.15-3.02 (m, 1H), 2.96 (s, 3H), 2.94 (s, 3H), 2.44-2.18 (m, 3H), 2.10 (dd, *J* = 14.6, 10.2 Hz, 1H), 1.95 (ddq, *J* = 15.9, 8.5, 2.4 Hz, 1H), 1.68-1.16 (m, 6H), 1.04 (s, 9H); $^{13}$C NMR (100 MHz, $CDCl_3$) $\delta_C$: 172.61, 135.77, 135.52, 134.06, 130.21, 129.47, 127.53, 63.87, 43.45, 41.29, 37.42, 37.10, 35.42, 33.17, 32.82, 30.30, 26.87, 24.95, 19.21; IR (neat) cm$^{-1}$: 3049, 1652, 1111, 823; HRMS (ESI, positive) *m/z* $[M+Na]^+$ Calcd. for $C_{29}H_{41}NNaO_2Si$; 486.2799, Found: 486.2790.

**Synthesis of Iodo lactone S8.** To a solution of acetamide **S7** (1.40 g, 3.01 mmol) in THF (22 mL) and buffer (pH 6.0, 22 mL) was added $I_2$ (1.53 g, 6.03 mmol). The solution was stirred at room temperature for 16 h and quenched with saturated $Na_2S_2O_3$. The water layer was extracted with $Et_2O$. The combined organic layers were washed with saturated aqueous NaCl, dried over $Na_2SO_4$, and filtered. The residue was purified by medium-pressure chromatography (Isolela, eluent: *n*-hexane/EtOAc = 95:5 to *n*-hexane/EtOAc = 60:40) to give **S8** (1.33 g, 79 %) as a pale yellow solid. $[\alpha]_D^{25}$ +4.1 (*c* 0.94, $CHCl_3$); $^1$H NMR (400 MHz, $CDCl_3$) $\delta_H$: 7.72-7.60 (m, 4H), 7.47-7.32 (m, 6H), 5.26 (d, *J* = 6.4 Hz, 1H), 4.45 (d, *J* = 5.2 Hz, 1H), 3.67 (t, *J* = 6.2 Hz, 2H), 3.15-3.03 (m, 1H), 2.73-2.58 (m, 1H), 2.57 (dd, *J* = 18.8, 10.0 Hz, 1H), 2.47 (dd, *J* = 18.8, 3.6 Hz, 1H), 2.07 (dd, *J* = 14.8, 6.0 Hz, 1H), 1.68-1.23 (m, 7H), 1.05 (s, 9H); $^{13}$C NMR (100 MHz, $CDCl_3$) $\delta_C$: 176.61, 135.65, 134.02, 129.70, 127.73, 92.81, 63.56, 40.39, 40.19, 38.86, 32.55, 29.77, 28.75, 28.23, 26.97, 24.72, 19.29; IR (film) cm$^{-1}$: 2931, 1786, 1111, 823; HRMS (ESI, positive) *m/z* $[M+Na]^+$ Calcd. for $C_{27}H_{35}INaO_3Si$: 585.1292, Found: 585.1279.

**Synthesis of diol S9.** To a solution of **S8** (1.33 g, 2.36 mmol) in THF (15 mL) was added DBU (0.46 mL, 3.08 mmol). After being stirred at reflux temperature for 16.5 h, the mixture was cooled to -30 °C, added $LiAlH_4$ (271 mg, 7.15 mmol), and stirred for 45 min. The reacting solution was quenched by EtOAc and $SiO_2/H_2O$ (10/3, 11.1 g). The solution was filtrated and concentrated. The residue was purified by medium-pressure chromatography (Isolela, eluent: *n*-hexane/EtOAc = 90:10 to *n*-hexane/EtOAc = 40:60) to give **S9** (566 mg, 55%) as a yellow oil. $[\alpha]_D^{27}$ +38.4 (*c* 0.91, $CHCl_3$); $^1$H NMR (400 MHz, $CDCl_3$) $\delta_H$: 7.72-7.60 (m, 4H), 7.47-7.32 (m, 6H), 6.18 (dd, *J* = 5.7, 2.8 Hz, 1H), 5.96 (ddd, *J* = 5.7, 2.4, 1.4 Hz, 1H), 4.60 (m, 1H), 3.89 (dt, *J* = 9.8, 4.9 Hz, 1H), 3.76 (td, *J* = 9.8, 3.9 Hz, 1H), 3.65 (td, *J* = 6.3, 0.9 Hz, 2H), 2.53-2.39 (m, 1H), 2.19-2.18 (m, 3H), 1.73-1.06 (m, 6H), 1.05 (s, 9H); $^{13}$C NMR (100 MHz, $CDCl_3$) $\delta_C$: 140.83, 135.40, 133.92, 131.69, 129.34, 127.41, 76.19, 63.63, 62.68, 46.59, 44.42, 33.11, 32.58, 27.79, 26.69, 24.10, 19.04; IR (neat) cm$^{-1}$: 3349, 2932, 1110, 823; HRMS (ESI positive) m/z $[M+Na]^+$ Calcd. for $C_{27}H_{38}NaO_3Si$: 461.2482, Found: 461.2475.

**Synthesis of bis-TES ether S10.** To a solution of **S9** (585 mg, 1.33 mmol) in DMF (9 mL) was added TESCl (0.74 mL, 4.42 mmol) and imidazole (390 mg, 5.72 mmol). After stirring at room temperature for 20 h under an argon atmosphere, the reaction was diluted with $H_2O$ with vigorous stirring. The water layer was extracted with EtOAc. The combined organic layers were washed with saturated aqueous NaCl, dried over $Na_2SO_4$, and filtered. The residue was purified by medium-pressure chromatography (Isolela, eluent: *n*-hexane/EtOAc = 99:1 to *n*-hexane/EtOAc = 90:10) to give **S10** (729 mg, 82%) as a colorless oil. $[\alpha]_D^{26}$ +0.6 (*c* 1.14, $CHCl_3$); $^1$H NMR (400 MHz, $CDCl_3$) $\delta_H$: 7.72-7.60 (m, 4H), 7.47-7.32 (m, 6H), 6.10 (dd, *J* = 6.0, 2.8 Hz, 1H), 5.96 (ddd, *J* = 6.0, 2.2, 1.2 Hz, 1H), 4.46 (dd, *J* = 5.8, 2.2 Hz, 1H), 3.76-3.56 (m, 4H), 2.44-2.28 (m, 1H), 2.07 (quintet, *J* = 7.0 Hz, 1H), 1.78 (dq, *J* = 13.5, 7.0 Hz, 1H), 1.69-1.07 (m, 7H), 1.04 (s, 9H), 0.96 (t, *J* = 8.0 Hz, 9H), 0.93 (t, *J* = 8.0 Hz, 9H), 0.60 (q, *J* = 8.0 Hz, 6H), 0.55 (q, *J* = 8.0 Hz, 6H); $^{13}$C NMR (100 MHz, $CDCl_3$) $\delta_C$: 140.29, 135.72, 134.34, 132.87, 129.62, 127.71, 76.29, 64.17, 62.14, 46.19, 42.92, 33.14, 32.54, 28.90, 27.01, 24.31, 19.36, 7.07, 6.97, 5.36, 4.57; IR (neat) cm$^{-1}$: 3050, 2954, 1109, 823; HRMS (ESI positive) m/z $[M+Na]^+$ Calcd. for $C_{39}H_{66}NaO_3Si_3$: 689.4212, Found: 689.4191.

**Synthesis of (+)-8-(*R*)-hydroxy-phytodienoyl alcohol.** To a solution of DMSO (890 µL, 12.5 mmol) in $CH_2Cl_2$ (9.5 mL) was added oxalyl chloride (520 µL, 6.06 mmol) at -78 °C under an argon atmosphere. After the reaction mixture was stirred at -78 °C for 10 min, a solution of **S10** (723 mg, 1.08 mmol) in $CH_2Cl_2$ (10 mL) was slowly added. After stirring the reaction mixture at -65 °C for 1 h, $Et_3N$ (1.7 mL, 12.3 mmol) was slowly added. After the reaction mixture was stirred at -65 °C for 1 h, the mixture was gradually warmed to room temperature with stirring for 2 h. The reaction mixture was quenched with saturated aqueous $NH_4Cl$. The mixture was extracted with *n*-hexane. The organic layer was washed with saturated aqueous NaCl, dried over $Na_2SO_4$, and filtered. The reaction mixture was concentrated under reduced pressure to afford the mixture (706 mg) as a brown oil. The crude product was used for the next reaction without further purification.

A solution of $Ph_3P^+CH_2CH_2CH_2CH_3\ Br^-$ (1.27 g, 3.30 mmol) in THF (28 mL) was cooled to 0 °C and stirred under an argon atmosphere. After 20 min, NaHMDS (1.9 M in THF, 2.0 mL, 3.8 mmol) was added and warmed to room temperature for 40 min. Then, the mixture was cooled to -78 °C and added DMF (2.1 mL) and the previous reaction mixture (706 mg) in THF (13 mL). The mixture was warmed to 0 °C and stirred for 4.5 h. The reaction mixture was quenched with saturated aqueous $NH_4Cl$. The mixture was extracted with EtOAc. The organic layer was washed with saturated aqueous NaCl, dried over $Na_2SO_4$, and filtered. The reaction mixture was purified by medium-pressure chromatography (Isolela, eluent: *n*-hexane to *n*-hexane/EtOAc = 94:6) to give a mixture (552 mg) as a yellow oil. This mixture was used for the following reaction without further purification.

To a solution of a mixture (552 mg) in THF (136 mL) was added TBAF (1M in THF, 9.57 mL, 9.57 mmol) and stirred at reflux temperature for 3 h under an argon atmosphere. The solvent was removed under reduced pressure. The reaction mixture was roughly purified by medium-pressure chromatography (Isolela, eluent: 99:1 $CH_3Cl$/MeOH to 90:10 $CH_3Cl$/MeOH) to give (+)-8-(*R*)-hydroxy-phytodienoyl alcohol (161 mg, 66% in 3 steps) as a brown oil. $[\alpha]_D^{25}$ +55.4 (*c* 1.24, $CHCl_3$). $^1$H NMR (400 MHz, $CDCl_3$) $\delta_{H:}$ 6.24 (dd, *J* = 5.8, 2.6 Hz, 1H), 5.97 (ddd, *J* = 5.8, 2.5, 1.3 Hz, 1H), 5.50-5.37 (m, 2H), 4.52 (dd, *J* = 6.0, 2.4 Hz, 1H), 3.65 (t, *J* = 6.4 Hz, 2H), 2.55-2.44 (m, 1H), 2.40-2.27 (m, 1H), 2.25-1.93 (m, 4H), 1.73-1.41 (m, 4H), 1.40-1.20 (m, 3H), 1.20-1.05 (m, 1H), 0.99 (t, *J* = 7.6 Hz, 3H); $^{13}$C NMR (100 MHz, $CDCl_3$) $\delta_C$: 140.51, 132.59, 132.22, 127.85, 76.53, 62.91, 46.14, 45.99, 33.33, 33.04, 24.27, 23.08, 20.81, 14.28; IR (neat) cm$^{-1}$: 3365, 2933, 1459, 742; HRMS (ESI positive) m/z $[M+Na]^+$ Calcd. for $C_{14}H_{24}NaO_2$ 247.1668, Found 247.1665.

***Synthesis of 4,5-ddh-MeJA*:**

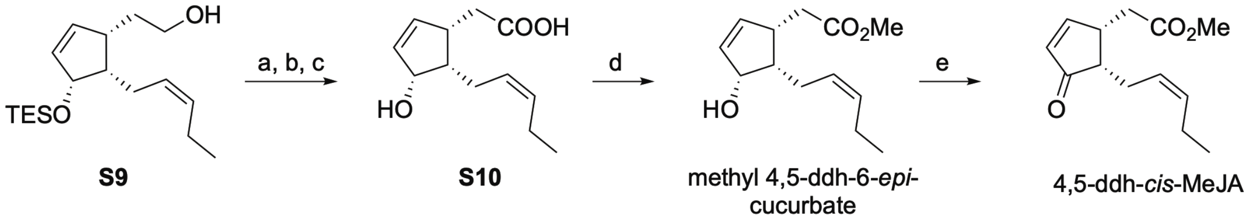


**Scheme S3. Synthesis of 4,5-ddh-MeJA.** *Reagents and conditions*: (a) Dess-Martin periodinane, $CH_2Cl_2$; (b) $NaClO_2$, $NaH_2PO_4$, 2-methyl-2-butene, *t*BuOH, $H_2O$; (c) TBAF, THF, quant. (3 steps); (d) $TMSCHN_2$, MeOH, benzene, 62%; (e) Jones reagent, acetone, -20 °C, 55%.

**Synthesis of carboxylic acid S12.** To a solution of **S11** (30.0 mg, 96.6 µmol) (*1*, *2*) in $CH_2Cl_2$ (3 mL) was added Dess-Martin periodinane (63.2 mg, 149 µmol). After being stirred for 80 min, the reaction mixture was quenched with saturated aqueous $NaHCO_3$ and extracted with *n*-hexane. The organic layer was washed with saturated aqueous NaCl, dried over $Na_2SO_4$, and filtered. The reaction mixture was concentrated under reduced pressure to afford the aldehyde (37.2 mg, mixture) as a colorless oil. The crude product was used for the following reaction without further purification.

To a solution of the above aldehyde and 2-methyl-2-butene (1.8 ml) in *t*BuOH (5.3 ml) were added $H_2O$ (1.3 ml), $NaH_2PO_4 \cdot 2H_2O$ (469 mg, 3.00 mmol) and $NaClO_2$ (114 mg, 1.26 mmol) and the mixture was stirred at room temperature for 70 min. The reaction was quenched with saturated aqueous $NH_4Cl$. The water layer was extracted with EtOAc, and the combined organic layers were dried over $Na_2SO_4$ and concentrated under reduced pressure to afford the carboxylic acid (65.2 mg, mixture) as a colorless oil. The crude product was used for the following reaction without further purification.

To a solution of the above carboxylic acid in THF (6.7 mL) was added 1 M TBAF in THF (330 µL, 330 µmol). After stirring at room temperature for 1 h, the solvent was removed under reduced pressure. The residue was purified by medium-pressure chromatography (Isolera, eluent: AcOH/*n*-hexane/EtOAc = 0.1:88:12 to AcOH/EtOAc = 0.1:99.9) to give **S8** (21.8 mg, quant. in 3 steps) as a colorless oil. $[\alpha]_D^{23}$ +44.4 (*c* 0.70, $CHCl_3$). $^1H$ NMR (400 MHz, $CDCl_3$) $\delta_H$: 6.15 (dd, *J* = 5.8, 2.7 Hz, 1H), 6.01 (dd, *J* = 5.8, 2.5 Hz, 1H), 5.50-5.38 (m, 2H), 4.55 (dd, *J* = 5.6, 2.7 Hz, 1H), 3.02-2.91 (m, 1H), 2.63 (dd, *J* = 16.1, 5.4 Hz, 1H), 2.39-2.06 (m, 6H), 0.99 (t, *J* = 7.6 Hz, 3H); $^{13}C$ NMR (100 MHz, $CDCl_3$) $\delta_C$: 178.36, 139.32, 133.26, 133.08, 127.04, 76.29, 45.07, 42.32, 36.37, 23.15, 20.78, 14.15; IR (film) $cm^{-1}$: 3377, 3058, 2963, 2931, 1715, 1702; HRMS (ESI, negative) *m/z* $[M-H]^-$ Calcd. for $C_{12}H_{17}O_3$: 209.1183, Found: 209.1186.

**Synthesis of (+)-methyl 4,5-ddh-6-*epi*-cucurbate.** To a solution of **S8** (15.0 mg, 71.4 µmol) in MeOH (1 mL) and benzene (1 mL) was added TMS diazomethane solution (0.6 M in *n*–hexane, 600 µL, 0.36 mmol) at 0 °C for 10 min, the solvent was removed under reduced pressure. The residue was purified by medium-pressure chromatography (Isolera, eluent: *n*-hexane/EtOAc = 98:2 to *n*-hexane/EtOAc = 80:20) to give 4,5-ddh-6-*epi*-cucurbate (10.0 mg, 62%) as a colorless oil. $[\alpha]_D^{25}$ +38.4 (*c* 0.50, $CHCl_3$). $^1H$ NMR (400 MHz, $CDCl_3$) $\delta_H$: 6.09 (dd, *J* = 5.9, 2.8 Hz, 1H), 6.00 (dd, *J* = 5.9, 2.6 Hz, 1H), 5.50-5.38 (m, 2H), 4.53 (brdd, *J* = 5.3, 2.4 Hz, 1H), 3.03-2.90 (m, 1H), 2.58 (dd, *J* = 16.0, 5.6 Hz, 1H), 2.39-2.03 (m, 6H), 0.98 (t, *J* = 7.4 Hz, 3H); $^{13}C$ NMR (100

MHz, $CDCl_3$) $\delta_C$: 173.87, 138.92, 133.43, 132.88, 127.24, 76.30, 51.62, 45.08, 42.62, 36.27, 23.22, 20.80, 14.18; IR (film) $cm^{-1}$: 3440, 2920, 1733, 1718, 1438, 1168; HRMS (ESI, positive) *m/z* $[M-H]^-$ Calcd. for $C_{13}H_{20}NaO_3$: 247.1310, Found: 247.1305.

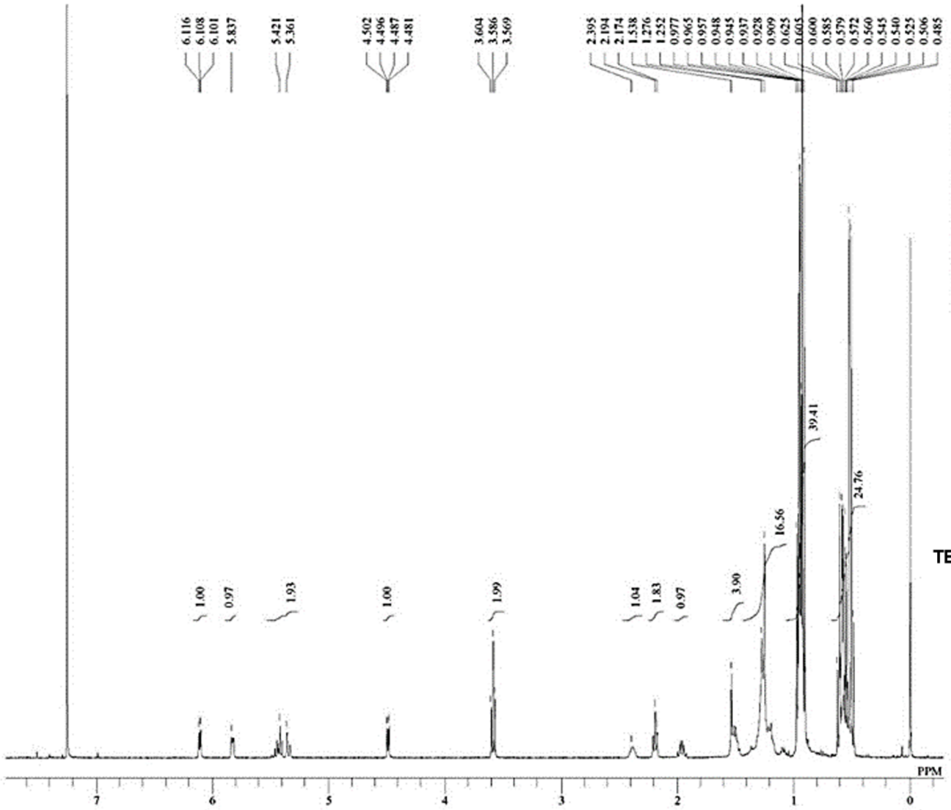


DFILE OPDA-d5-Ile TES protection(S
COMNT single_pulse
DATIM 21-07-2022 21:17:00
OBNUC 1H
EXMOD proton.jxp
OBFRQ 399.78 MHz
OBSET 4.19 KHz
OBFIN 7.29 Hz
POINT 16400
FREQU 7503.00 Hz
SCANS 8
ACQTM 2.1837 sec
PD 5.0000 sec
PW1 3.40 usec
IRNUC 1H
CTEMP 22.2 c
SLVNT CDCL3
EXREF 0.00 ppm
BF 1.00 Hz
RGAIN 74

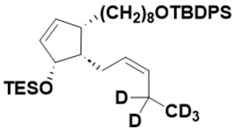


S3

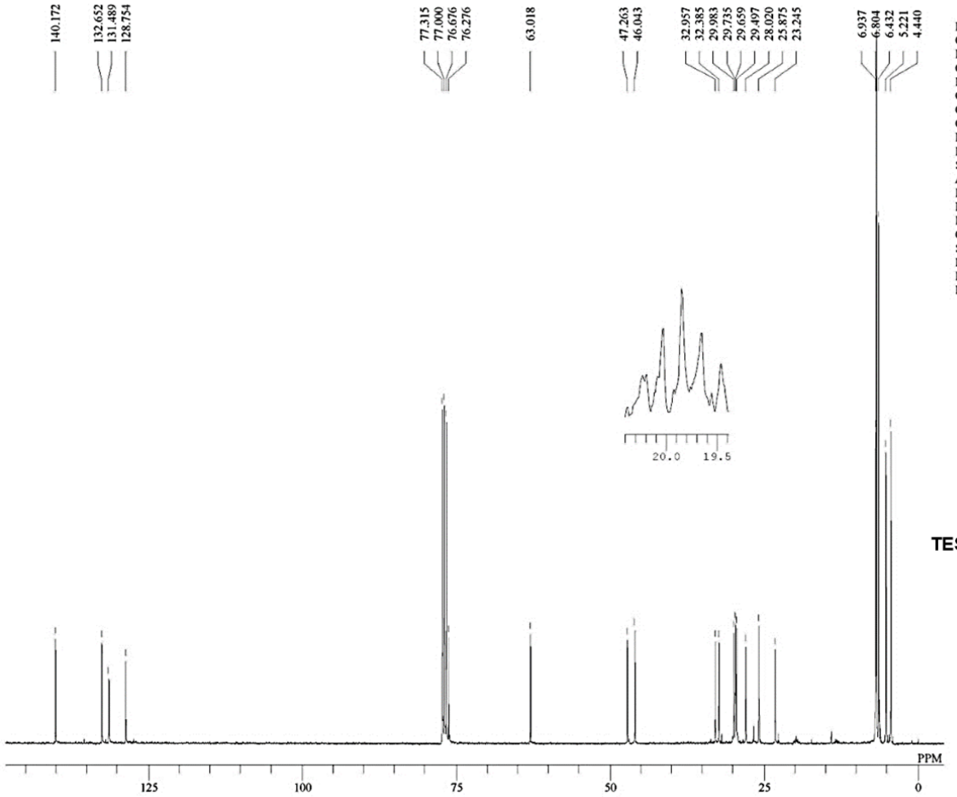


DFILE OPDA-d5-Ile TES protection(S
COMNT single pulse decoupled gated N
DATIM 22-07-2022 00:07:52
OBNUC 13C
EXMOD carbon.jxp
OBFRQ 100.53 MHz
OBSET 5.35 KHz
OBFIN 5.86 Hz
POINT 32780
FREQU 31407.04 Hz
SCANS 12342
ACQTM 1.0433 sec
PD 2.0000 sec
PW1 3.53 usec
IRNUC 1H
CTEMP 21.9 c
SLVNT CDCL3
EXREF 77.00 ppm
BF 1.00 Hz
RGAIN 50

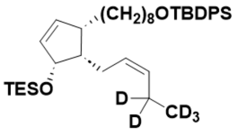


S3

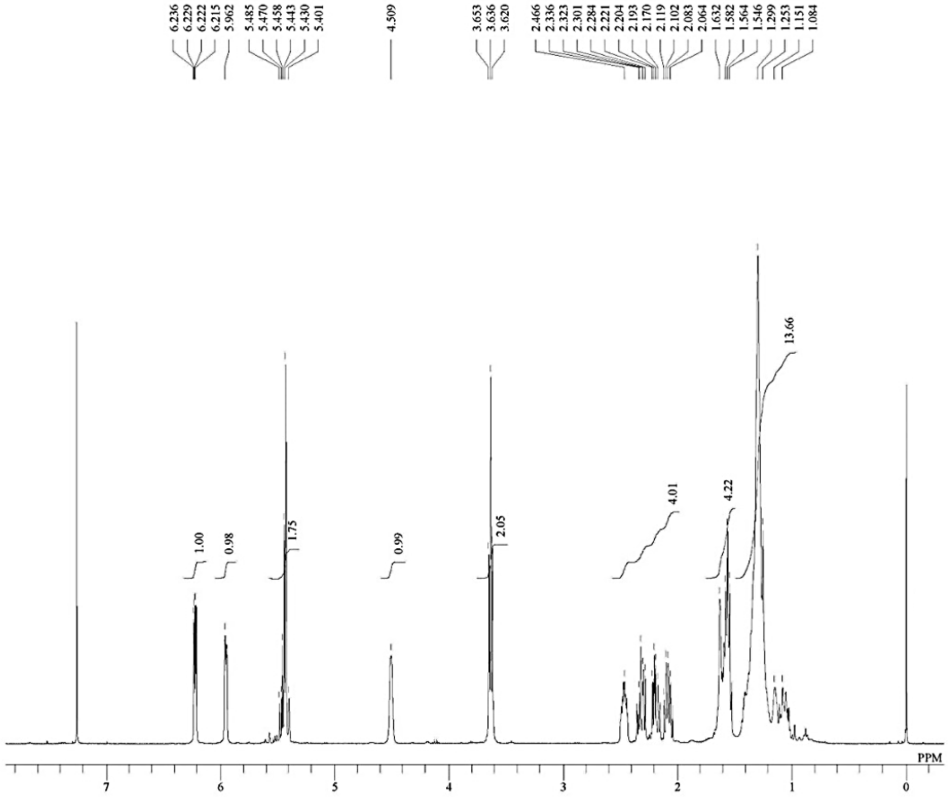


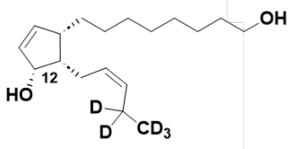


**(+)-12-(*R*)-hydroxy-phytodienoyl alcohol-$d_5$**

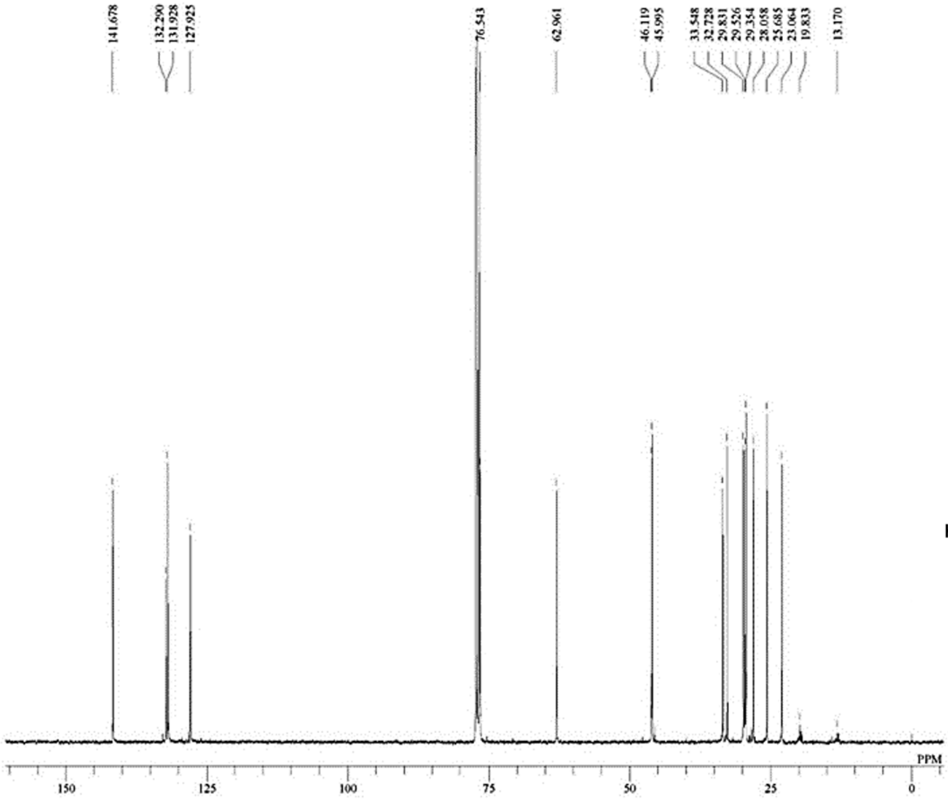


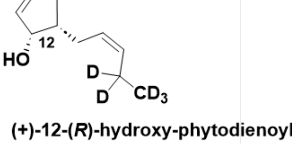


**(+)-12-(*R*)-hydroxy-phytodienoyl alcohol-$d_5$**

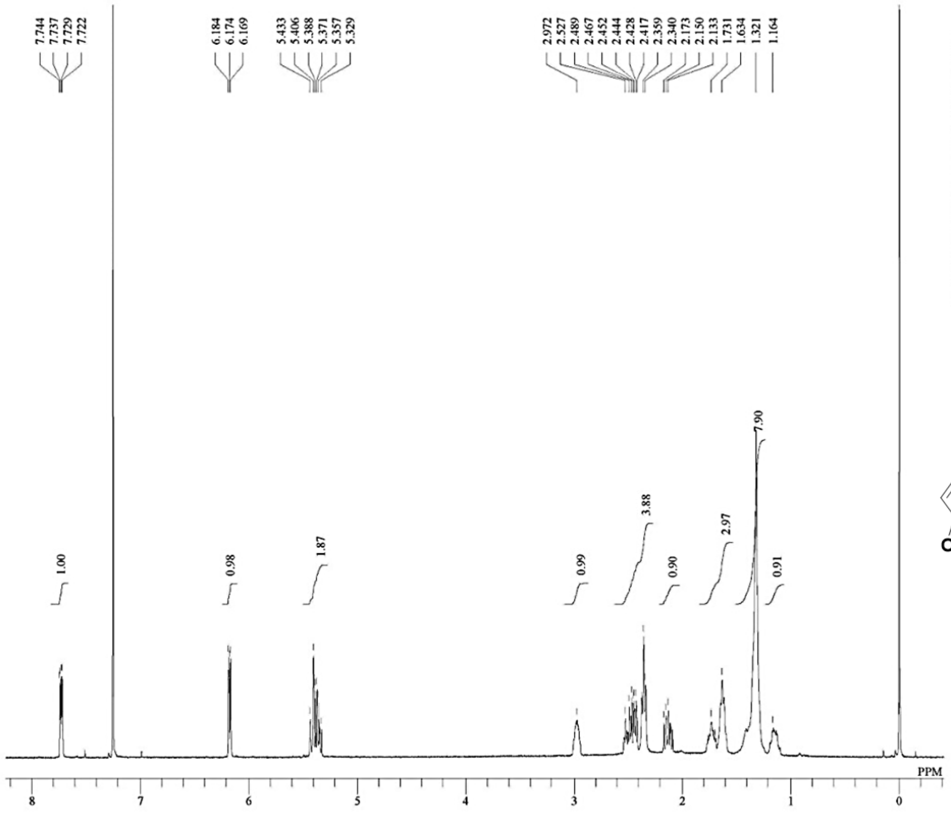


DFILE cis-OPDA-d5 proton.jdf
COMNT single_pulse
DATIM 09-09-2021 19:53:25
OBNUC 1H
EXMOD proton.jxp
OBFRQ 399.78 MHz
OBSET 4.19 KHz
OBFIN 7.29 Hz
POINT 16400
FREQU 7503.00 Hz
SCANS 8
ACQTM 2.1837 sec
PD 5.0000 sec
PW1 2.95 usec
IRNUC 1H
CTEMP 21.9 c
SLVNT CDCL3
EXREF 0.00 ppm
BF 1.00 Hz
RGAIN 64

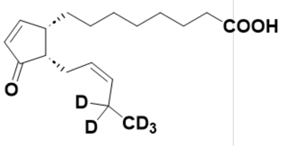


*cis*-OPDA-$d_5$

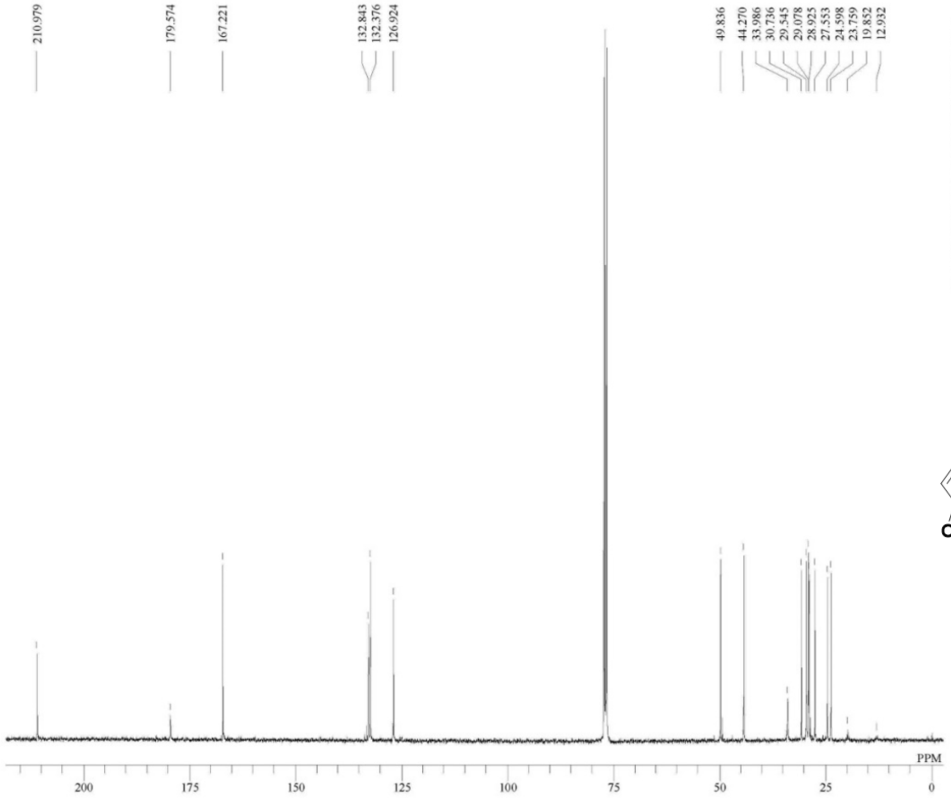


DFILE cis-OPDA-d5 carbon.jdf
COMNT single pulse decoupled gated N[illegible]
DATIM 12-06-2022 06:18:29
OBNUC 13C
EXMOD carbon.jxp
OBFRQ 100.53 MHz
OBSET 5.35 KHz
OBFIN 5.86 Hz
POINT 32780
FREQU 31407.04 Hz
SCANS 33722
ACQTM 1.0433 sec
PD 2.0000 sec
PW1 3.53 usec
IRNUC 1H
CTEMP 21.2 c
SLVNT CDCL3
EXREF 77.00 ppm
BF 1.00 Hz
RGAIN 50

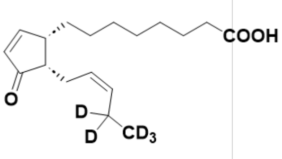


*cis*-OPDA-$d_5$

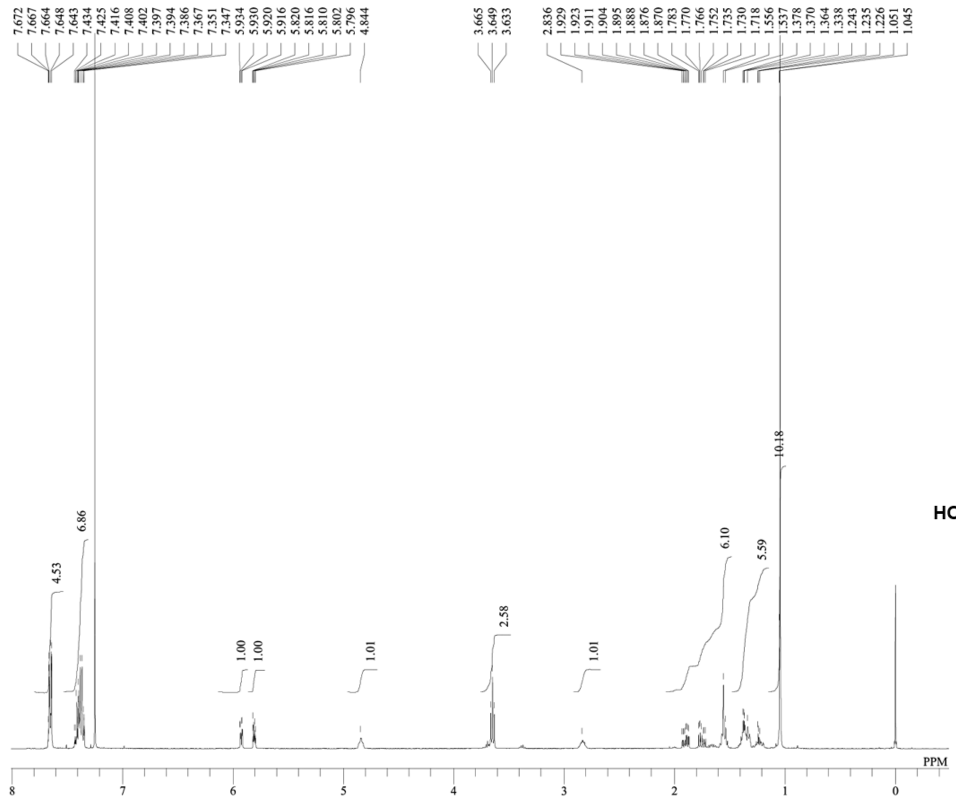


DFILE S10proton.als
COMNT single_pulse
DATIM 29-08-2022 14:11:40
OBNUC 1H
EXMOD proton.jxp
OBFRQ 399.78 MHz
OBSET 4.19 KHz
OBFIN 7.29 Hz
POINT 13120
FREQU 6002.40 Hz
SCANS 8
ACQTM 2.1837 sec
PD 5.0000 sec
PW1 3.40 usec
IRNUC 1H
CTEMP 21.3 c
SLVNT CDCL3
EXREF 0.00 ppm
BF 1.00 Hz
RGAIN 62

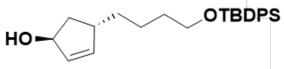


**S5**

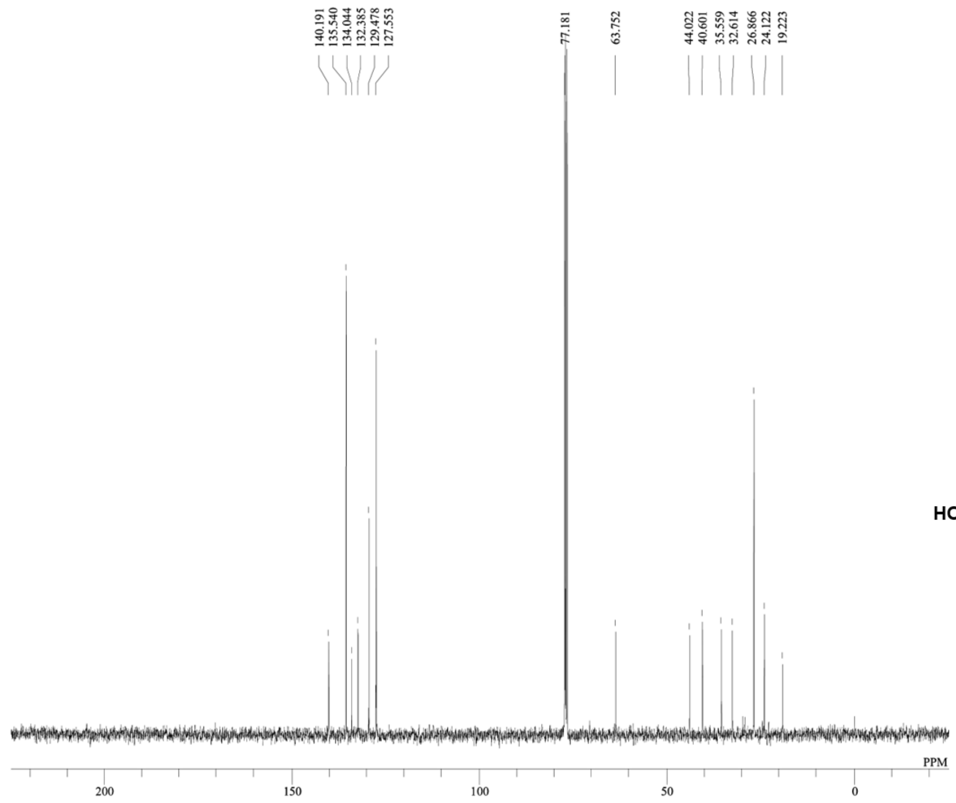


DFILE S10carbon.als
COMNT single pulse decoupled gated N
DATIM 29-08-2022 15:44:04
OBNUC 13C
EXMOD carbon.jxp
OBFRQ 100.53 MHz
OBSET 5.35 KHz
OBFIN 5.86 Hz
POINT 26224
FREQU 25125.63 Hz
SCANS 304
ACQTM 1.0433 sec
PD 2.0000 sec
PW1 3.53 usec
IRNUC 1H
CTEMP 21.3 c
SLVNT CDCL3
EXREF 77.00 ppm
BF 1.00 Hz
RGAIN 50

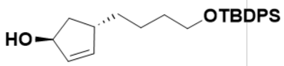


**S5**

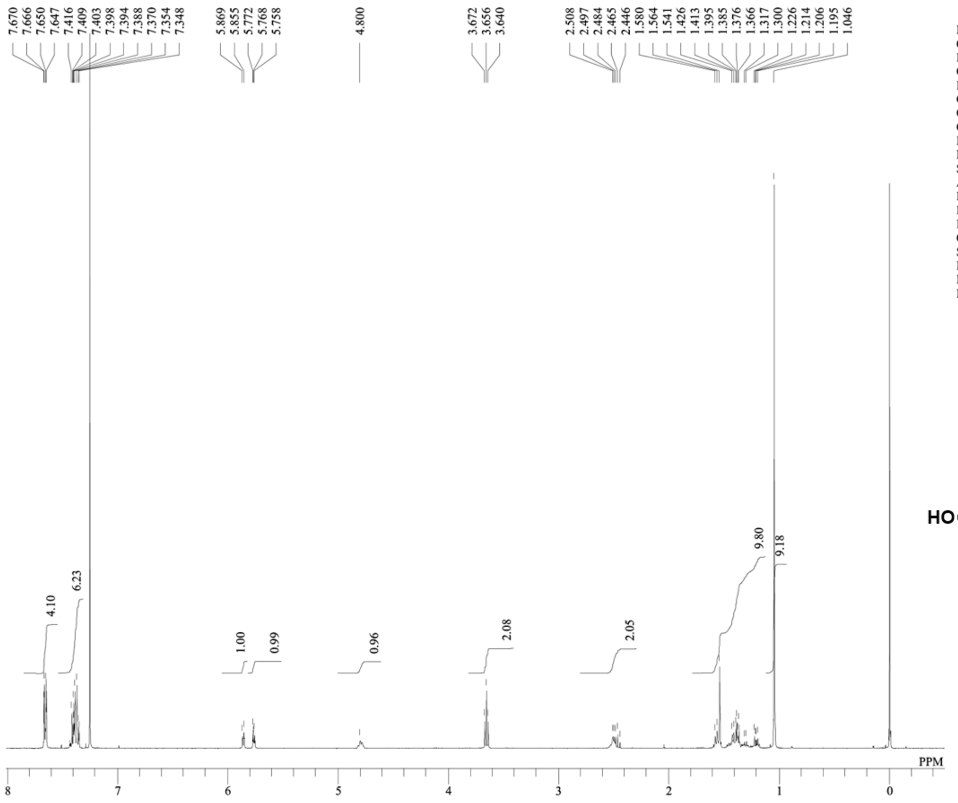


| | |
|---|---|
| DFILE | S11proton.als |
| COMNT | single_pulse |
| DATIM | 14-09-2022 15:47:18 |
| OBNUC | 1H |
| EXMOD | proton.jxp |
| OBFRQ | 399.78 MHz |
| OBSET | 4.19 KHz |
| OBFIN | 7.29 Hz |
| POINT | 13120 |
| FREQU | 6002.40 Hz |
| SCANS | 8 |
| ACQTM | 2.1837 sec |
| PD | 5.0000 sec |
| PW1 | 3.40 usec |
| IRNUC | 1H |
| CTEMP | 21.5 c |
| SLVNT | CDCL3 |
| EXREF | 0.00 ppm |
| BF | 1.00 Hz |
| RGAIN | 76 |

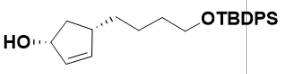


**S6**

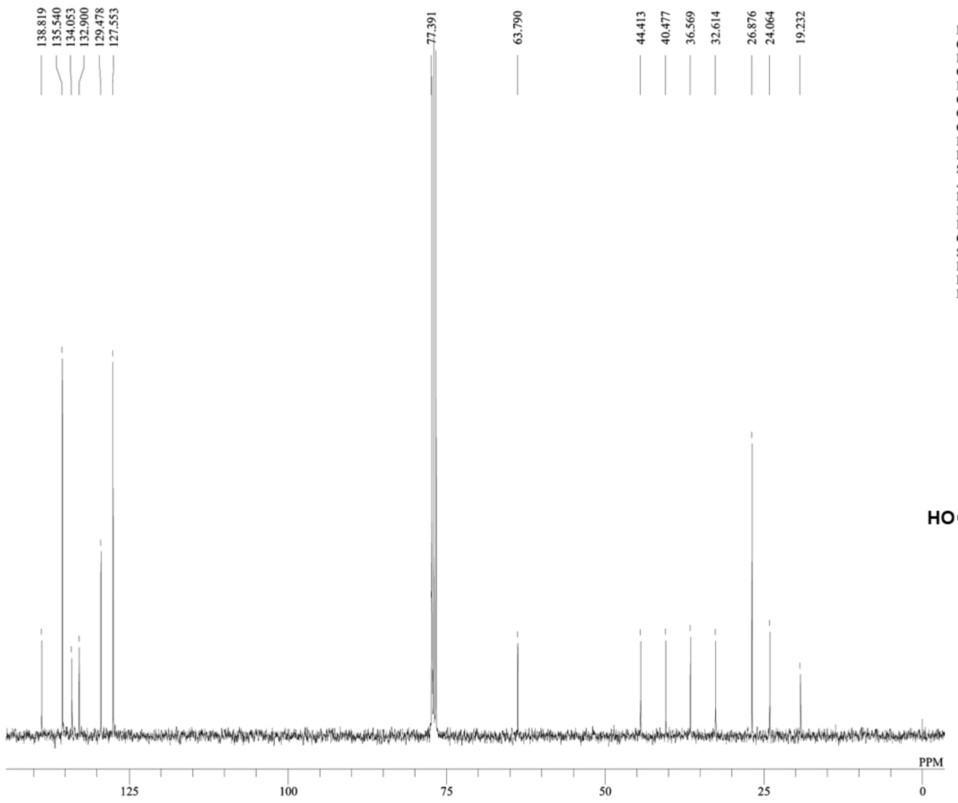


| | |
|---|---|
| DFILE | S11carbon.als |
| COMNT | single pulse decoupled gated N |
| DATIM | 14-09-2022 16:28:39 |
| OBNUC | 13C |
| EXMOD | carbon.jxp |
| OBFRQ | 100.53 MHz |
| OBSET | 5.35 KHz |
| OBFIN | 5.86 Hz |
| POINT | 26224 |
| FREQU | 25125.63 Hz |
| SCANS | 300 |
| ACQTM | 1.0433 sec |
| PD | 2.0000 sec |
| PW1 | 3.53 usec |
| IRNUC | 1H |
| CTEMP | 21.5 c |
| SLVNT | CDCL3 |
| EXREF | 77.00 ppm |
| BF | 1.00 Hz |
| RGAIN | 50 |

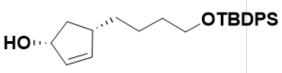


**S6**

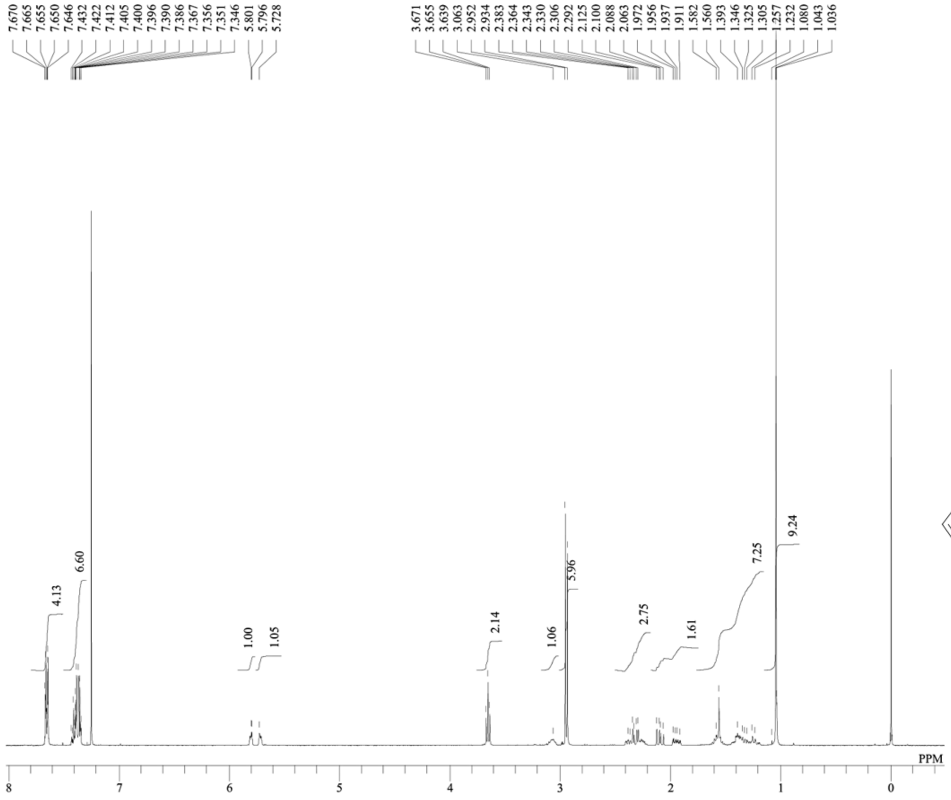
7.670
7.665
7.655
7.650
7.646
7.432
7.422
7.412
7.405
7.400
7.396
7.390
7.386
7.367
7.356
7.351
7.346
5.801
5.796
5.728
3.671
3.655
3.639
3.063
2.952
2.934
2.383
2.364
2.343
2.330
2.306
2.292
2.125
2.100
2.088
2.063
1.972
1.956
1.937
1.911
1.582
1.560
1.393
1.346
1.325
1.305
1.257
1.232
1.080
1.043
1.036
4.13
6.60
1.00
1.05
2.14
1.06
5.96
2.75
1.61
7.25
9.24
PPM
8
7
6
5
4
3
2
1
0
DFILE S12proton.als
COMNT single_pulse
DATIM 20-09-2022 15:30:52
OBNUC 1H
EXMOD proton.jxp
OBFRQ 399.78 MHz
OBSET 4.19 KHz
OBFIN 7.29 Hz
POINT 13120
FREQU 6002.40 Hz
SCANS 8
ACQTM 2.1837 sec
PD 5.0000 sec
PW1 3.40 usec
IRNUC 1H
CTEMP 21.7 c
SLVNT CDCL3
EXREF 0.00 ppm
BF 1.00 Hz
RGAIN 70

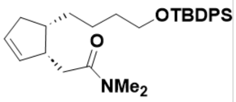
OTBDPS
O
NMe2

S7

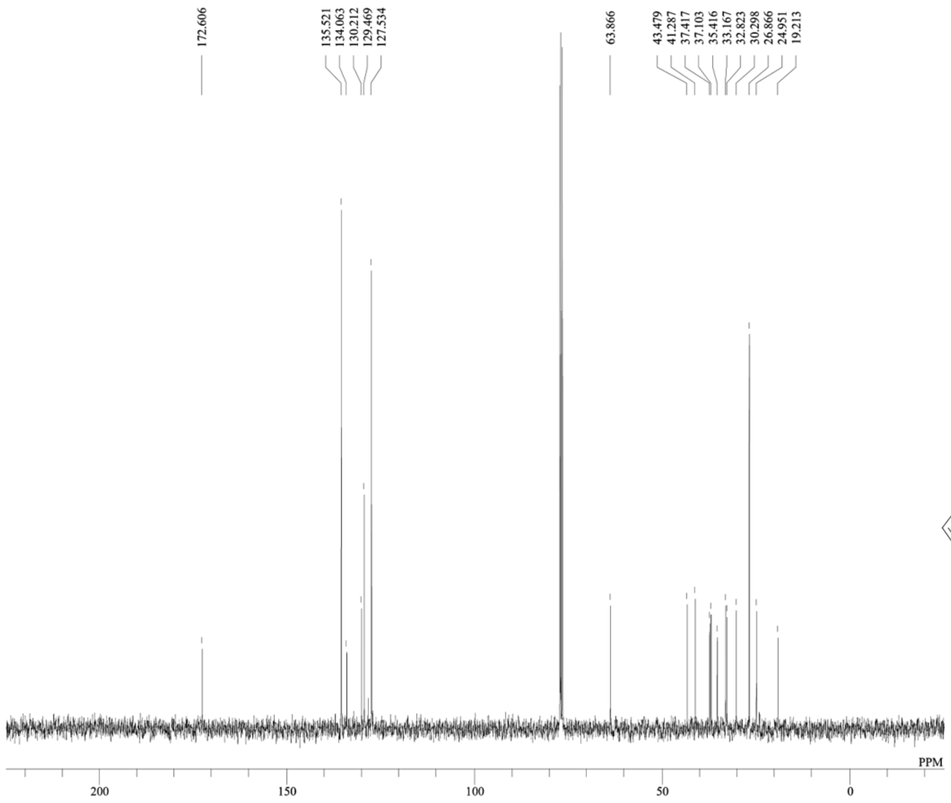
172.606
135.521
134.063
130.212
129.469
127.534
63.866
43.479
41.287
37.417
37.103
35.416
33.167
32.823
30.298
26.866
24.951
19.213
PPM
200
150
100
50
0
DFILE S12carbon.als
COMNT single pulse decoupled gated N
DATIM 20-09-2022 15:41:04
OBNUC 13C
EXMOD carbon.jxp
OBFRQ 100.53 MHz
OBSET 5.35 KHz
OBFIN 5.86 Hz
POINT 26224
FREQU 25125.63 Hz
SCANS 133
ACQTM 1.0433 sec
PD 2.0000 sec
PW1 3.53 usec
IRNUC 1H
CTEMP 21.6 c
SLVNT CDCL3
EXREF 77.00 ppm
BF 1.00 Hz
RGAIN 50

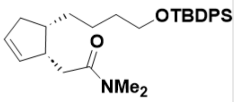
OTBDPS
O
NMe2

S7

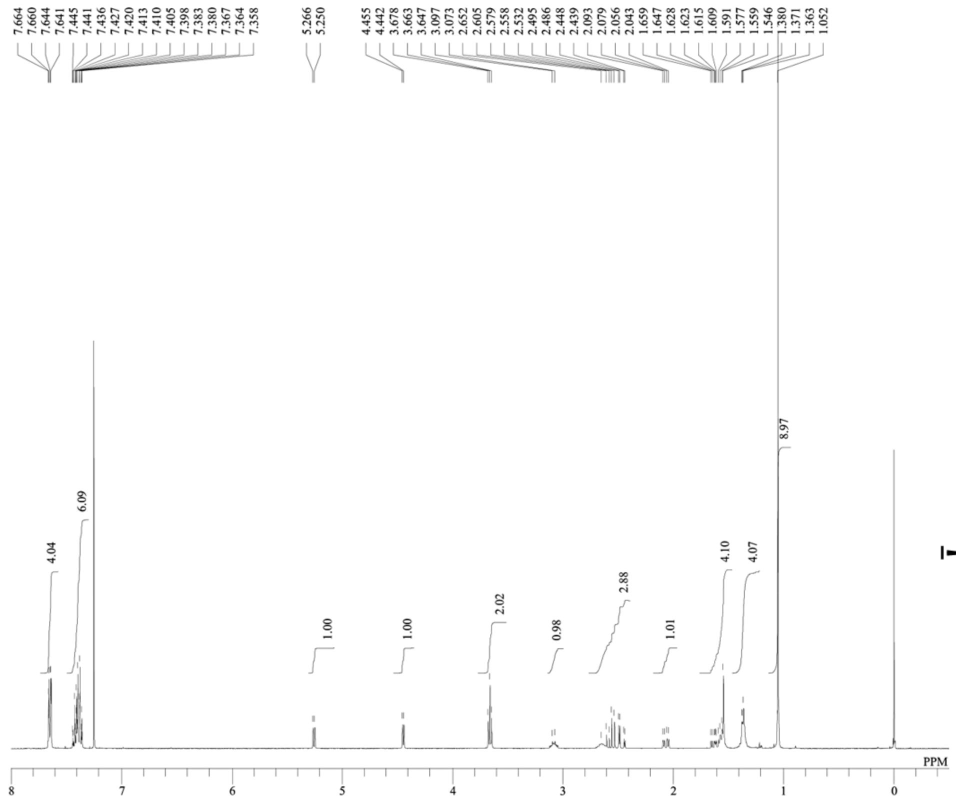


DFILE S13proton.als
COMNT single_pulse
DATIM 22-09-2022 15:16:55
OBNUC 1H
EXMOD proton.jxp
OBFRQ 399.78 MHz
OBSET 4.19 KHz
OBFIN 7.29 Hz
POINT 13120
FREQU 6002.40 Hz
SCANS 8
ACQTM 2.1837 sec
PD 5.0000 sec
PW1 3.40 usec
IRNUC 1H
CTEMP 21.3 c
SLVNT CDCL3
EXREF 0.00 ppm
BF 1.00 Hz
RGAIN 70

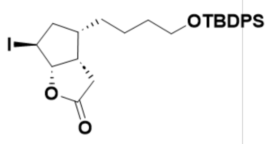


S8

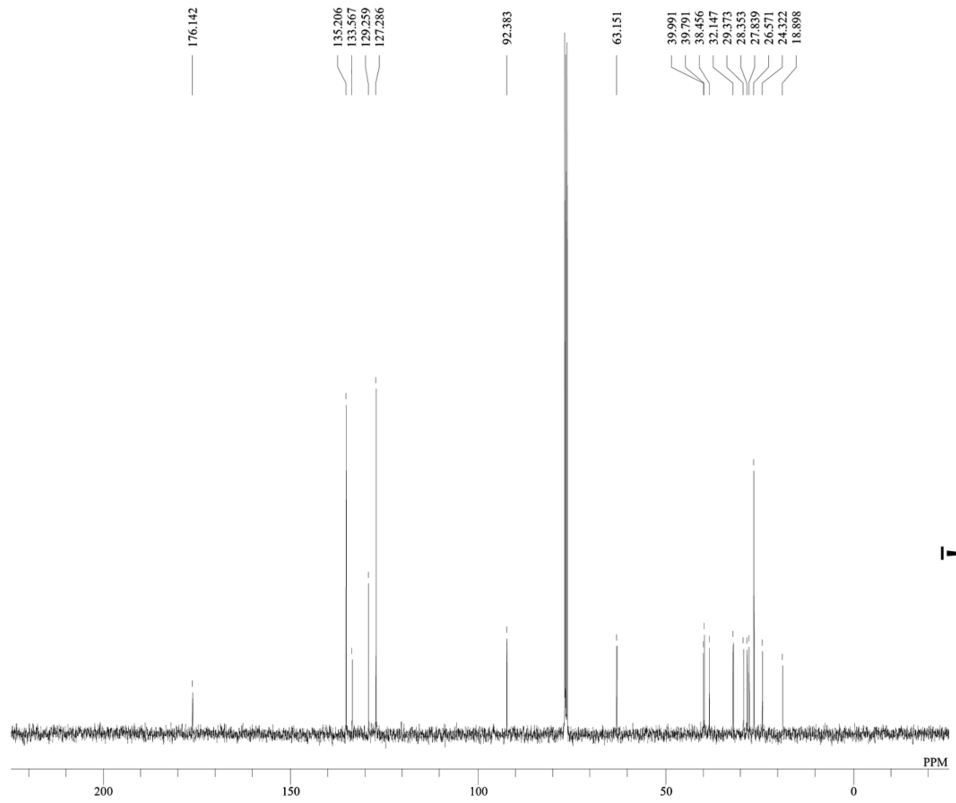


DFILE S13carbon.als
COMNT single pulse decoupled gated N
DATIM 22-09-2022 14:53:28
OBNUC 13C
EXMOD carbon.jxp
OBFRQ 100.53 MHz
OBSET 5.35 KHz
OBFIN 5.86 Hz
POINT 26224
FREQU 25125.63 Hz
SCANS 265
ACQTM 1.0433 sec
PD 2.0000 sec
PW1 3.53 usec
IRNUC 1H
CTEMP 21.4 c
SLVNT CDCL3
EXREF 77.00 ppm
BF 1.00 Hz
RGAIN 50

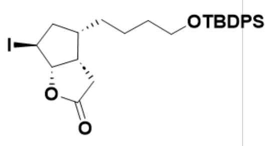


S8

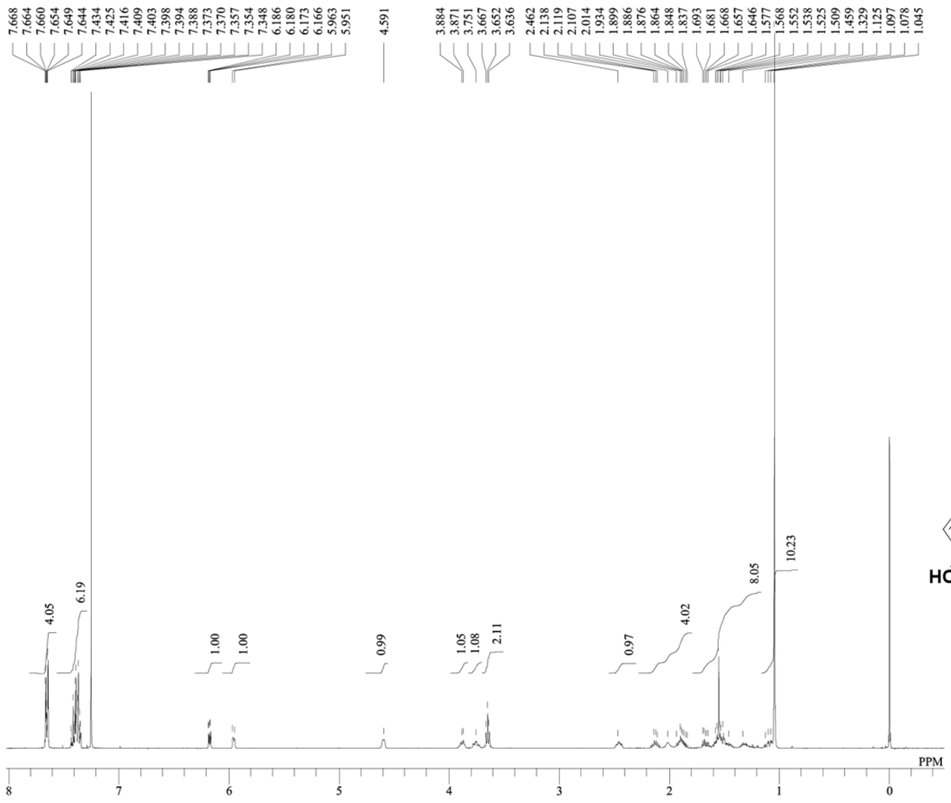


DFILE S14proton.als
COMNT single_pulse
DATIM 29-09-2022 11:07:20
OBNUC 1H
EXMOD proton.jxp
OBFRQ 399.78 MHz
OBSET 4.19 KHz
OBFIN 7.29 Hz
POINT 13120
FREQU 6002.40 Hz
SCANS 8
ACQTM 2.1837 sec
PD 5.0000 sec
PW1 3.40 usec
IRNUC 1H
CTEMP 21.4 c
SLVNT CDCL3
EXREF 0.00 ppm
BF 1.00 Hz
RGAIN 72

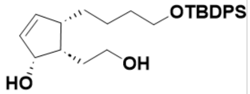


**S9**

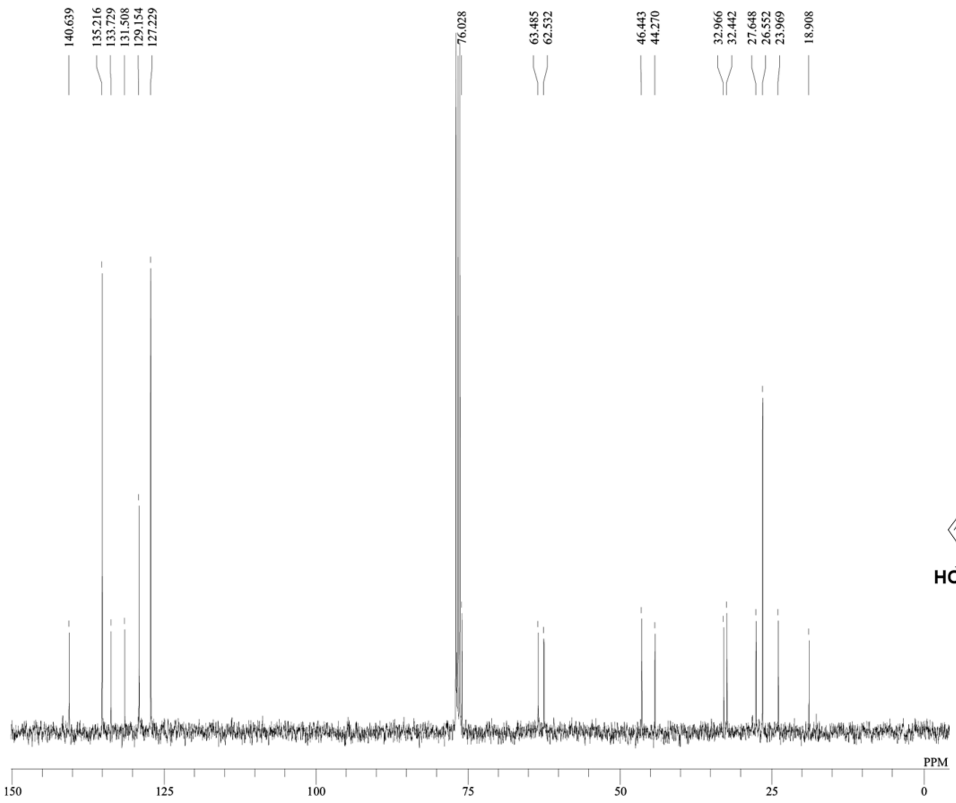


DFILE S14carbon.als
COMNT single pulse decoupled gated N
DATIM 29-09-2022 11:14:29
OBNUC 13C
EXMOD carbon.jxp
OBFRQ 100.53 MHz
OBSET 5.35 KHz
OBFIN 5.86 Hz
POINT 26224
FREQU 25125.63 Hz
SCANS 144
ACQTM 1.0433 sec
PD 2.0000 sec
PW1 3.53 usec
IRNUC 1H
CTEMP 21.4 c
SLVNT CDCL3
EXREF 77.00 ppm
BF 1.00 Hz
RGAIN 50

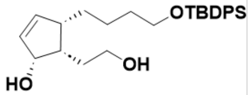


**S9**

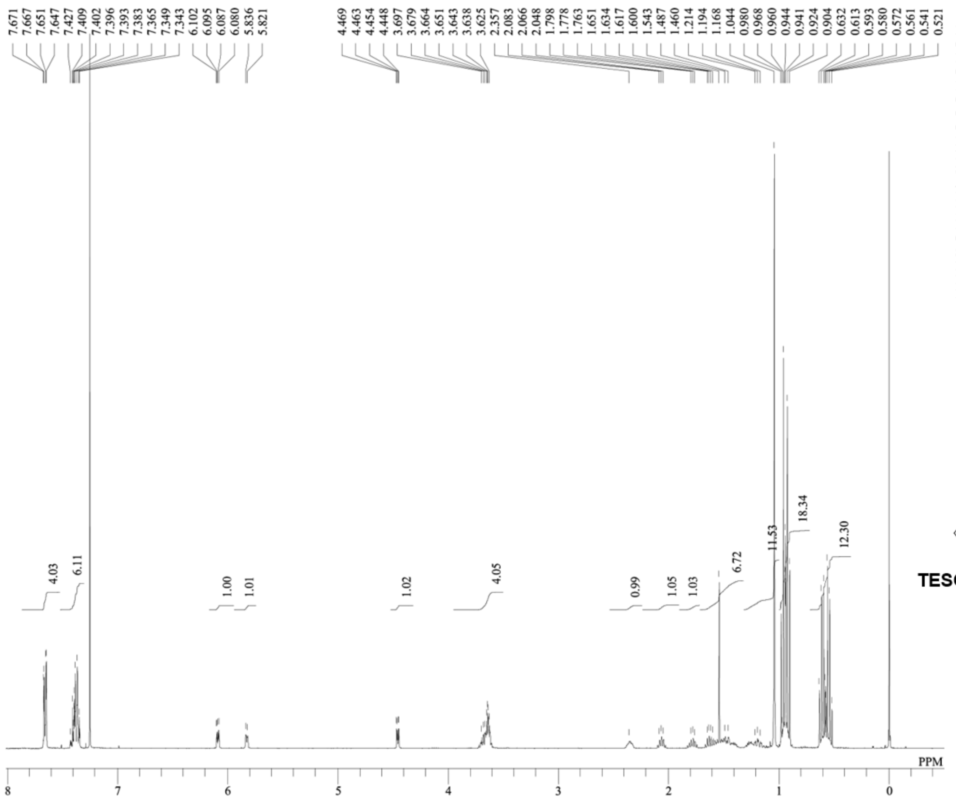


DFILE S15proton.als
COMNT single_pulse
DATIM 30-09-2022 19:23:00
OBNUC 1H
EXMOD proton.jxp
OBFRQ 399.78 MHz
OBSET 4.19 KHz
OBFIN 7.29 Hz
POINT 13120
FREQU 6002.40 Hz
SCANS 8
ACQTM 2.1837 sec
PD 5.0000 sec
PW1 3.40 usec
IRNUC 1H
CTEMP 21.2 c
SLVNT CDCL3
EXREF 0.00 ppm
BF 1.00 Hz
RGAIN 70

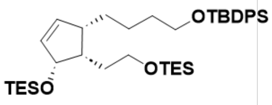


**S10**

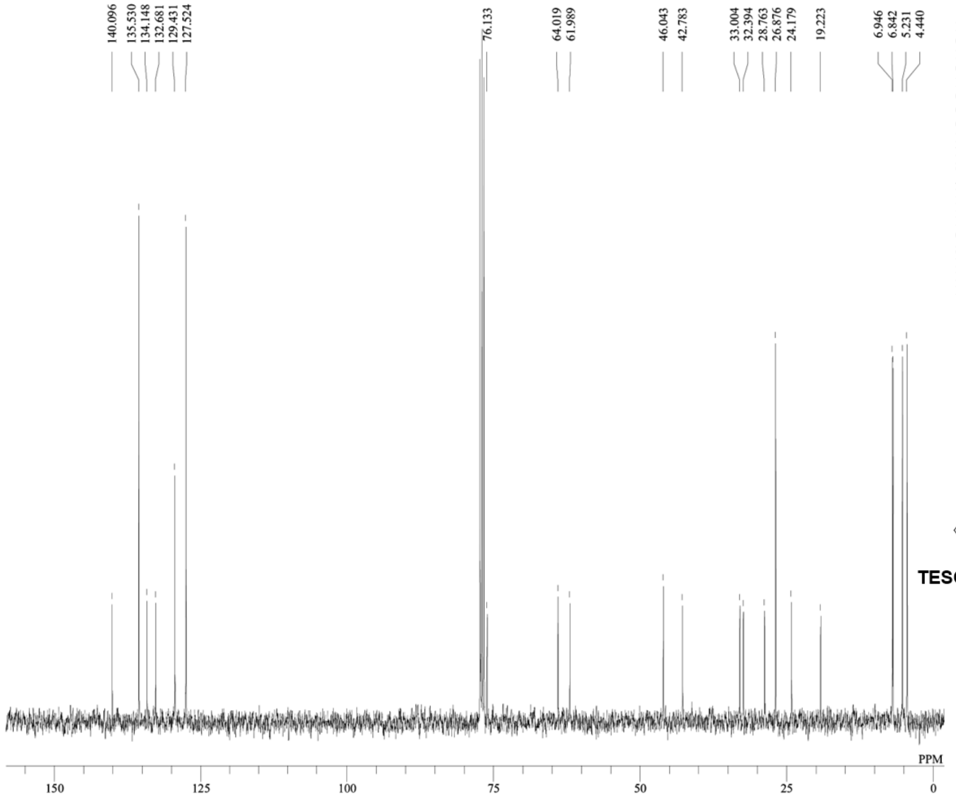


DFILE S15carbon.als
COMNT single pulse decoupled gated N
DATIM 30-09-2022 19:46:55
OBNUC 13C
EXMOD carbon.jxp
OBFRQ 100.53 MHz
OBSET 5.35 KHz
OBFIN 5.86 Hz
POINT 26224
FREQU 25125.63 Hz
SCANS 88
ACQTM 1.0433 sec
PD 2.0000 sec
PW1 3.53 usec
IRNUC 1H
CTEMP 21.2 c
SLVNT CDCL3
EXREF 77.00 ppm
BF 1.00 Hz
RGAIN 50

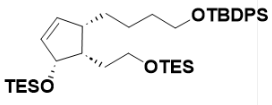


**S10**

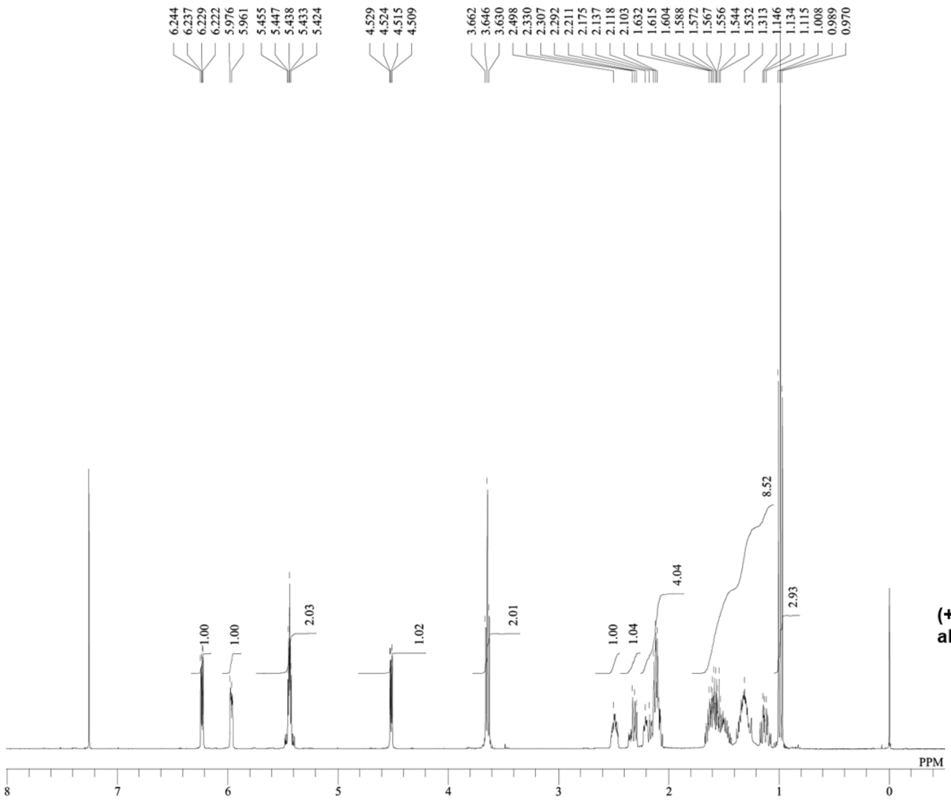


DFILE (+)-8-(R)-hydroxy-phytodienoy
COMNT single_pulse
DATIM 08-11-2022 20:34:44
OBNUC 1H
EXMOD proton.jxp
OBFRQ 399.78 MHz
OBSET 4.19 KHz
OBFIN 7.29 Hz
POINT 13120
FREQU 6002.40 Hz
SCANS 8
ACQTM 2.1837 sec
PD 5.0000 sec
PW1 3.40 usec
IRNUC 1H
CTEMP 20.9 c
SLVNT CDCL3
EXREF 0.00 ppm
BF 1.00 Hz
RGAIN 64

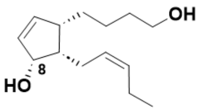


**(+)-8-(*R*)-hydroxy-phytodienoyl alcohol**

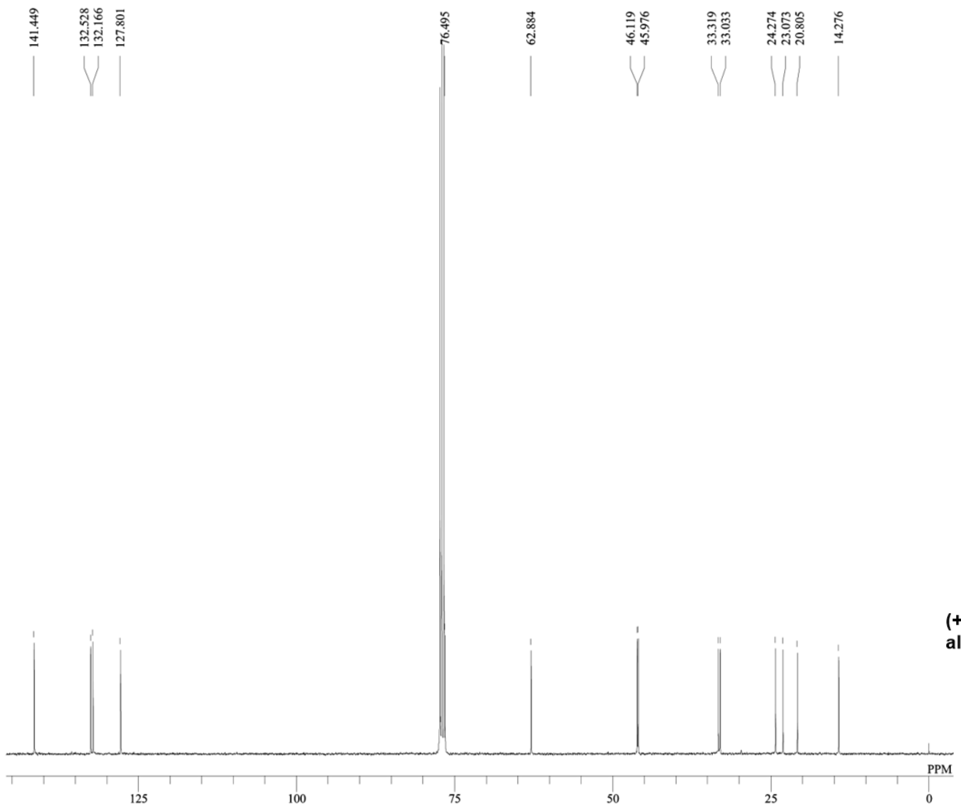


DFILE (+)-8-(R)-hydroxy-phytodienoy
COMNT single pulse decoupled gated N
DATIM 09-11-2022 00:30:58
OBNUC 13C
EXMOD carbon.jxp
OBFRQ 100.53 MHz
OBSET 5.35 KHz
OBFIN 5.86 Hz
POINT 26224
FREQU 25125.63 Hz
SCANS 10402
ACQTM 1.0433 sec
PD 2.0000 sec
PW1 3.53 usec
IRNUC 1H
CTEMP 20.9 c
SLVNT CDCL3
EXREF 77.00 ppm
BF 1.00 Hz
RGAIN 50

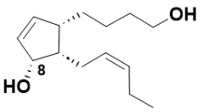


**(+)-8-(*R*)-hydroxy-phytodienoyl alcohol**

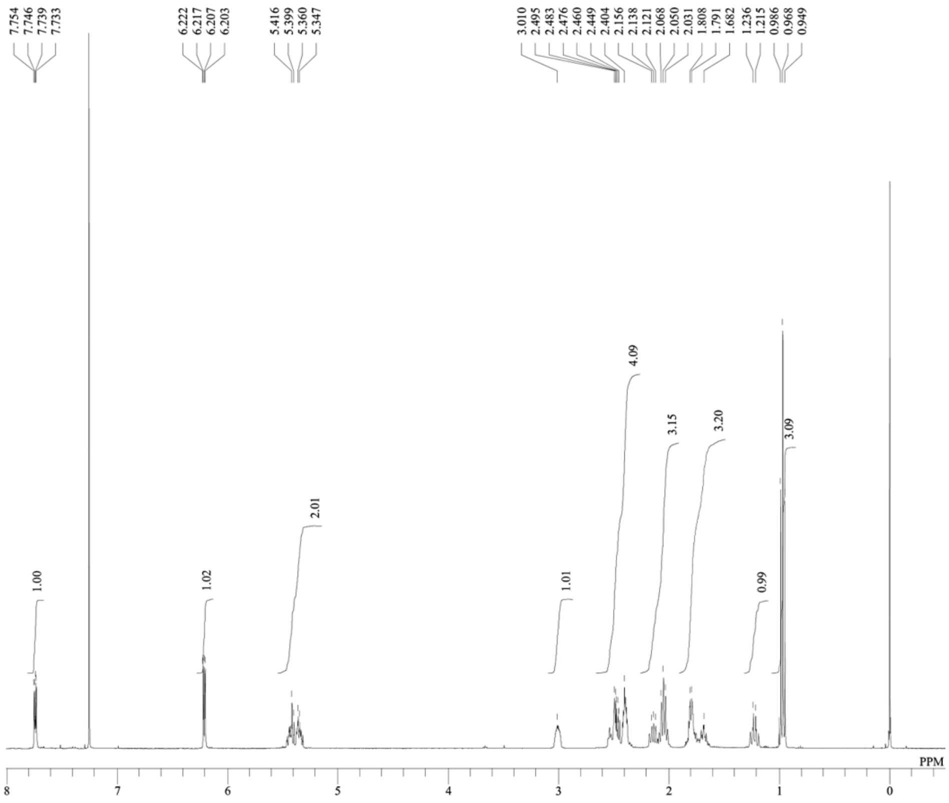


DFILE tn-cis-OPDA_proton.als
COMNT single_pulse
DATIM 27-10-2022 14:04:28
OBNUC 1H
EXMOD proton.jxp
OBFRQ 399.78 MHz
OBSET 4.19 KHz
OBFIN 7.29 Hz
POINT 13120
FREQU 6002.40 Hz
SCANS 52
ACQTM 2.1837 sec
PD 5.0000 sec
PW1 3.40 usec
IRNUC 1H
CTEMP 20.8 c
SLVNT CDCL3
EXREF 0.00 ppm
BF 1.00 Hz
RGAIN 72

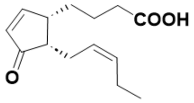


**tn-*cis*-OPDA**

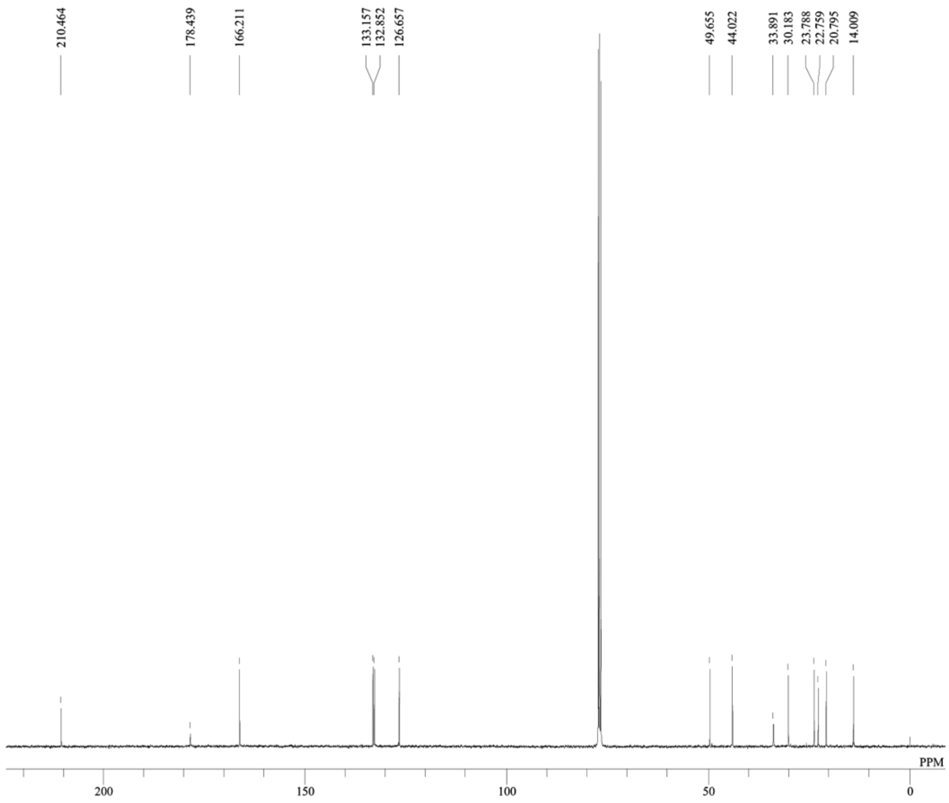


DFILE tn-cis-OPDA_carbon.als
COMNT single pulse decoupled gated N
DATIM 28-10-2022 23:46:23
OBNUC 13C
EXMOD carbon.jxp
OBFRQ 100.53 MHz
OBSET 5.35 KHz
OBFIN 5.86 Hz
POINT 26224
FREQU 25125.63 Hz
SCANS 11164
ACQTM 1.0433 sec
PD 2.0000 sec
PW1 3.53 usec
IRNUC 1H
CTEMP 20.6 c
SLVNT CDCL3
EXREF 77.00 ppm
BF 1.00 Hz
RGAIN 50

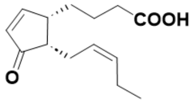


**tn-*cis*-OPDA**

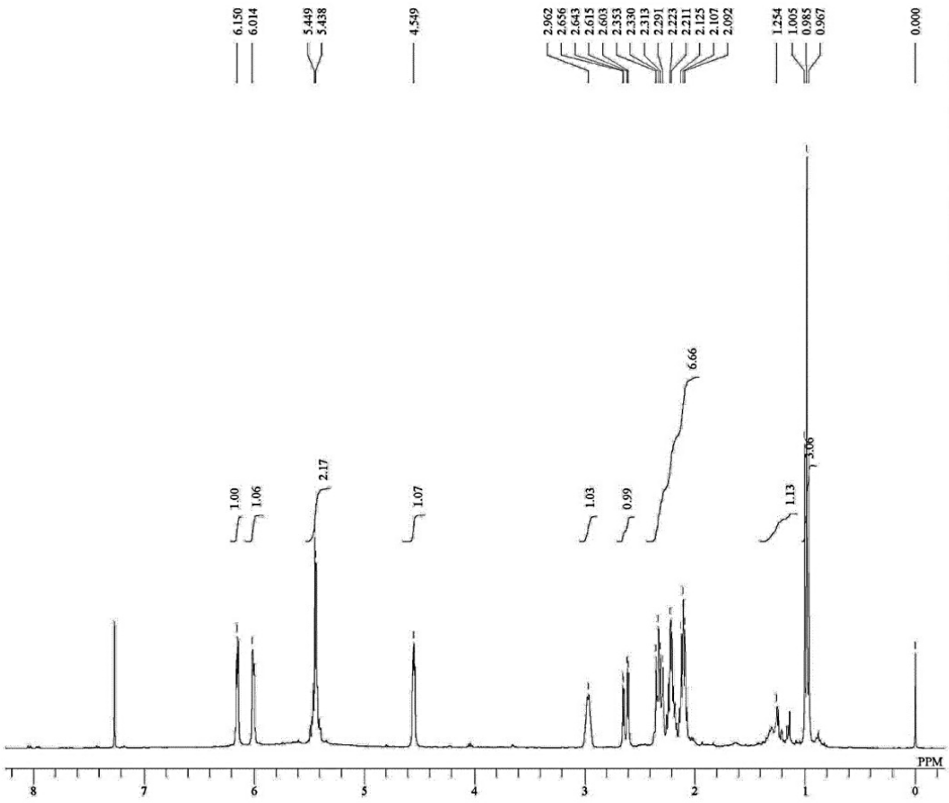


DFILE 09163p_Proton-5-1.jdf
COMNT single_pulse
DATIM 04-06-2022 00:34:06
OBNUC 1H
EXMOD proton.jxp
OBFRQ 399.78 MHz
OBSET 4.19 KHz
OBFIN 7.29 Hz
POINT 16384
FREQU 7503.00 Hz
SCANS 8
ACQTM 2.1837 sec
PD 5.0000 sec
PW1 3.40 usec
IRNUC 1H
CTEMP 21.4 c
SLVNT CDCL3
EXREF 0.00 ppm
BF 1.00 Hz
RGAIN 60

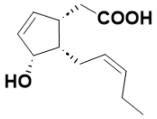


**S12**

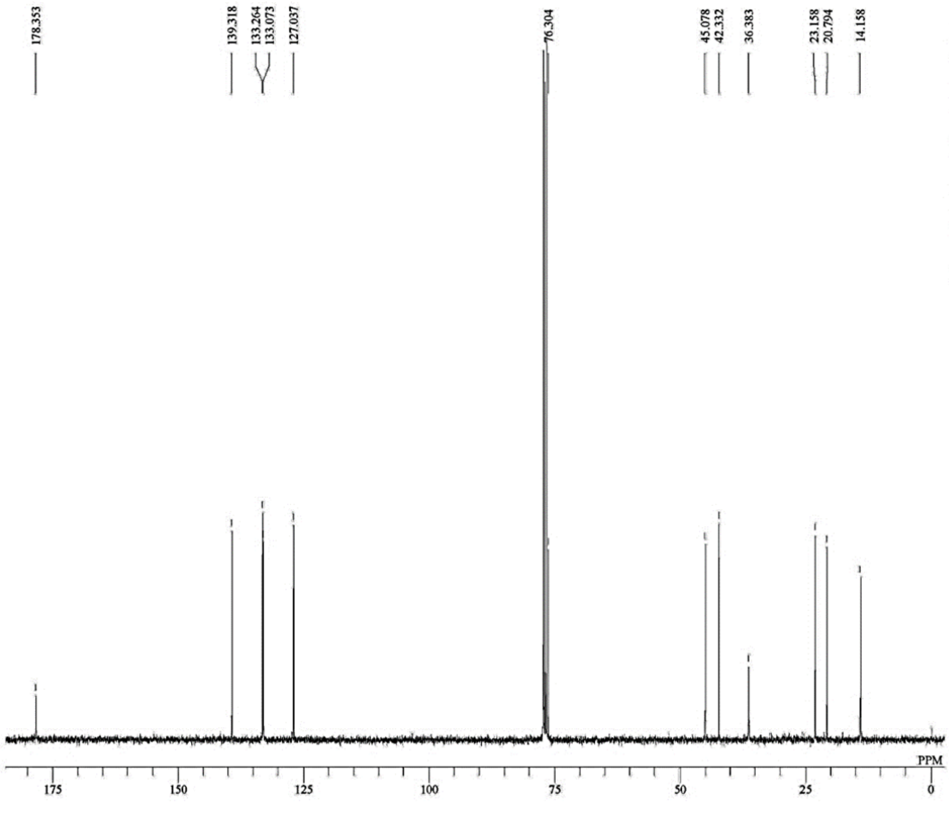


DFILE 09163p_Carbon-1-1.jdf
COMNT single pulse decoupled gated N
DATIM 04-06-2022 00:35:19
OBNUC 13C
EXMOD carbon.jxp
OBFRQ 100.53 MHz
OBSET 5.35 KHz
OBFIN 5.86 Hz
POINT 32767
FREQU 31407.04 Hz
SCANS 1024
ACQTM 1.0433 sec
PD 2.0000 sec
PW1 3.53 usec
IRNUC 1H
CTEMP 21.2 c
SLVNT CDCL3
EXREF 77.00 ppm
BF 1.00 Hz
RGAIN 50

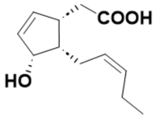


**S12**

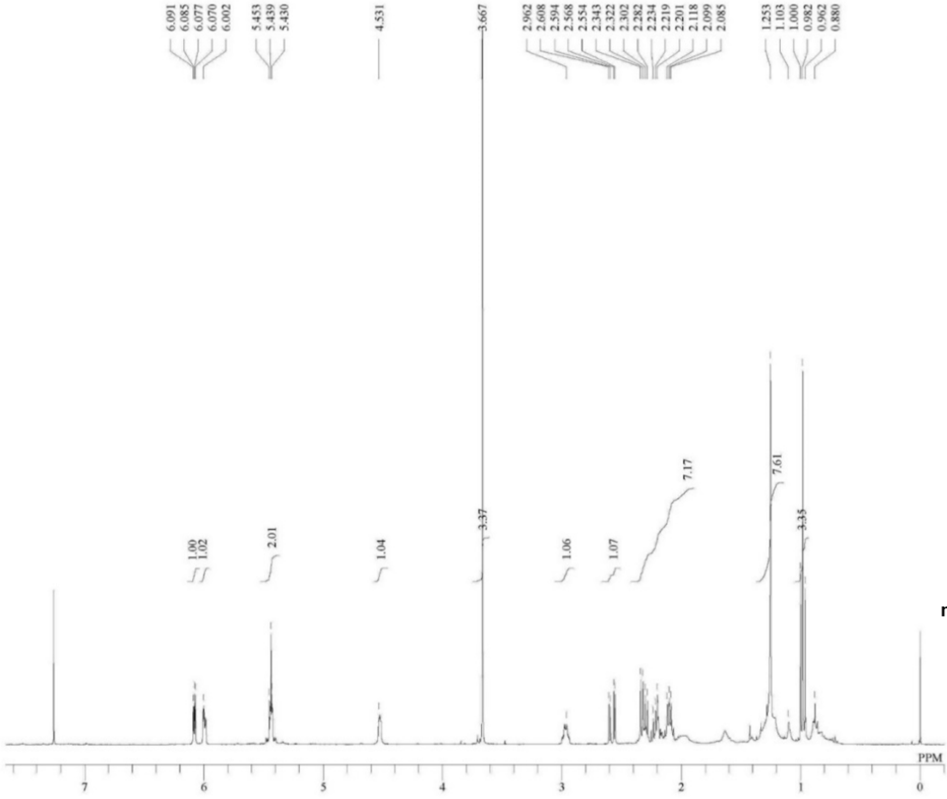


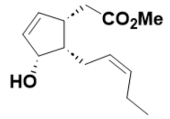


**methyl 4,5-ddh-6-*epi*-cucurbate**

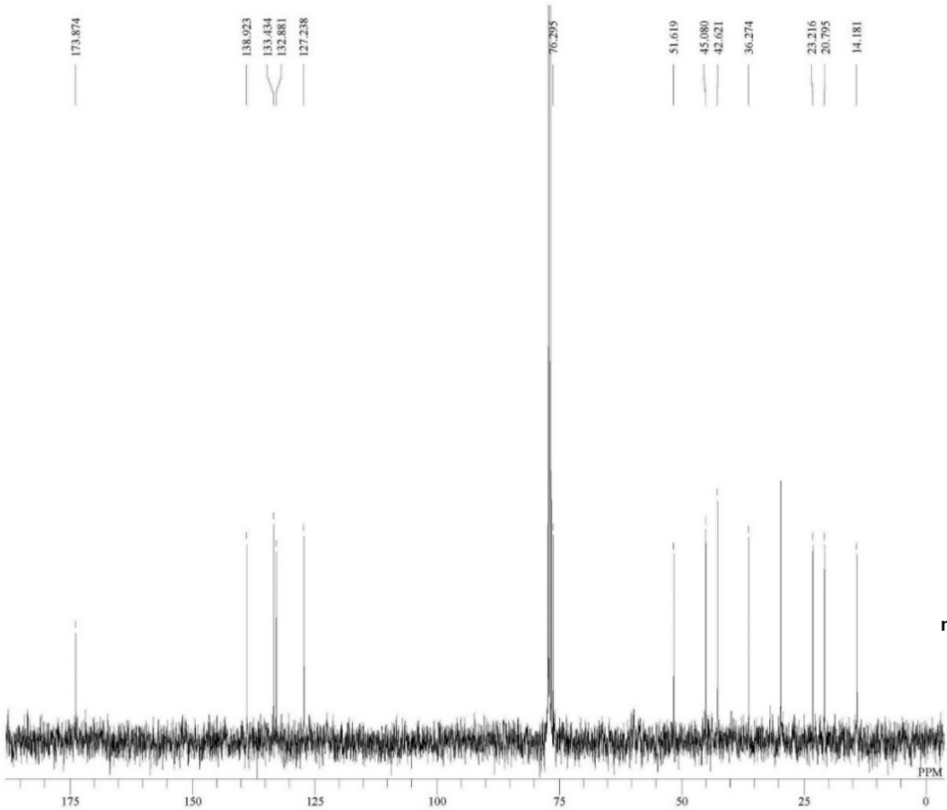


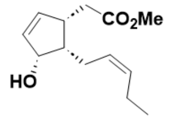


**methyl 4,5-ddh-6-*epi*-cucurbate**

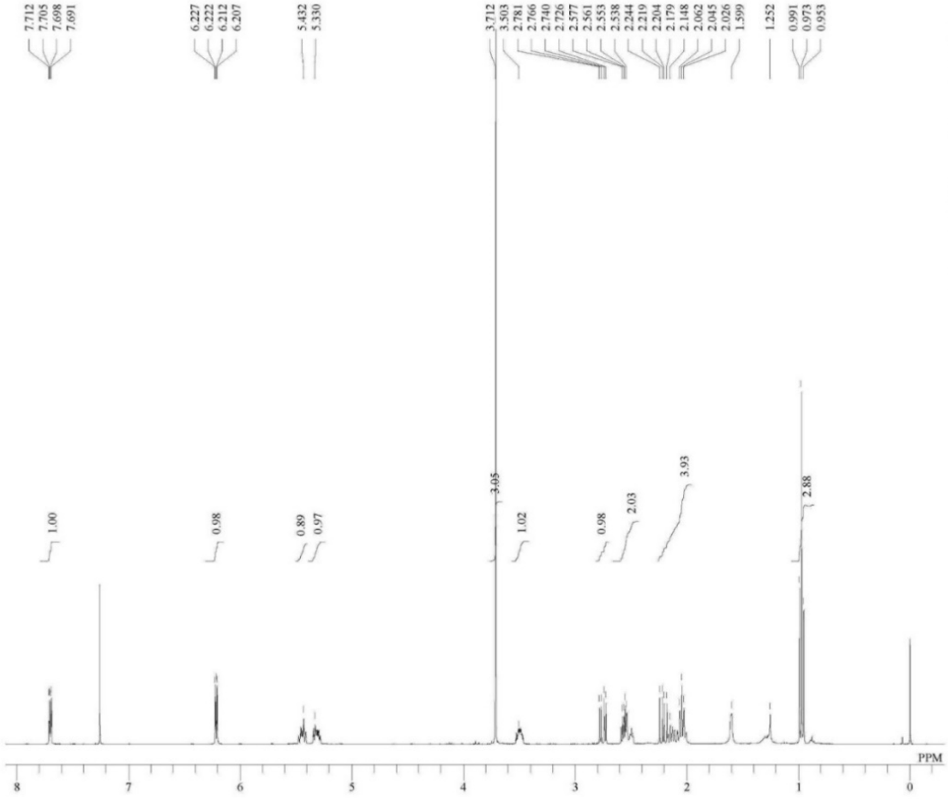


DFILE 09179p-1.jdf
COMNT single_pulse
DATIM 03-07-2022 22:47:56
OBNUC 1H
EXMOD proton.jxp
OBFRQ 399.78 MHz
OBSET 4.19 KHz
OBFIN 7.29 Hz
POINT 16400
FREQU 7503.00 Hz
SCANS 8
ACQTM 2.1837 sec
PD 5.0000 sec
PW1 3.40 usec
IRNUC 1H
CTEMP 21.3 c
SLVNT CDCL3
EXREF 0.00 ppm
BF 1.00 Hz
RGAIN 70

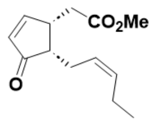


**4,5-ddh-*cis*-MeJA**

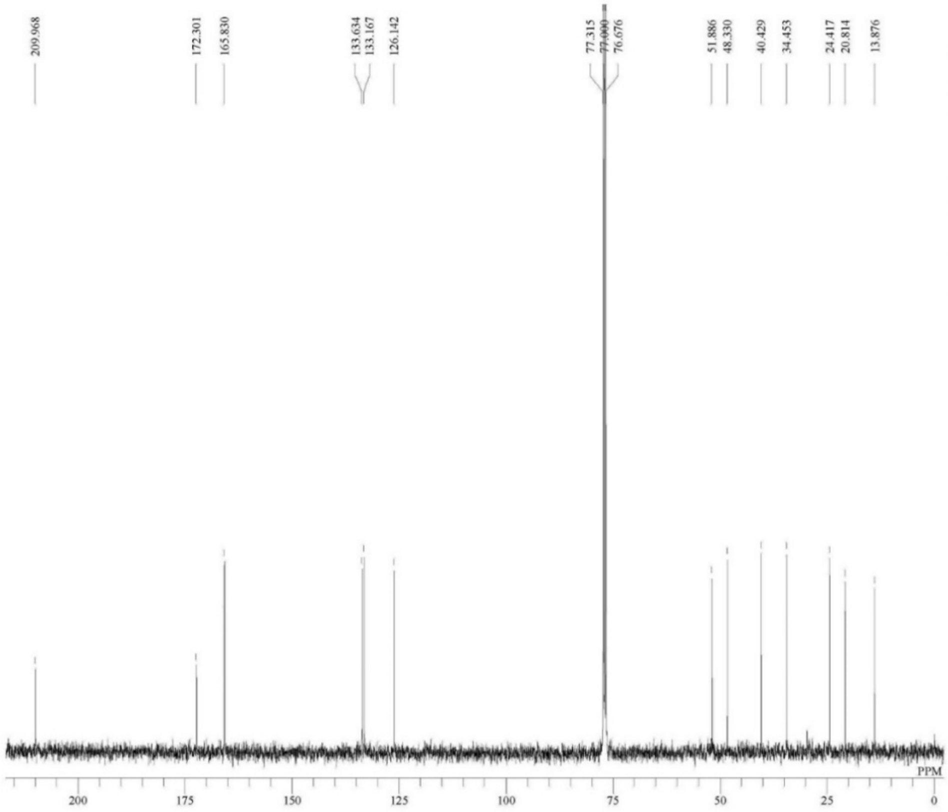


DFILE 09179p_13C.jdf
COMNT single pulse decoupled gated N
DATIM 03-07-2022 23:15:29
OBNUC 13C
EXMOD carbon.jxp
OBFRQ 100.53 MHz
OBSET 5.35 KHz
OBFIN 5.86 Hz
POINT 32780
FREQU 31407.04 Hz
SCANS 1024
ACQTM 1.0433 sec
PD 2.0000 sec
PW1 3.53 usec
IRNUC 1H
CTEMP 21.1 c
SLVNT CDCL3
EXREF 77.00 ppm
BF 1.00 Hz
RGAIN 50

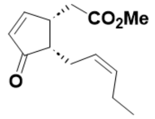


**4,5-ddh-*cis*-MeJA**
